# Sequencing by Expansion (SBX) – a novel, high-throughput single-molecule sequencing technology

**DOI:** 10.1101/2025.02.19.639056

**Authors:** Mark Kokoris, Robert McRuer, Melud Nabavi, Aaron Jacobs, Marc Prindle, Cynthia Cech, Kendall Berg, Taylor Lehmann, Cara Machacek, John Tabone, Jagadeeswaran Chandrasekar, Lacey McGee, Matthew Lopez, Tommy Reid, Cara Williams, Salka Barrett, Alex Lehmann, Michael Kovarik, Robert Busam, Scott Miller, Brent Banasik, Brittany Kesic, Anasha Arryman, Megan Rogers-Peckham, Alan Kimura, Megan LeProwse, Mitchell Wolfin, Svetlana Kritzer, Joanne Leadbetter, Majid Babazadeh, John Chase, Greg Thiessen, William Lint, Drew Goodman, Dylan O’Connell, Nadya Lumanpauw, John Hoffman, Samantha Vellucci, Kendra Collins, Jessica Vellucci, Amy Taylor, Molly Murphy, Michael Lee, Matthew Corning

## Abstract

Remarkable advances in high-throughput sequencing have enabled major biological discoveries and clinical applications, but achieving wider distribution and use depends critically on further improvements in scale and cost reduction. Nanopore sequencing has long held the promise for such progress, but has had limited market penetration. This is because efficient and accurate nanopore sequencing of nucleic acids has been challenged by fundamental signal-to-noise limitations resulting from the poor spatial resolution and molecular distinction of nucleobases. Here, we describe Sequencing by Expansion (SBX), a single-molecule sequencing technology that overcomes these limitations by using a biochemical conversion process to encode the sequence of a target nucleic acid molecule into an Xpandomer, a highly measurable surrogate polymer. Expanding over 50 times longer than the parent DNA templates, Xpandomers are engineered with high signal-to-noise reporter codes to enable facile, high-accuracy nanopore sequencing. We demonstrate the performance of SBX and present the specialized molecular structures, chemistries, enzymes and methods that enable it. The innovative molecular and systems engineering in SBX create a transformative technology to address the needs of existing and emerging sequencing applications.

---

Dear Readers,

Nucleic acid sequencing is now well established as a core readout for all molecular, cell and tissue biology, with applications from basic discovery to clinical practice. With every increase in scale or drop in cost, the applications and impact of sequencing have increased, as evidenced in the last few years by the rise of population scale efforts like UKBB, scRNA-seq based screens (Perturb-Seq) or the Human Cell Atlas. Despite great advances, high quality sequencing at scale, in terms of both speed and cost, remains a challenge. While costs decreased and scale increased dramatically, it is still an expensive step in every lab experiment and it still limits the routine application of sequencing in molecular diagnostics.

Nanopore sequencing originally offered a promising direction, but the signal-to-noise challenges of direct DNA measurement limited application of the technology. The Sequencing by Expansion (SBX) technology was invented in 2007 at Stratos Genomics, Inc. (now Roche) to address this challenge. Here, we report a key breakthrough step: Sequencing by Expansion on a single pore system. This submission will be the first publication of SBX and will describe molecular designs, syntheses and processes along with sequencing results that were generated prior to the Roche acquisition in 2020. In the coming months, we intend to publish current results for SBX, combined with our high-throughput 8M sensor array, to demonstrate the technology’s performance for key application areas like somatic and germline whole genome sequencing as well as read-intensive methods like perturb seq (scRNA) and spatial sequencing, to name a few. We consider the advancements described in this manuscript to be an important demonstration of what is possible with innovative molecular, protein and systems engineering.

This technical achievement was made possible because of the development of the following key innovations:

– XNTPs, nucleoside triphosphates modified with structures engineered to control translocation, expand, provide well-resolved nanopore based reporter codes and enable template-dependent incorporation with a modified polymerase (Xp Synthase).
– PEMs, a class of critical cofactors that enable polymerase extension of XNTPs beyond 10 bases by stabilizing the elongating heteroduplex structure.
– Xp Synthases, highly mutated DNA polymerases that processively and accurately incorporate XNTPs to synthesize Xpandomers (Xp).
– Reaction formulations and processes, optimized for accurate, template-dependent synthesis of Xp using a solid-phase workflow.
– Xp end structures and sequencing conditions, engineered to efficiently and accurately thread Xp molecules though a nanopore.

For a quick overview of SBX, the URL for a concept animation is included on the following webpage: https://sequencing.roche.com/global/en/article-listing/nanopore-sequencing.html

Thank you,

Mark Kokoris (corresponding author –)

Head of SBX Technology, Roche Sequencing Solutions, Seattle WA, USA

Co-founder and CEO of Stratos Genomics, Inc. (2007-2020)

## Introduction

Sequencing is now a mainstream tool that drives biological and genomics research as well as clinical applications (*1*). Sequencing technologies have been evolving continually with performance improvements and lower costs (*2*), driving broader utilization and development of new applications by offering higher accuracies, higher throughputs, faster turnaround and increasingly simplified workflows.

Nanopore measurement and its potential for sequencing has been in development for over three decades (*3–6*) because of its simple electrical implementation and single-molecule measurement sensitivity. A typical nanopore sequencer has two reservoirs of electrolyte that are fluidically connected through a nanometer-scale pore inserted into an ion barrier membrane. A potential difference between the reservoirs results in a measurable ion current passing through the pore.

If a negative polyelectrolyte such as DNA is introduced to the negative potential reservoir, it can be threaded through a right-sized pore and drawn into the positive potential reservoir. Measurement of the ion current records the level of ion blockage during the polymer translocation through the pore and holds the promise of yielding sequence information.

However, efficient and accurate single-molecule nanopore sequencing of nucleic acids remains challenging because of fundamental signal-to-noise limitations resulting from the poor spatial resolution and molecular distinction of nucleobases. For example, the Oxford Nanopore (ONT) technology (*7*), which directly sequences DNA, is limited by raw read average error rates >3% (sup basecaller) (*8*) and generally poor homopolymer resolution (*9*). While their read lengths lead the industry (*10*, *11*), throughput, costs and accuracy are not competitive with existing platforms (*12*, *13*). As a result, this approach is limited in its application. An alternative to direct nanopore sequencing with new chemistry and measurement innovations is required to meet this difficult challenge.

Here, we introduce Sequencing by Expansion (SBX), a nanopore-based single-molecule approach that overcomes these measurement limitations through a biochemical process that encodes the sequence of a target nucleic acid molecule into a highly measurable surrogate polymer called an Xpandomer (Xp) (*14*). Xpandomers encode the sequence information with high signal-to-noise reporters, enabling high-fidelity, single-molecule nanopore sequencing. We describe the innovations in molecular and protein engineering that enable SBX and demonstrate its performance with data generated using a simple, accessible nanopore set up. When combined with array-based systems, the near real-time high-throughput and workflow flexibility of SBX will allow researchers to rethink how sequencing projects can be implemented.

## Results

SBX was conceived as a solution to the fundamental signal-to-noise challenges of direct DNA sequencing. Our data demonstrates that by spatially expanding the sequence information and adding translocation control features, single-molecule sequencing is achieved with high resolution and throughput [Fig. 1E-G]. Using a systems approach, purpose-built molecular structures and workflows were co-optimized to produce the observed results.

**Fig. 1.**
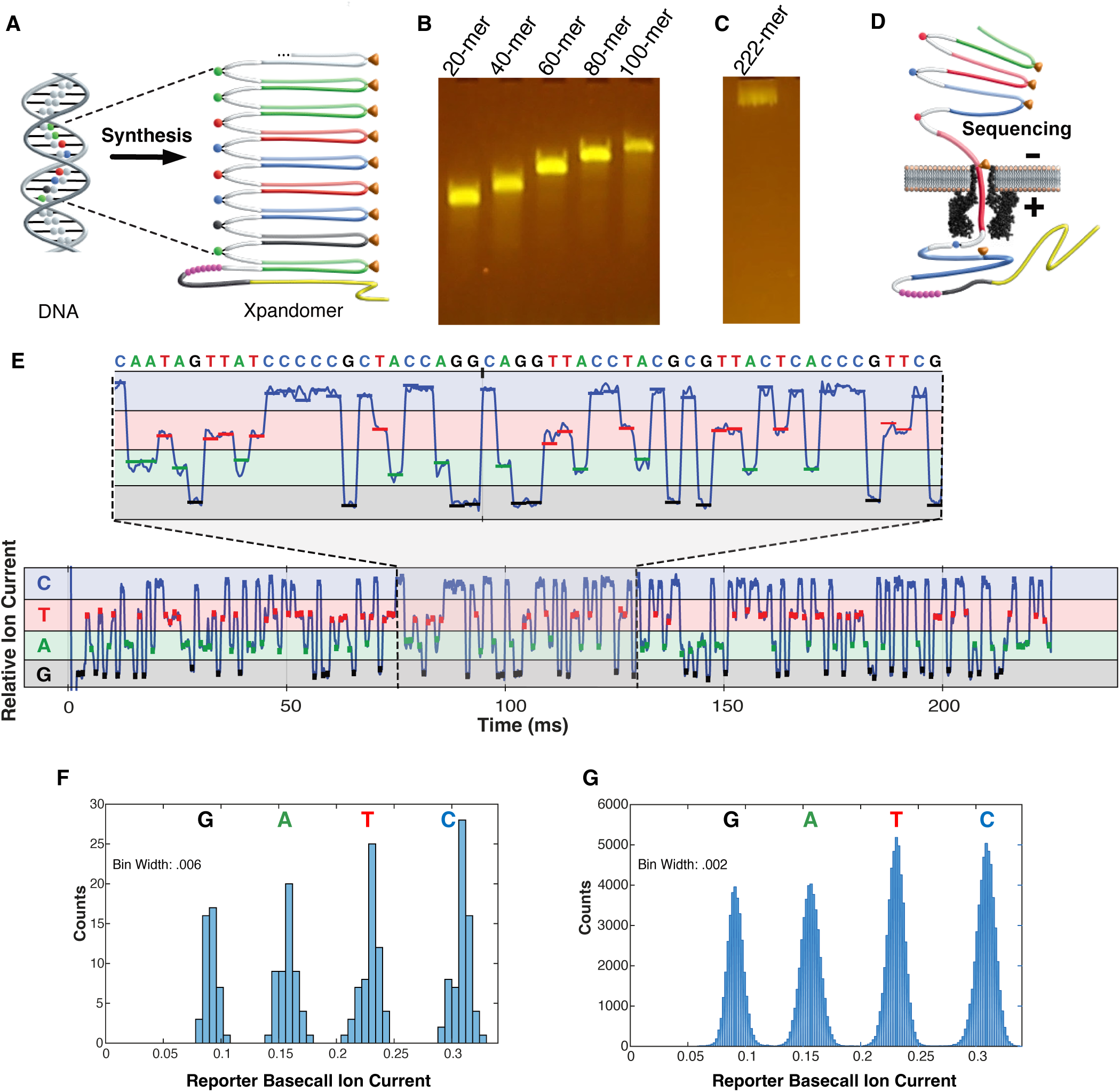
SBX separates chemistry from measurement. **(A)** SBX chemistry synthesizes a surrogate of the DNA called the Xpandomer (Xp). **(B)** Unexpanded Xp synthesized from templates of varying length using a fluorescent primer, analyzed by electrophoresis in a 2.5% agarose gel. **(C)** Expanded Xp synthesized from a 222-mer template using a sequencing-compatible extension oligo (EO). Xp products were analyzed by electrophoresis in a 2.5% agarose gel after hybridizing a fluorescent reporter to the EO. **(D)** A biological nanopore inserted into a lipid bilayer constrains the Xp to sense individual reporters enabling high signal-to-noise, single-molecule measurement and simple sequence readout. **(E)** Example ion current trace recorded during translocation of an Xp synthesized from a 222 base *Streptococcus* DNA template. During translocation, each Xp reporter was paused in the nanopore for a 1 ms period, during which the ion current was sampled. The median ion current derived from samples captured in the last half of the period was normalized by the open pore current (72 pA) to determine the Xp reporter basecall ion current shown as 1 ms colored bars along the above trace. **(F)** Histogram of the reporter basecall ion currents from the above molecular trace. **(G)** Histogram of the reporter basecall ion currents for the set of 966 aligned Xp reads analyzed in this paper.

## SBX Overview

SBX consists of two workflows: synthesis and sequencing. First, during synthesis, the DNA template is enzymatically and chemically converted into the lengthened Xp molecule that contains high signal-to-noise reporters [Fig. 1A-C]. Next, the Xp is sequenced by stepwise translocation through a nanopore, [Fig. 1D] where each step measures a well-defined current specific to one of four reporters that identifies the DNA base [Fig. 1E-G]. Separation of the DNA conversion chemistry from the molecular measurement allows each workflow to be independently optimized to its own unique requirements without compromising the other. This decoupling is a fundamental advantage of the SBX process. These workflows, along with the enabling molecular structures, chemistries and methods are summarized below and are further elaborated in the Supplement.

In the synthesis step, a solid-phase method is used to convert DNA to Xp. A DNA template is hybridized to a specialized, acid-stable primer called an extension oligo (EO) [Fig. 2H], which is anchored to the cyclic olefin walls of a flow channel [Fig. 2A-D]. Next, the primed DNA template is exposed to a reaction mix that comprises a highly-engineered DNA polymerase (Xp Synthase), the four base-specific expandable nucleoside triphosphate substrates (XNTPs) [Fig. 2E, Fig. 3A-B], and a reaction cofactor called a polymerase enhancing moiety (PEM), in a formulation of salts, buffer, solvents and crowding agents optimized for Xp synthesis. The Xp Synthase performs template-dependent primer extension using XNTP substrates under customized reaction conditions.

**Fig. 2.**
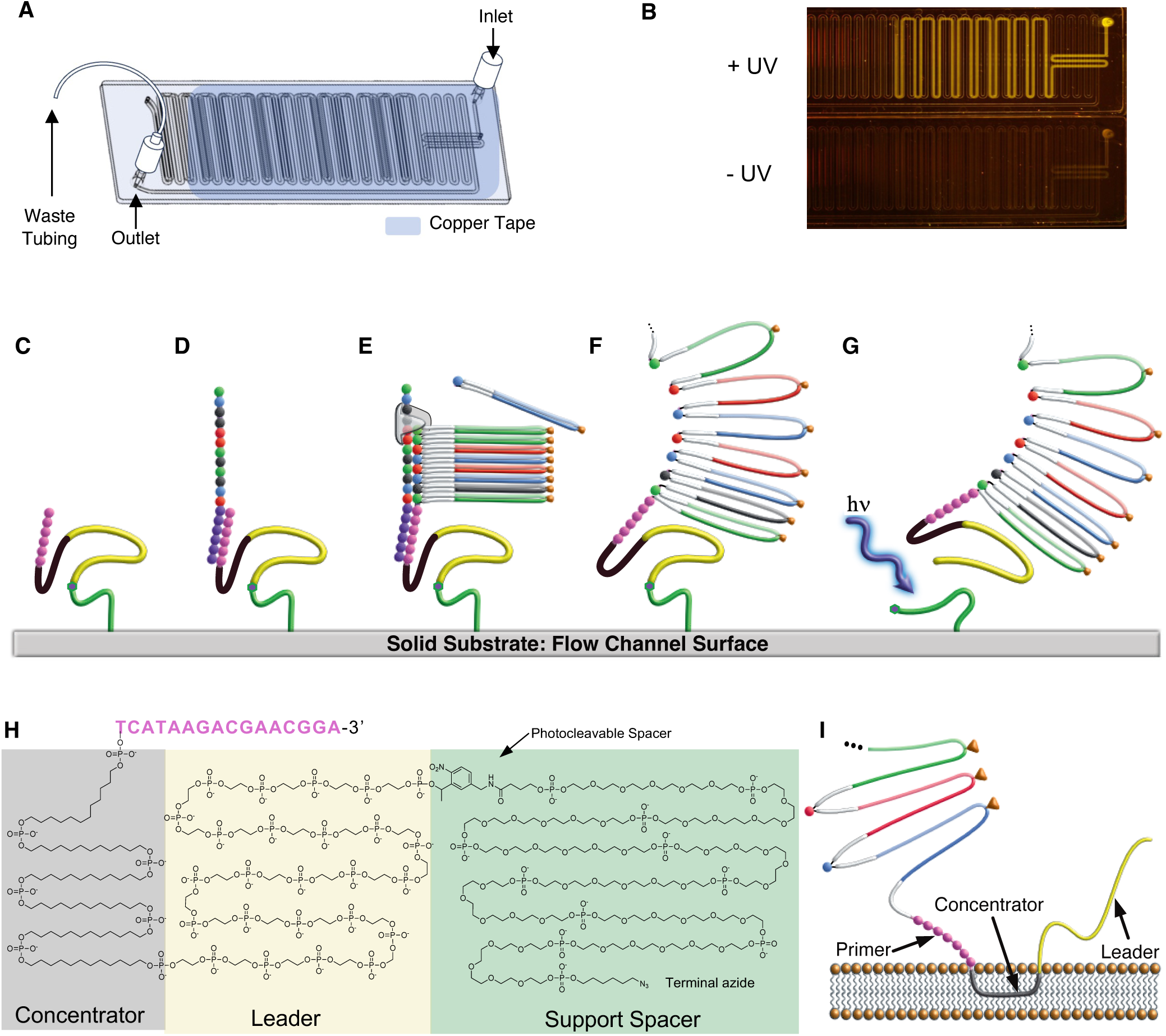
Synthesis of Xp from DNA on solid support. **(A)** Cyclic olefin microfluidic chip (Chipshop, Fluidic 243). 0.6 mm x 0.3 mm x 1,637 mm channel dimension. ∼500 mm^2^ was functionalized to support Xp synthesis. **(B)** Functionalization was carried out in the first 50 µL of the channel in 80% DMSO, 1% SDS, 2 mM propargyl PEG-4 maleimide, which was irradiated (365 nm, 4.2 mW/cm^2^) for 20 minutes followed by a washing step (see Supplemental). For testing purposes, a fluorescent 5’-azide oligonucleotide was clicked to the surface and unbound material was washed away. In the absence of irradiation, minimal background was observed. **(C)** For Xp synthesis, the functionalized solid substrate is conjugated to the EO through click chemistry. **(D)** Target DNA hybridizes to the support-bound EO. **(E)** Xp Synthase performs template-dependent extension with XNTP substrates. **(F)** The Xp is expanded using acid cleavage. **(G)** Release of expanded Xp from solid substrate by photocleavage. **(H)** EO features regions of hydrophobicity (concentrator), high negative charge density (leader), a photocleavable spacer (PC) and a terminal azide for conjugation to the solid support. The 3’ terminus contains acid-stable 2’-OMe nucleotides to hybridize the template and support extension. After photocleavage from support, the leader terminates in a 5’-phosphate. **(I)** When used in sequencing, the concentrator and leader domains carry out specific functions, localizing the Xp to the lipid membrane and promoting capture by the pore.

**Fig. 3.**
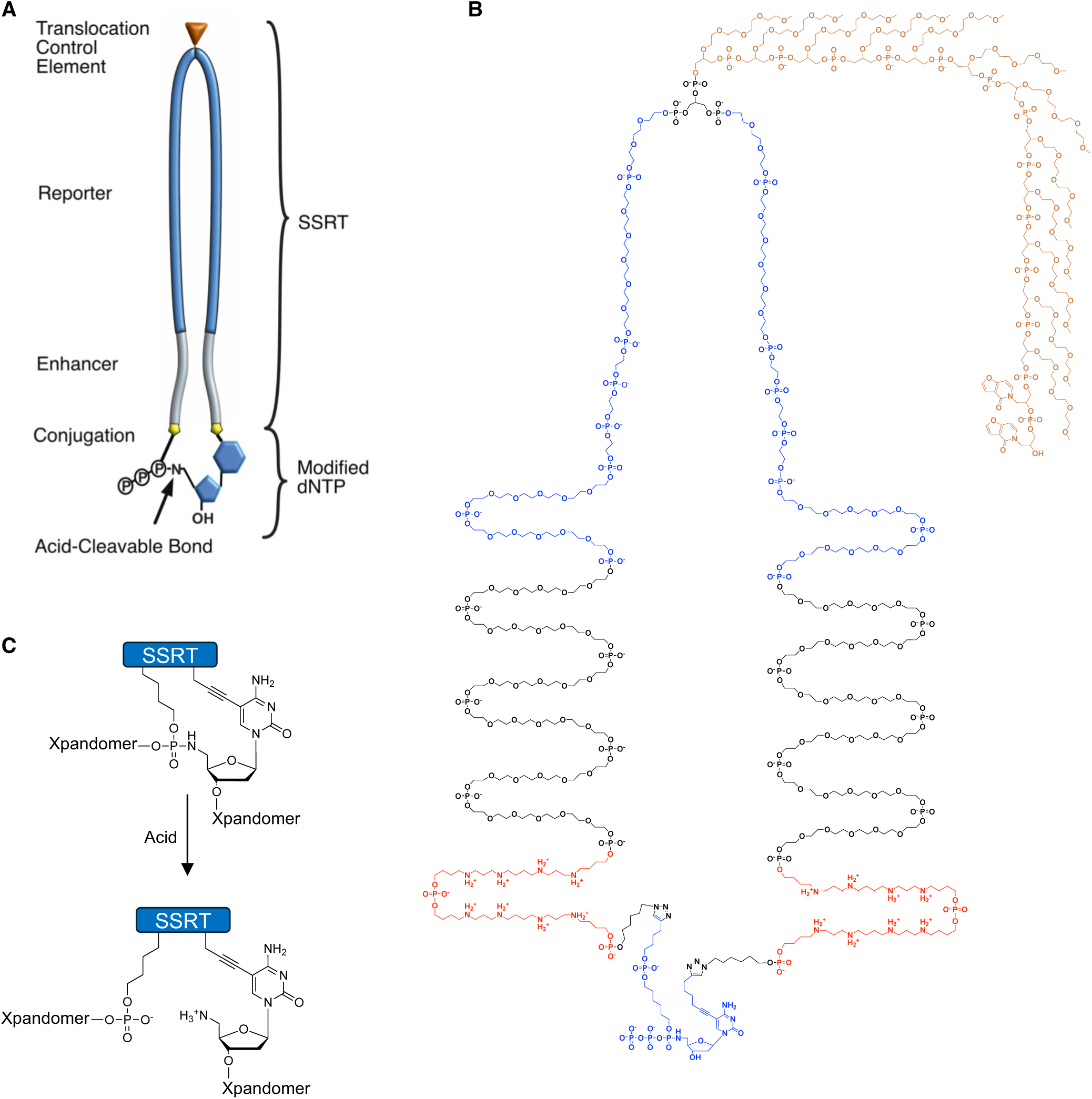
Expandable nucleotide structure. **(A)** Functional schematic of the XNTP. The XNTP is synthesized by double conjugation of the SSRT (symmetrically-synthesized reporter tether) and the modified dNTP. Each SSRT contains an identifying reporter that can be read in the nanopore when held in place by the translocation control element (TCE). Each XNTP also contains an enhancer region consisting of polyamines, which is necessary for efficient incorporation by the Xp Synthase. **(B)** Chemdraw image of XCTP. The SSRT containing the enhancer (red), level 1 (blue) and TCE (gold) is clicked to the bis-alkyne dCTP. The synthesis of the SSRT follows modified oligonucleotide synthesis (See Supplement), with many incorporations of specific spacer phosphoramidites. The average XNTP has a molecular weight of 16 kDa, charge of –38 and can be handled and purified much like a 50-mer oligonucleotide. **(C)** When the XNTPs are incorporated into an Xp, the resulting molecule can be expanded with an acid treatment. The P–N bond is broken, reforming the backbone to include the entire length of the tether.

A subsequent acid treatment [Fig. 2F, Fig. 3C] degrades the DNA template and cleaves each incorporated XNTP at the sugar-phosphate linkage, expanding the Xp backbone to include the length of the tethers located between adjacent nucleobases. Each tether contains a reporter segment corresponding to the identity of the subsequent DNA base. The fully formed Xp is released from the solid substrate by photocleavage [Fig. 2G], and the eluted sample is ready for measurement in the nanopore sequencer.

In the sequencing step, the Xp is measured in the nanopore. Here, we use an alpha-hemolysin (αHL) nanopore inserted into a lipid bilayer membrane which fluidically connects the electrolyte of the *cis* and *trans* reservoirs, and functions as a conduit for the Xp [Fig. 4A-B]. A periodic electrical field is imposed between the *cis* and *trans* reservoirs through Ag/AgCl electrodes [Fig. 4D, Fig. S1A-C]. When Xp is added to the *cis* reservoir, the electrical potential drives it to thread the nanopore [Fig. 4E]. Threading of the Xp occurs at the 5’ end due to two capture enhancement features of the EO: the concentrator and the leader [Fig. 2H-I].

**Fig 4.**
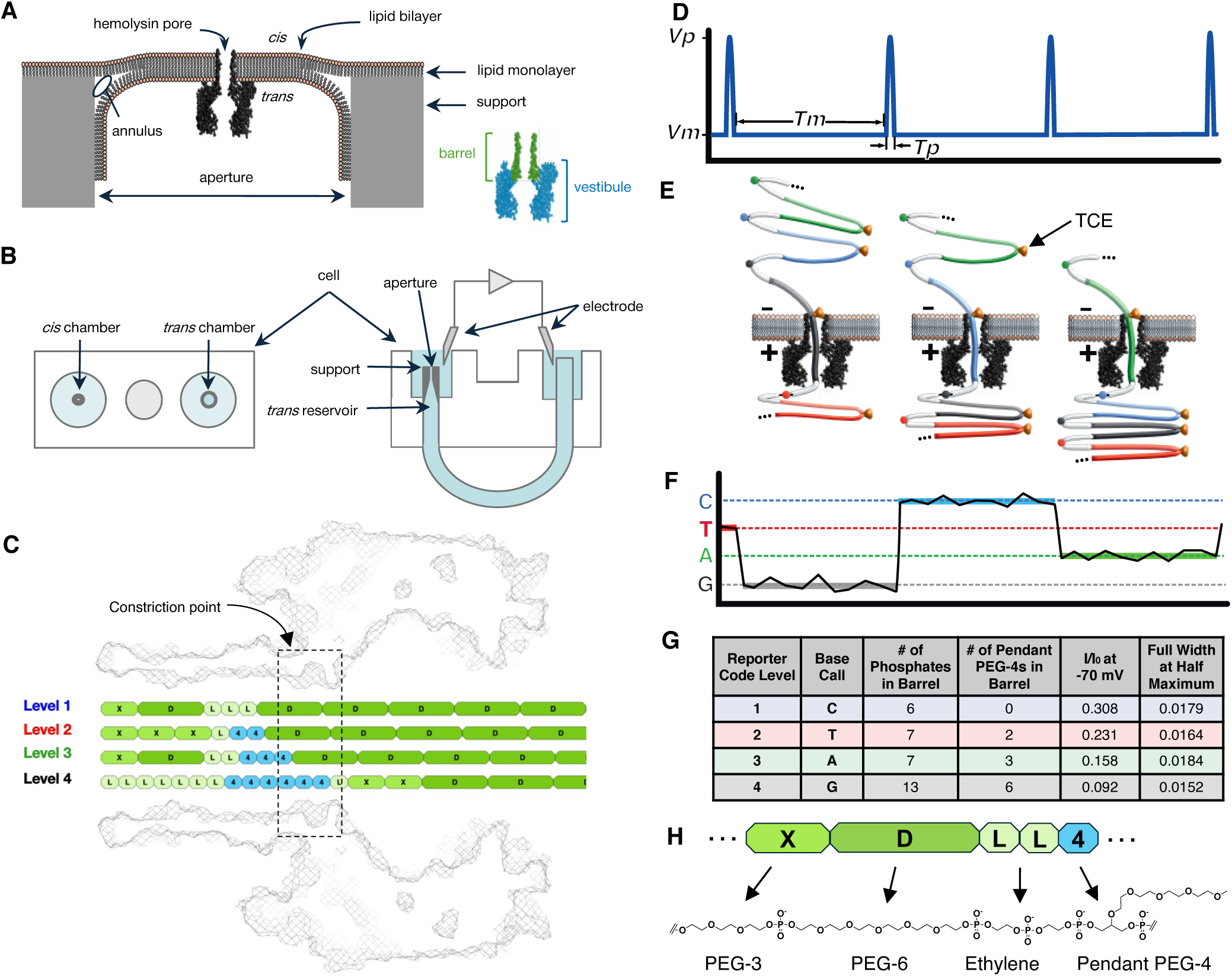
Stepped translocation and measurement of Xp in nanopore. **(A)** Cross-section of the sequencing environment. An assembled heptameric alpha-hemolysin pore is oriented with the barrel facing the *cis* side in a lipid bilayer. The lipid bilayer is formed across the aperture of a PTFE/FEP support. **(B)** Top-down view (left image) and cross-section side view (right image) of the sequencing cell. A PTFE/FEP tube bridges *cis* and *trans* chambers which are each filled with electrolyte solution. A voltage potential is imposed by positioning AgCl electrodes connected to an Axopatch amplifier into the *cis* and *trans* chambers (*44*). After forming the lipid-pore complex, Xp sample is added to the *cis* chamber and driven through the pore via an imposed electric field. **(C)** Base reporters were designed to have four distinct ion current level responses, with similar voltage-pulse translocation control. **(D)** A periodic waveform applied across the nanopore (*cis* to *trans*) consists of a baseline measurement voltage *Vm* and duration *Tm* followed by a short translocation pulse with peak voltage *Vp* and duration *Tp*. Typical times and voltages for the measurement and pulse periods are 1.0 ms / –70 mV and 6.0 μs / –600 mV respectively. **(E)** An Xp in three consecutive measurement states of three consecutive reporters. In each state, the TCE holds the reporter in the nanopore until the pulse advances the TCE through the nanopore to stop at the next TCE along the Xp. **(F)** Depiction of a molecular trace showing three measurement states. **(G)** Characteristics of the reporters producing the best signal levels. The reporter signal values and the FWHM distribution widths for their ion current levels (I) are normalized to the open pore ion current level (I_0_), typically ∼70 pA @ –70 mV. **(H)** Components of the reporter are units of **D** (PEG-6), **X** (PEG-3), **L** (ethylene diol) and **4** (pendant PEG-4) assembled by phosphoramidite chemistry.

After threading, the Xp continues to translocate under the applied potential until paused by a molecular structure called a Translocation Control Element (TCE) [Fig. 4E]. The TCE is positioned to hold its adjacent base reporter in the 5 nm long barrel of the nanopore while the ion current is measured. Four different reporter types [Fig. 4C] yield four distinct ion current levels that correspond to each XNTP base-type [Fig. 4F-G]. The applied waveform sets the period of measurement (typically 1 ms @ –70 mV) and is followed by a brief, high-voltage release pulse (typically 6 µs @ –600 mV) before repeating [Fig. 4D]. The pulse controls the translocation of the TCE, precisely advancing the Xp to the next TCE and subsequent reporter measurement [Fig. 4E]. This cycle continues until the Xp is fully translocated and the nanopore becomes open again to capture another Xp. The resulting ion current trace from Xp translocation shows a series of periodic steps between the four possible levels that are easily translated into the DNA sequence [Fig. 4F].

## Molecular Structures, Chemistries and Methods that Enable SBX

Novel molecules, synthetic routes and enzymes, along with specialized solid-phase synthesis workflows, reagent formulations and sequencing methods were developed for this technology. The molecular design required hundreds of iterations of the XNTP and the EO, a process facilitated by the strategic decision to leverage phosphoramidite chemistries. With in-house capability and access to >200 phosphoramidites, designs could be routinely conceived, synthesized and tested within days. The modular design of molecular systems meant that separate functional parts of these molecules, such as the base reporters and enhancers, could be optimized with relative independence from each other. Similar systems approaches were used in developing PEMs [Fig. S2] and the Xp Synthase. With few exceptions, SBX optimization required the nanopore measurement to resolve molecular level effects [Fig. 5A-C]. Here, these designed elements and workflows underlying SBX chemistry and sequencing are described in detail, followed by a demonstration of the method.

**Fig. 5.**
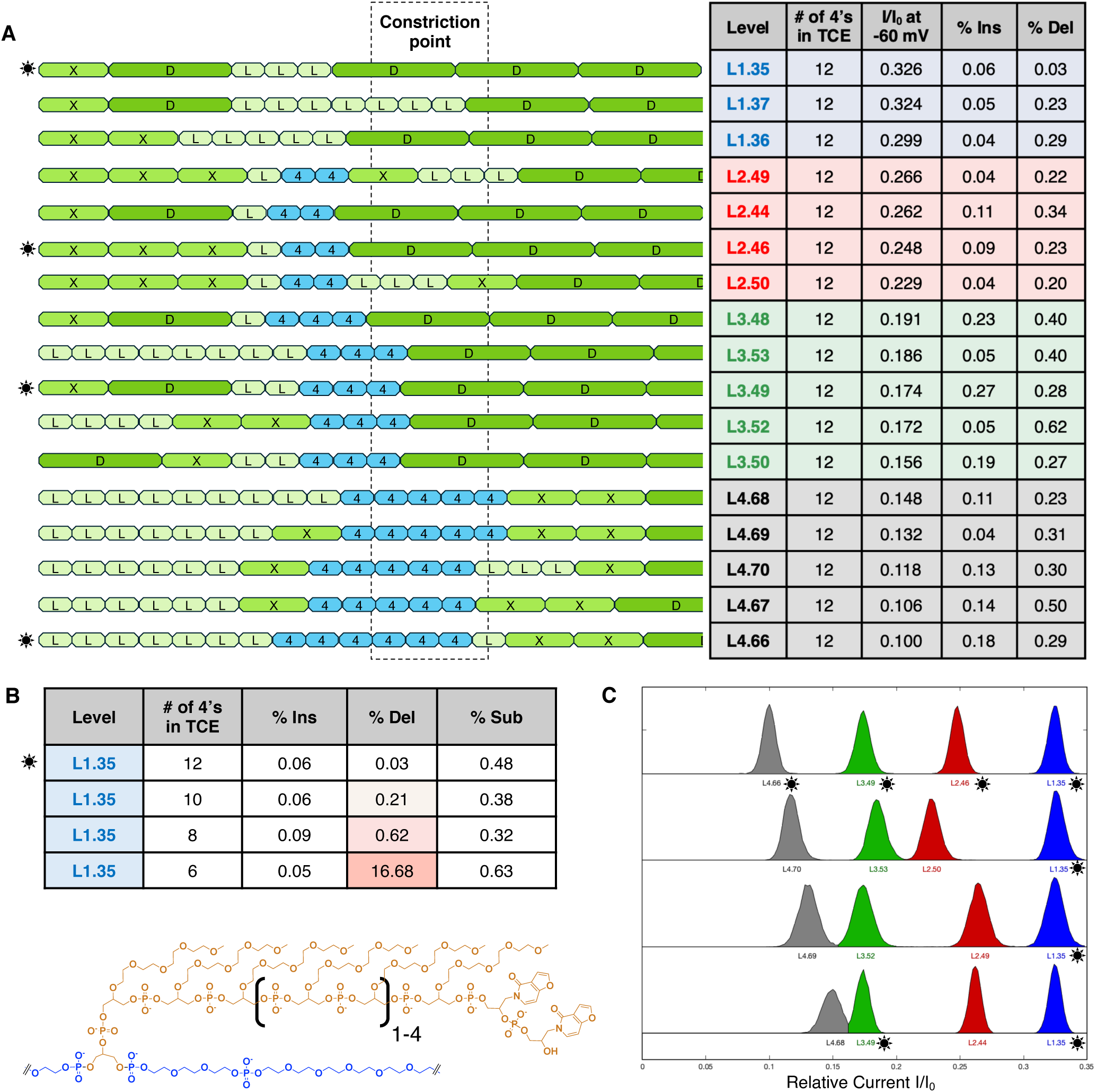
SSRT composition determines control over levels, translocation and noise. **(A)** Various reporter code designs with corresponding performance data. When bulky pendant PEG-4 units (light blue) are added to the reporter code, the observed I/I_0_ decreases. For a given number of pendant PEG-4 residues, additional tuning of their placement can change the I/I_0_. The nanopore constriction point has the most sensitivity for ion current adjustment. Structures used for the sequencing demonstration are highlighted with stars and were chosen for their separation and signal quality. **(B)** Changing the length of the TCE for a given level had little impact on the I/I_0_, but a profound impact on the deletion profile. In this example of a Level 1, a TCE of 6 pendant PEG-4 units loses its ability to stall the reporter code in the barrel of the nanopore. For the sequencing data presented herein, all TCEs had 12 pendant PEG units. **(C)** As I/I_0_ varies with reporter code design, the levels can become less resolved as the signals begin to overlap, resulting in non-enzymatic substitution errors.

## Expandable Nucleoside Triphosphate (XNTP)

The key building blocks of Xp synthesis are the four XNTPs [Fig. S3-S6], one for each nucleotide. Each of the XNTPs uniquely encodes the base identity of its cognate nucleotide and serves as the substrate for Xp Synthase. The XNTP consists of a modified deoxynucleoside triphosphate (dNTP) that is doubly-conjugated to a symmetrically-synthesized reporter tether (SSRT) to form a cycle using copper-catalyzed azide-alkyne “click” chemistry (*15*). The SSRT is synthesized using phosphoramidite chemistry which enables precise design and rapid optimization of the XNTP’s molecular features, including high signal-to-noise reporters and non-stochastic translocation control [Fig. 6A-B]. A glossary of all phosphoramidite structures used is shown in Fig. S7.

**Fig. 6.**
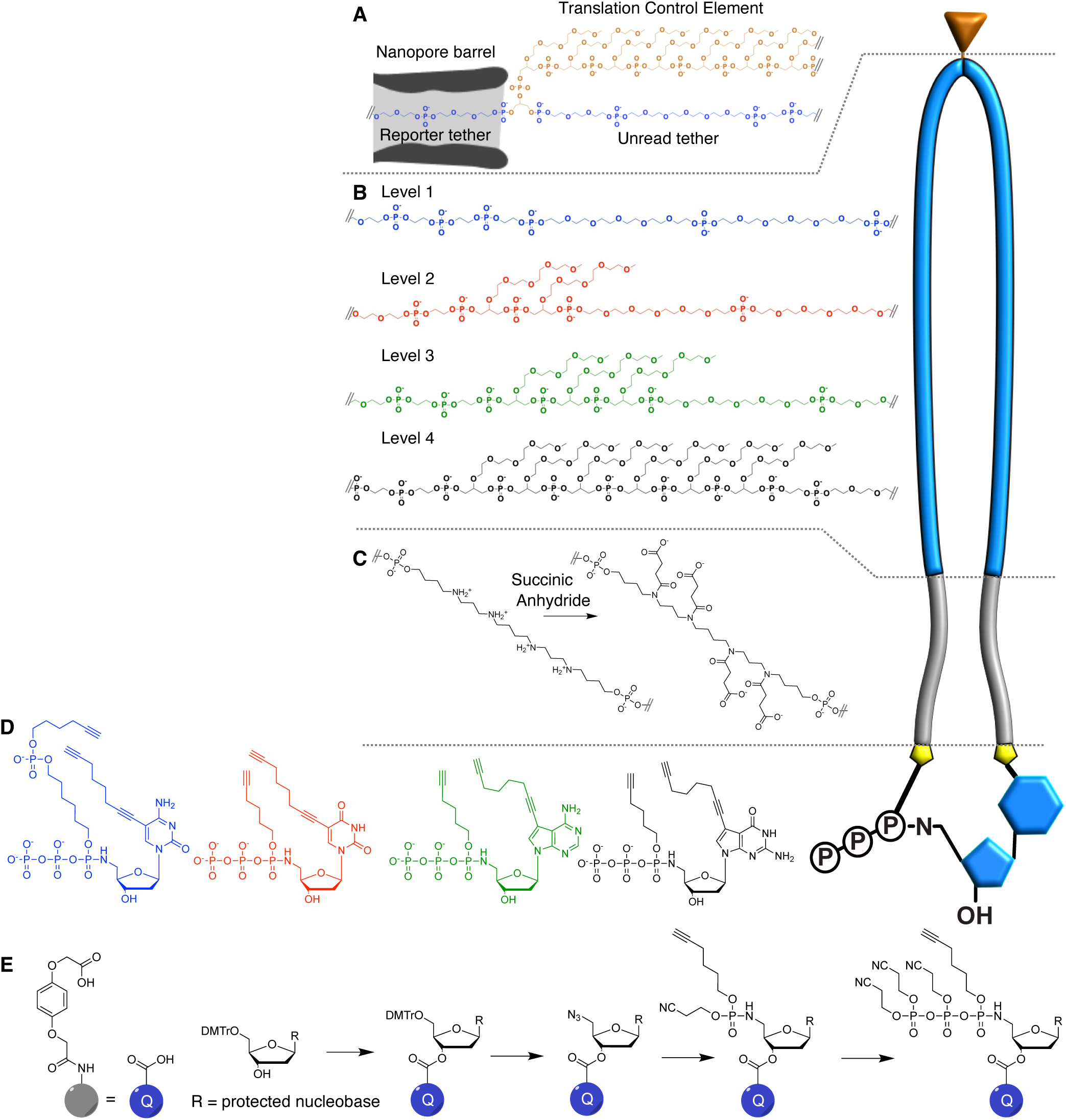
Anatomy of the XNTP. **(A)** At the measurement potential, the TCE (gold) holds one reporter tether in the barrel of the pore while folding over on the unread tether. During the translocation pulse it is squeezed through the pore and the Xp pauses at the next TCE. **(B)** The reporter contains spacer and pendant PEG units. The quantity and positioning of the pendant PEG units leads to a characteristic blockage (I/I_0_) for each level. **(C)** The polyamine enhancer region is positively charged and enables Xp synthesis. While being measured, excess positive charge can produce noisy levels. To mitigate this, the Xp undergoes a post-treatment with succinic anhydride, neutralizing the charge. **(D)** The four modified dNTPs are doubly-functionalized with alkynes to enable bis-click. **(E)** The dNTPs were synthesized on solid support by converting the 5’-azide to a phosphoramidate, followed by iterative phosphorylation.

Where we could not achieve the desired performance with commercially available products, we synthesized new phosphoramidites [Fig. S8A-C], developing a diverse repertoire of these building blocks to optimize SSRT structure and function. Solubility, length, spacing of charge and positioning along the reporter code were engineered to enhance performance. We synthesized the SSRT as a symmetric molecule by utilizing a branching phosphoramidite [Fig. S9]. This simplifies the final double conjugation of dNTP to the SSRT. XNTP and Xp function is not affected by the orientation of the SSRT-dNTP conjugation due to the symmetry of the SSRT. The XNTP macrocycle products showed an expected lower gel mobility and were easily separated from the linear SSRT by HPLC [Fig. S10A]. Through control of the substrate concentration, SSRT structure, and click reaction conditions we isolated pure cyclized product in excess of 50% yield [Fig. S10C]. The click reaction is known to produce reactive oxygen species that can be detrimental to nucleotides, so XNTP click reactions were performed under argon using minimal copper to avoid these products (*16*). During XNTP isolation, uncyclized and concatemer click products were also removed [Fig. S10B-C]. Any side product other than cyclized XNTP, with a single acid-cleavable bond, could lead to a strand break in the final Xp.

## Symmetrically Synthesized Reporter Tether (SSRT)

### Translocation Control Element (TCE)

Controlling DNA movement through a nanopore has been previously considered (*17*, *18*). The TCE regulates Xp translocation through the nanopore as described above. The structure is composed of a series of pendant polyethylene glycol (PEG) residues (*19*), followed by a symmetric brancher to which the remaining features of the SSRT are constructed through stepwise synthesis. Fig. 6A illustrates our hypothesis of how the TCE pendant PEGs fold back at the nanopore entrance and sterically prevent translocation under the low force of the applied measurement potential (*20*, *21*). In contrast, the TCE will pull through the nanopore barrel under the brief high force delivered by the voltage pulse that follows measurement. This periodic measurement process provides well-controlled, reliable performance, with few incidents of stochastic TCE translocation, which simplifies analysis and allows for highly efficient measurement [Fig. 1E]. Translocation fidelity is important, as failure to advance the Xp by a single reporter per pulse results in either a deletion or insertion sequencing error.

### Base Reporter

During Xp nanopore measurement, the TCE precisely positions the reporter in the barrel of the nanopore, impeding the ion flow to one of four current measurement levels associated with each of the four XNTP base types [Fig. 6B]. Base reporters are composed of a series of spacer and pendant PEG phosphoramidites, experimentally optimized to reproducibly generate distinct, well-separated current levels, with minimal noise during measurement. Design considerations included size, charge density, chemical properties and precise positioning in the nanopore. Due to the interdependent function of the reporters and TCE, molecular design optimization occurred simultaneously, to balance both the DC measurement voltage and the release pulse for each reporter [Fig. 5A-C]. Although the SSRT is synthesized symmetrically, only the leading arm occupies the nanopore to generate a base reporter ion current measurement.

### Enhancer

This SSRT functional group is essential for efficient Xp synthesis [Fig. 7A] and is composed of polyamine phosphoramidites [Fig. 3B] that provide positive charge density. While it is not fully understood how the enhancer region promotes Xp synthesis, this part of the structure likely interacts with the Xp Synthase, where it could encourage better XNTP binding or reduce the Xp-enzyme dissociation rate. It is also possible that the charge creates organization of the SSRTs in the growing Xp which stabilizes synthesis. The enhancer’s positive charge density, while beneficial for Xp synthesis, requires chemical modification prior to Xp sequencing in the nanopore to mitigate measurement interference [Fig. 6C].

**Fig. 7.**
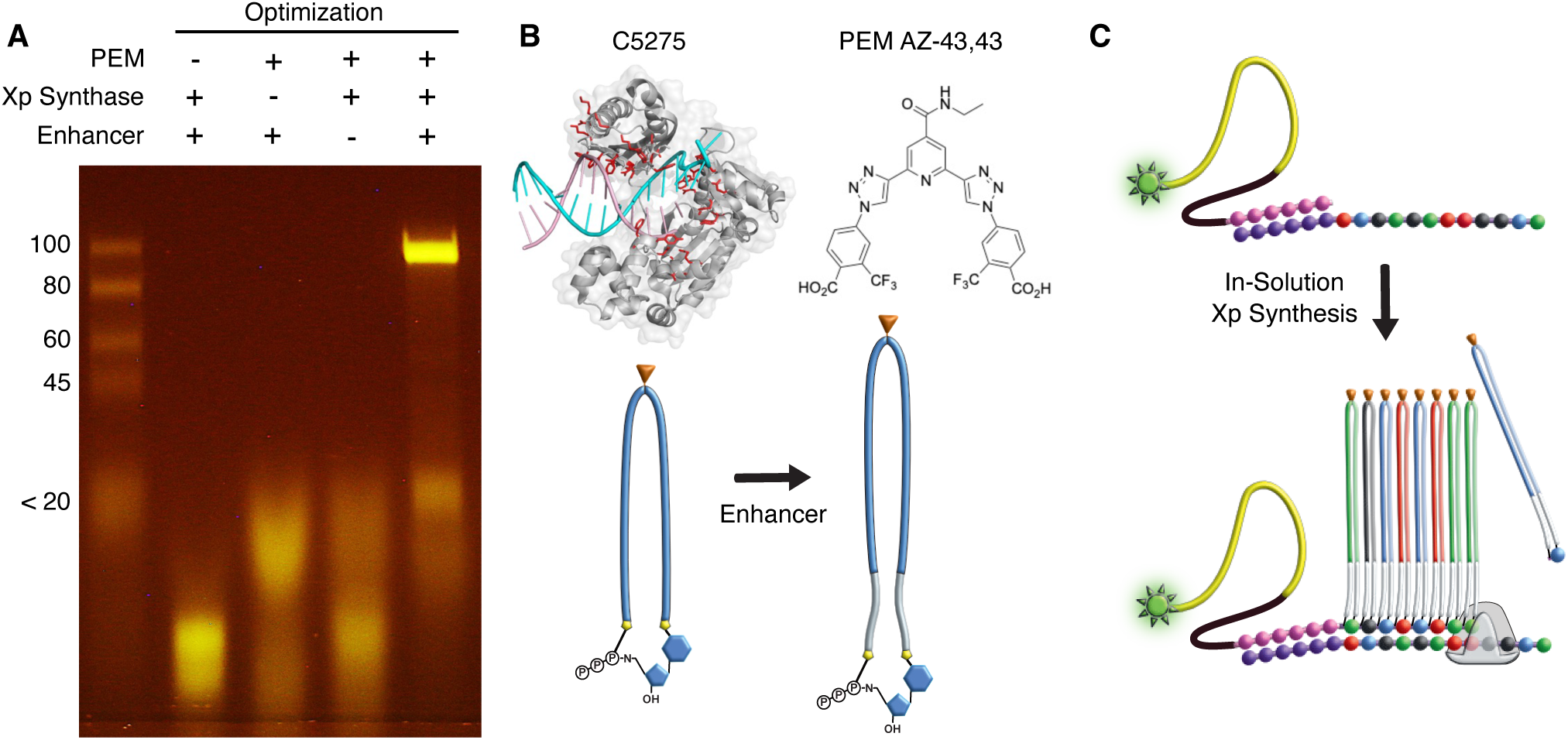
Critical elements in the synthesis of Xp. **(A)** PAGE analysis of uncleaved Xp products synthesized from a 100-mer template in solution. Reversions that eliminate any critical element prevent formation of full-length product. **(B)** Reversions in **A** included WT DPO4 replacing Xp Synthase C5275, removal of PEMs and XNTPs missing the enhancer region. **(C)** In-solution synthesis using a 5’-SIMA HEX labeled EO permitted rapid testing of components for optimization of Xp synthesis. In-solution extensions were performed similarly to on-support extensions but were quenched with EDTA and proteinase K. Products from templates shorter than 100-mer can be analyzed by 4-12% polyacrylamide TBE gel, and longer Xp can be analyzed by 2.5% agarose TBE gel.

### Azide Terminal Ends

Following the enhancer, the final functional element added to each SSRT arm is a 6-carbon terminal azide linker [Fig. S11] that is used for macrocyclization of the SSRT and modified dNTP via click conjugation (*22*, *23*).

## Modified dNTP

### Acid-Cleavable Phosphoramidate Bond

The alpha phosphoramidate modification (5’ P–N–C 3’) of the dNTP permits template-dependent extension of XNTPs and enables selective cleavage downstream (*24*). After Xp synthesis, the P–N bond is cleaved by treatment with strong acid, which opens the macrocycle to form the expanded Xp backbone to include the length of the SSRTs [Fig. 3C]. Since all other components of the Xp were engineered to survive acid, the expanded molecule provides well-separated, signal-distinct reporters that can be measured independently [Fig. S12A-C].

### Alkyne Linker Arms

To enable macrocyclization via a bis-click strategy (*15*, *25–27*) the modified dNTPs have alkyne-functionalized arms [Fig. 6D]. Octadiynyl arms (*28*) were used for the heterocycle linkage (C-5 for pyrimidines, deaza-7 for purines) while hexynyl arms were attached to the alpha phosphoramidate of A/T/G. With this configuration, cleavage of the P–N bond allows the Xp molecule to expand. The structural characteristics of the alkyne arms have a large influence on the incorporation efficiency of the XNTP [Fig. S13A-C]. For example, to increase accuracy and prevent misincorporation of XCTP, a compound arm consisting of hexyl-phosphate-hexyne was used to normalize incorporation efficiency relative to XTTP. Without this adjustment, T to C substitution errors were increased [Fig. S13D].

### Synthesis of Modified dNTPs

Synthesis of such a highly modified dNTP requires a specialized synthetic route [Fig. 6E, Fig. S14A]. The electronics of the 5’-aminonucleoside make classical dNTP synthesis routes such as the Yoshikawa (*29*) and Ludwig-Eckstein (*30*) methods extremely low yielding. Other routes employing trimetaphosphate are incapable of placing the linker arm on the alpha phosphate (*31*). Ultimately, we developed a method based off Engel’s chemistry, running through the azide intermediate (*15*, *32*). When incubated with the phosphite of an alkyne arm, the resulting Staudinger reduction can undergo a Michaelis-Arbuzov rearrangement (*33*) to produce the functionalized monophosphate. This chemistry was adapted to a solid support (*34*) to continue building the triphosphate through iterative phosphorylation with phosphoramidites (*15*, *35*).

While modified nucleotides have seen significant utility in many applications (*31*), this is the first report of dNTPs containing a 5’-phosphoramidate diester. These modified dNTPs are stable in neutral to alkaline buffers but the phosphoramidate breaks down under acidic conditions. As expected from the geometry of the Dpo4 active site, only one isomer at the alpha phosphoramidate is a substrate for the Xp Synthase. Therefore, the diastereomers were separated by HPLC prior to click conjugation [Fig. S14B].

## Extension Oligo (EO)

The EO [Fig. 2H-I] comprises three functional components: the leader, the concentrator and the Xp primer. The EO has additional functional groups on the leader end that support solid phase synthesis, including an azide for click conjugation to an alkyne-functionalized cyclic olefin substrate and a photocleavable linker to release the Xp after synthesis [Fig. 2G, Fig. S15]. The Xp primer is synthesized with acid-stable 2’-O-methyl A, C and G modified bases, and unmodified acid-stable 2’-deoxy T bases to prevent breakdown during the cleavage step. The primer portion of the EO hybridizes to the DNA template and initiates Xp extension from the 3’end.

The leader and concentrator components promote capture of the Xp by the nanopore [Fig. S16A-B]. The leader, on the 5’-terminus of every Xp, is a linear polymer element with high negative charge density designed to maximize capture by the nanopore biased with a positive potential on the trans reservoir. Following the leader is the concentrator, a hydrophobic linear polymer element designed to associate with, and diffuse along, the surface of the lipid bilayer. This acts to confine the leader near the lipid surface, which increases Xp capture rates by greater than 1000x [Fig. 2I, Fig. S16B].

## Polymerase Enhancing Moiety (PEM)

In an effort to increase the average length of Xp synthesis to >10 bases, we discovered that some classes of DNA minor groove binders improved Xp extension length dramatically [Fig. S17A-B]. Based on this finding, we developed a novel family of molecules using a bis-alkyne core that could be double-conjugated to a variety of azide arms (*36*). With a systematic synthesis and screening effort, we further discovered higher-performing combinations, using a large matrix of cores and arms [Fig. S2A-B]. These molecules, which are termed Polymerase Enhancing Moieties (PEMs), have a profound effect on Xp Synthase processivity, enabling Xp synthesis lengths that exceed 400 bases [Fig. 7A, Fig. S2E].

Several molecules in our initial screening demonstrated a PEM effect on a 45-mer template. Both the angle of the arms and the presence of the salicylic acid correlated with improved extension [Fig. S2C-D]. We performed a second round of testing using a 100-mer template and varying the concentration of PEM [Fig. S2D]. While the PEM effect did increase in a concentration dependent fashion, when the concentration of the additives reached 1.5 mM there was a noticeable drop off in Xp extension length. The lead PEM after these screens (D-8,8) contained a 2,6-pyridine core with *p*-salicylic acid arms.

We next targeted select regions of the PEM for structural improvements [Fig. S18]. When the 4 position on the pyridine core was modified to the nitro, nitrile or *N*-ethylamide the PEM effect increased [Fig. S2C]. Additionally, the solubility improved which allowed use of up to 15 mM PEM in the extension reaction. The arm regions were made more acidic by replacement of the salicylate alcohol with the electron-withdrawing trifluoromethyl substituent. This PEM, called AZ-43,43 [Fig. 7B] resulted in full length Xp of a 411-mer template [Fig. S2E], and was used in the SBX demonstration described below. A potential mechanism for the PEM effect is the inherent instability of the DNA-Xp heteroduplex. Without the charged backbone of DNA, the DNA-Xp heteroduplex could fray once it has passed beyond the influence of the Xp Synthase, at about 9-10 bases in length, resulting in slippage and stalling further synthesis. X-ray crystallography or cryogenic electron microscopy studies of the Xp Synthase tertiary complex with PEMs could provide more insights into their function.

## Xp Synthase

The size of the SSRT and the uncharged phosphoramidate diester of the modified dNTP create significant barriers to primer extension with XNTPs by most DNA polymerases. Dpo4 (*Sulfolobus solfataricus*), a Y-family DNA polymerase, has an open DNA binding cleft (*37*, *38*) which provides a continuous passage for the large SSRT structures [Fig. 8A]. Wild-type Dpo4 was capable of only 1-2 XNTP incorporations under the original extension conditions with early SSRT designs [Fig. S19A]. Through a mutagenesis process, we evolved an Xp Synthase with 36 amino acid substitutions [Fig. 8B, Table S1] from wild-type (*39*, *40*), which is capable of generating Xp that are hundreds of bases in length under optimized conditions [Fig. 7A-C].

**Fig. 8.**
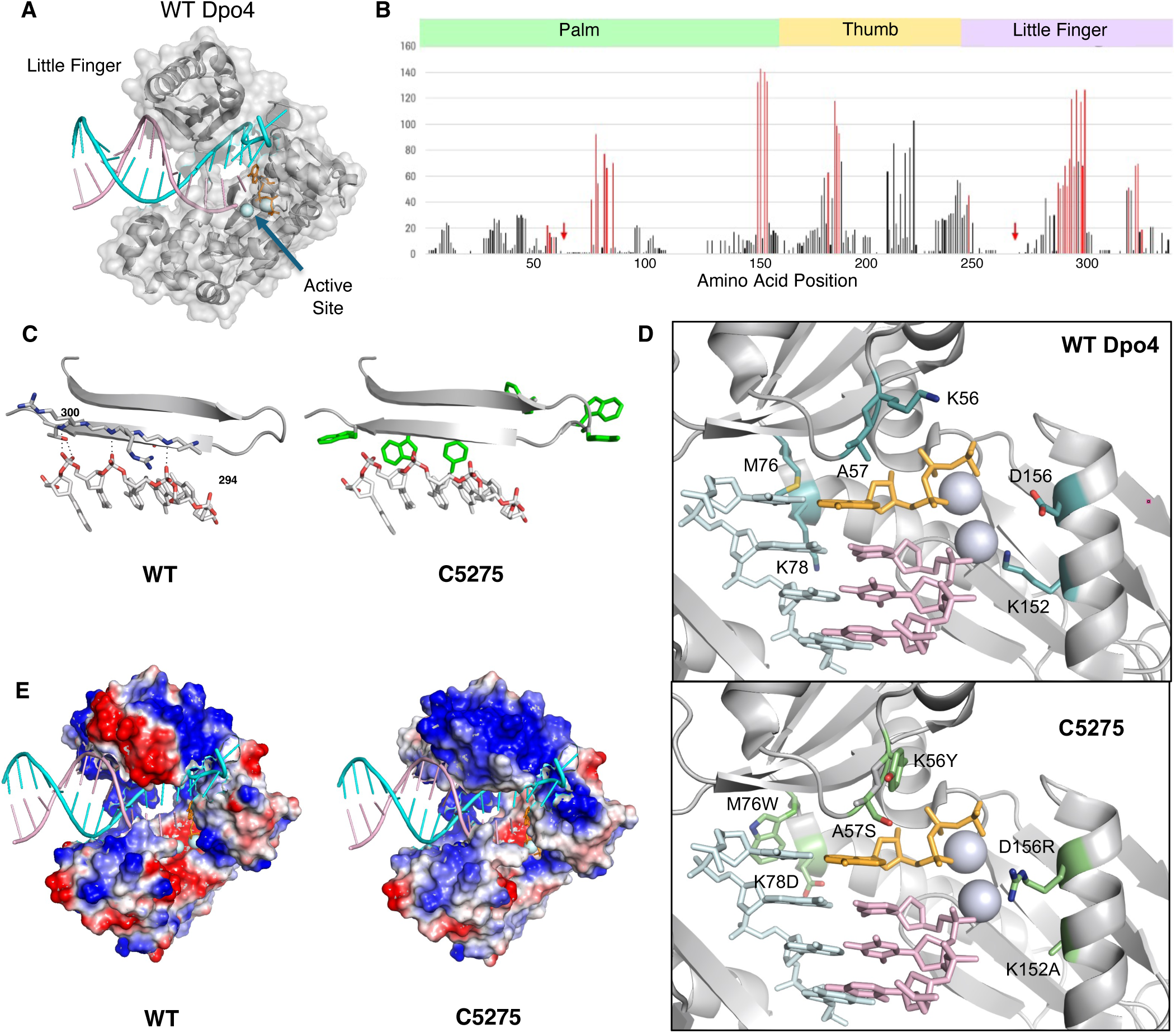
Evolution of Dpo4 polymerase into an Xp Synthase. **(A)** Dpo4 was selected as the parent DNA polymerase because of its open DNA-binding cleft that provides continuous passage for the XNTP reporter tethers, about 9-10 XNTP incorporations. PDB accession number 2AGQ with template DNA (cyan) and product strand (rose) was used for all structures. **(B)** Histogram of the number of times an amino acid position was targeted by library mutagenesis. The count includes the number of times a position appeared in a unique library, and the number of times that library was screened. Red indicates positions where mutations were adopted. The red arrows indicate positions 63 and 272, which are mutated in the Xp Synthase but were not targeted by library screening. **(C)** Little finger primer rail positions 293-301 showing WT Dpo4 positions and Xp Synthase mutations modeled in PyMol using lowest strain rotomer orientation. There are key hydrogen bonds (dotted lines) between the DNA backbone and peptide backbone at positions 295, 297, and 301 in the WT structure. The Xp Synthase has six aromatic positions added in this region (V289W, D292Y, L293W, S297Y, R299W, T301W) (green). **(D)** Dpo4 active site comparison showing key mutations that reshape the XNTP binding site. **(E)** Changes in surface charge of Dpo4 from WT to C5275 generated using the PyMol vacuum electrostatics tool. Red indicates higher average negative charge and blue indicates higher average positive charge. Mutations on the open face of the Xp Synthase that change the surface charge: K56Y, E63R, K152A, D156R, E291S, D292Y, D294N, R300S, K321Q, E324K, E325K, E327K.

For the initial phase of mutagenesis, we used large libraries in which 1-8 amino acids were targeted simultaneously for saturation mutagenesis to survey the polymerase, with a focus on regions proximal to the catalytic site and nucleic acid interaction surfaces [Fig. S19B]. Next, we used small libraries and designs focused on regions that generated hits in the initial surveys [Fig. S19E]. We evaluated variant polymerases for extension length using XNTP precursor substrates in a high-throughput functional screen [Fig. S19 C-D]. Most of the adopted amino acid substitutions occurred at nucleic acid binding surfaces, and probably reflect adaptations to the significant difference in charge between the DNA and Xp backbones. These modifications improved Xp extension length and yield, while fidelity was primarily affected by mutations near the catalytic site [Fig. 8D, Table S1]. Most base substitutions in SBX sequencing derive from Xp Synthase base selection errors and can be addressed by mutagenesis. For example, the A57S mutation led to a 1% net reduction in G(A) substitutions, and the V42A reversion improved all base substitutions [Fig. 8D, Table S1].

Without a structure of the Xp Synthase, it can be difficult to assign a mechanism for how individual mutations contribute to the observed improvements in SBX extension, accuracy, soluble yield or thermostability, but there are two observable trends of note: 1) Substitution for aromatic side chains is preferred at many positions that interact with the extending Xp strand [Fig. 8C]. 2) Reducing negative charge on the open side of the polymerase improves average extension lengths [Fig. 8E]. The phosphoramidate diester backbone of the Xp molecule is uncharged, so the preference for aromatic side chains on the primer interacting surfaces may indicate a new way of engaging with that strand. The reduction in negative charge on the open side of the polymerase was engineered to prevent undesired electrostatic interactions between the polyamine enhancer moieties on the SSRTs and the polymerase surface. The mechanistic basis of how the remaining mutations improve polymerase performance will be a topic of future study.

## Xp Synthesis Considerations

We optimized the extension master mix to balance yield and accuracy of Xp synthesis and to reduce template secondary structure. Adjustments to concentrations of PEMs, XNTPs, Mn^2+^, PEG, solvent and salts as well as buffer pH resulted in accuracy, extension length and yield outcomes that are consistent with the high interactivity of these reaction components. In particular, accounting for the broad interactivity of divalent Mn^2+^ was critical to balancing the reaction. Mn^2+^ is an essential cofactor for catalyzing the template-directed addition of XNTPs to the primer strand, and it cannot be substituted by Mg^2+^ without a significant loss of yield [Fig. S20A, D]. As in PCR, Mn^2+^ contributes to base substitution errors at elevated concentrations, but also enhances the catalytic efficiency of the enzyme [Fig. S20F] (*41*) (*42*) so understanding what factors can alter the effective Mn^2+^ concentration is key to optimizing the reaction. Oxidation of Mn^2+^ during Xp synthesis, which reduces the concentration of the divalent form, leads to slower reaction kinetics and reduced full length yield [Fig. S20B], requiring consideration of pH (*43*), reaction byproducts and reagent purity. It also led us to prepare the master mix immediately before use, with MnCl_2_ added last. PEMs and PEG both influence the effective concentration of Mn^2+^. PEMs appear to weakly chelate Mn^2+^, while the crowding effect of PEG may make Mn^2+^ more available [Fig. S20G, H]. Thus, the development of the SBX extension reaction often revolves around balancing desired primary effects of reaction components with how they might alter the effective Mn^2+^ concentration.

The solvent environment is also important to reaction formulation. Molecular crowding by inclusion of PEG 8000 at 20% wt./vol. was essential to achieve the reaction kinetics observed for Xp synthesis. Decreasing concentration to 16% wt./vol., for example, dramatically reduced Xp yield [Fig. S20C]. The co-solvent N-Methyl-2-pyrrolidone (NMP) improves accuracy at the expense of extension length [Fig. S20I]. The amount of solvent also affects the solubility of PEMs, with increased concentrations of solvent enabling the use of higher concentrations of PEMs [Fig. S20E]. Given that PEMs can affect accuracy as well as length, in a concentration-dependent manner [Fig. S20H], the interplay between solvent and PEMs is critical for balancing the reaction conditions.

## Sequencing Considerations

The choice of materials, design and protocols evolved to improve the quality and throughput of Xp sequencing in the nanopore. Using 1,2-di-O-phytanoyl-*sn*-glycero-3-phosphoethanolamine (DPhPE) lipid in the nanopore bilayer resulted in a >6-fold increase in Xp throughput over diphytanoylphosphatidyl choline (DPhPC), which could be attributed to better concentrator-lipid interaction [Fig. S16C]. We also evaluated the αHL orientation in the bilayer [Fig. S21A-D]. The barrel-to-vestibule orientation demonstrated higher throughput and much lower measurement noise relative to sequencing in the vestibule-to-barrel orientation. An essential feature of Xp sequencing is the inclusion of a negatively-charged leader and a hydrophobic concentrator on the EO [Fig. S16]. We extensively optimized both features for length and composition. The concentration and choice of sequencing salts also had a significant impact on signal quality and Xp measurement behavior. 1 M KCl is traditionally used for nanopore measurement (*44*), however, NH_4_Cl demonstrated the highest signal-to-noise and best overall sequence quality of all the salts evaluated. LiCl in particular had poor ion current signal, possibly due to lower relative ion mobility, and also negatively affected Xp translocation, demonstrating slower, less consistent translocation when compared to the NH_4_^+^ salt. Chaotropic salts, like GuHCl or Urea, were found to improve Xp yield and length when added to the *cis* reservoir at a concentration of 1 M.

## SBX Sequencing of 222 Base DNA Template

To demonstrate the single-molecule signal resolution of SBX, we combined the chemistries and methods above to synthesize and sequence an Xp derived from a 222-base *Streptococcus* DNA template. Additional supporting details for the protocol and reagent mixes are provided in the Supplement.

To generate the highest quality sequencing data, we performed solid-phase Xp synthesis, cleavage and modification steps using an off-the-shelf cyclic olefin chip (Chipshop, Fluidic 243) [Fig. 2A-B] with a functionalized flow channel that allowed for tight control of reaction timing, temperature and introduction of reagents. This approach facilitates removal of reaction components that would affect the stability of the lipid bilayer and overall sequence quality, such as solvents and detergents. All liquid handling steps were controlled by manual pipetting, and utilized 500 mm^2^ of the flow channel surface area (∼278 mm channel length) for synthesis. The flow channel was functionalized by propargyl-PEG4-maleimide addition, followed by irradiation (*45*). The template was selected for its sequence composition, which includes hairpins, homopolymer regions and dinucleotide repeats [Fig. 9C-D]. To enable efficient hybridization of the template to the EO, we strand-enriched the DNA template using established biotin-streptavidin methods (*46*). After flushing the click reagents and excess EO from the chip, we hybridized the template over 5 minutes on a temperature ramp from 72 C to 38 C.

**Fig. 9.**
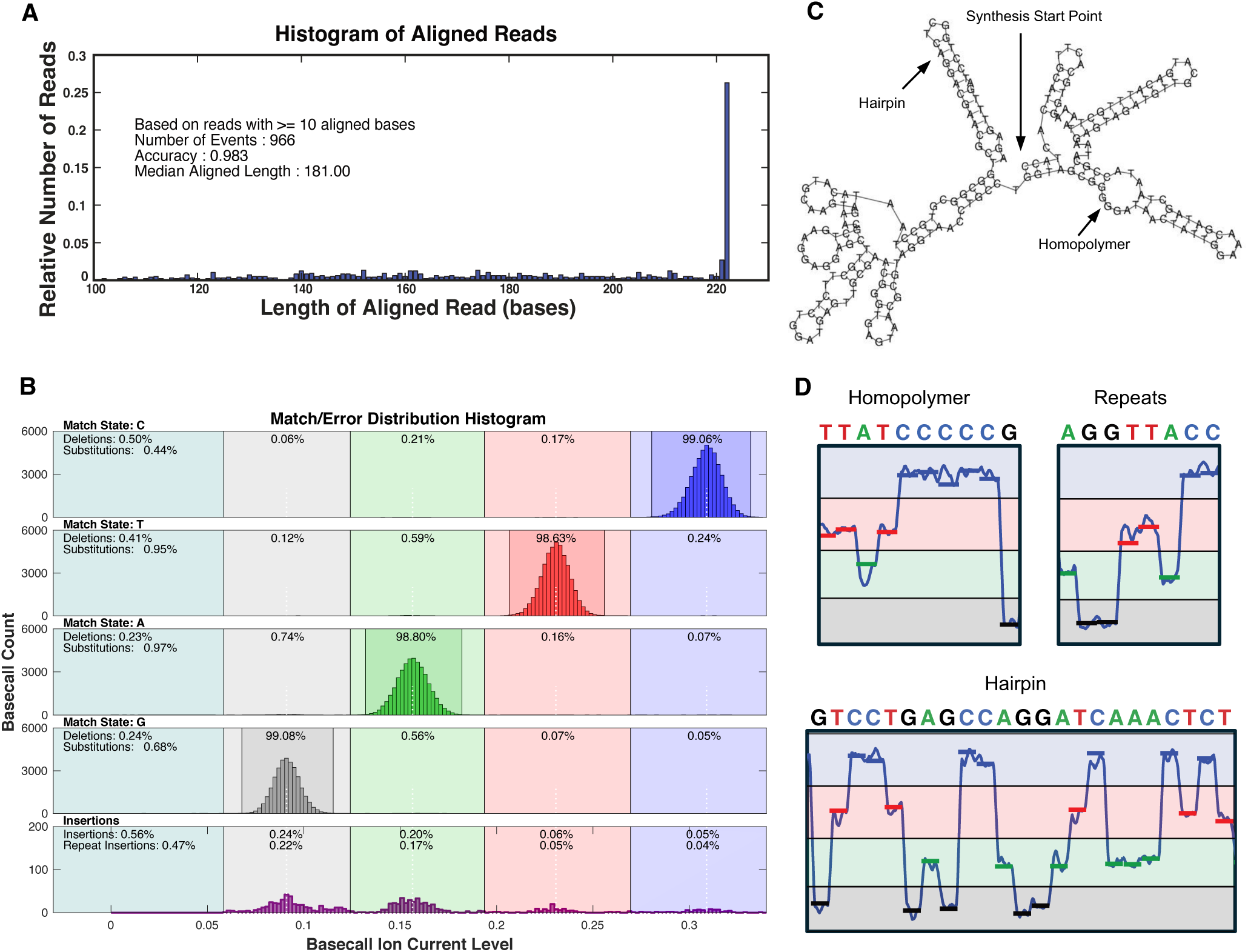
Xp measurements in a nanopore from 10-minute run. Ion current measurements, captured during each 1.0 ms period that a reporter is in the nanopore, were normalized to the open channel ion current and reduced to produce the basecall ion current level. Basecalls were assigned to these levels according to the ion current range they fall into. The basecalled reads were then aligned to the 222 base DNA template using the Smith-Waterman algorithm. **(A)** The distribution of 966 aligned Xp read lengths longer than 10 bases showed 26% of the reads are full length and have a median aligned read length of 181 bases. **(B)** These histograms present the basecall matches and error distributions of the 966 read set. Basecall ion current level is plotted along the abscissa with basecall call bin counts shown on the ordinate using bin widths of 0.002. The vertical demarcation lines define the ion current ranges, that are used to assign basecalls for <u>C</u>ytosine, <u>T</u>hymine, <u>A</u>denine, and <u>G</u>uanine ion current levels (in descending value order). The upper four histograms show sequence match and substitution distributions for each of these. Count distributions of matched basecalls for each basetype appear in the range containing the dominant peaks. Base substitutions appear in the three adjacent basecall ranges. Deletions are summarized with total base substitutions in the leftmost column. For each basetype, the percentage matches, substitutions and deletions are normalized to the total base count of that basetype found in the template portions of the aligned reads. As an example, in the top histogram for the cytosine basetype, 99.06% were matched, 0.44% were substitutions (G: 0.06%, A: 0.21%, T: 0.17%) and the remaining 0.50% were deletions. To illustrate the well differentiated signal-to-noise quality of this data, the four subranges shown with darker contrast bound 99.7% of the matches for their respective basetype. The insertion distribution (rescaled by 30x to visualize) is located in the bottom histogram. Insertion percentages (example: C insertions = 0.05%) are normalized to the total number of bases in the template portions of the aligned reads. Listed beneath the insertion percentages are the repeat insertions, a subset of insertions that have an adjacent base of the same basetype as discussed in Sequencing Fidelity. **(C)** Secondary structure of the 222-mer template, generated by the DNA Secondary Structure prediction tool on vectorbuilder.com. **(D)** Magnified regions of ion current traces demonstrating SBX sequencing of homopolymer, repeat, and hairpin regions.

We initiated Xp synthesis by the addition of SBX Extension Master Mix (see Supplement) and incubated the reaction for 1 h at 37 C, producing a heteroduplex Xp-DNA product that remained anchored on the flow channel. Extension was followed by acid cleavage of the P–N bonds on the Xp strand, allowing the Xp to expand into its sequence-ready structure while minimizing ancillary acid damage. After acid neutralization, the SSRT enhancer moieties were modified by succinic anhydride (SA), which partially neutralizes the positive charge of the polyamine enhancers that flank the nucleotide [Fig. 6C]. Finally, we photocleaved the Xp products from solid support and diluted them for sequencing.

For single pore Xp sequencing (and the associated nanopore instrument), we adapted prior approaches [Fig. S22] (*44*, *47*). Briefly, we inserted a single wild-type αHL nanopore in a DPhPE lipid bilayer, oriented with the barrel-side facing the *cis* reservoir using Insertion/Trans Buffer in both reservoirs. We qualified the correct pore orientation by conductance analysis and then replaced the buffer in the *cis* reservoir by perfusion with 3 mL of Sequencing Buffer, leaving 80 μL in the reservoir. Salt content in the *cis* and *trans* reservoirs during measurement were 0.4 M NH_4_Cl, 0.6 M GuHCl and 2 M NH_4_Cl respectively, all buffered to pH 7.4. The sample was sequenced for 10 minutes at 20 C.

We captured ion current measurements and applied potentials with patch-clamp instrumentation using a digital interface and a LabVIEW program for control [Fig. S22] (*48*). The potential applied to the *cis* reservoir was –70 mV DC baseline with –600 mV pulses (6 µs duration) applied at 1 kHz [Fig. 4D]. Amplifier voltage and current channel outputs were digitally recorded with 16-bit sampling from separate outputs of the Axopatch amplifier at 100 kSps and captured in a raw data file. The current channel was filtered by the amplifier at 10 kHz.

A MATLAB script was used to parse and translate the basecall in the raw data file from the 10-minute sequencing run [Fig. S23A], beginning with removal of the high voltage pulses from the trace [Fig. S23B-C]. Next, we computationally demarcated ion current ranges for each reporter using the four distinct peaks that emerged from analyzing the ion current data from the aggregate translocation events [Fig. S23D]. Each 1.0 ms reporter measurement was assigned a base identity according to the range in which its ion current fell. An example ion current trace of the full length 222 base *Streptococcus* DNA sequence is shown in Fig. 1E, S24 and the complete 10-minute run trace is in Fig. 10A-C.

**Fig. 10.**
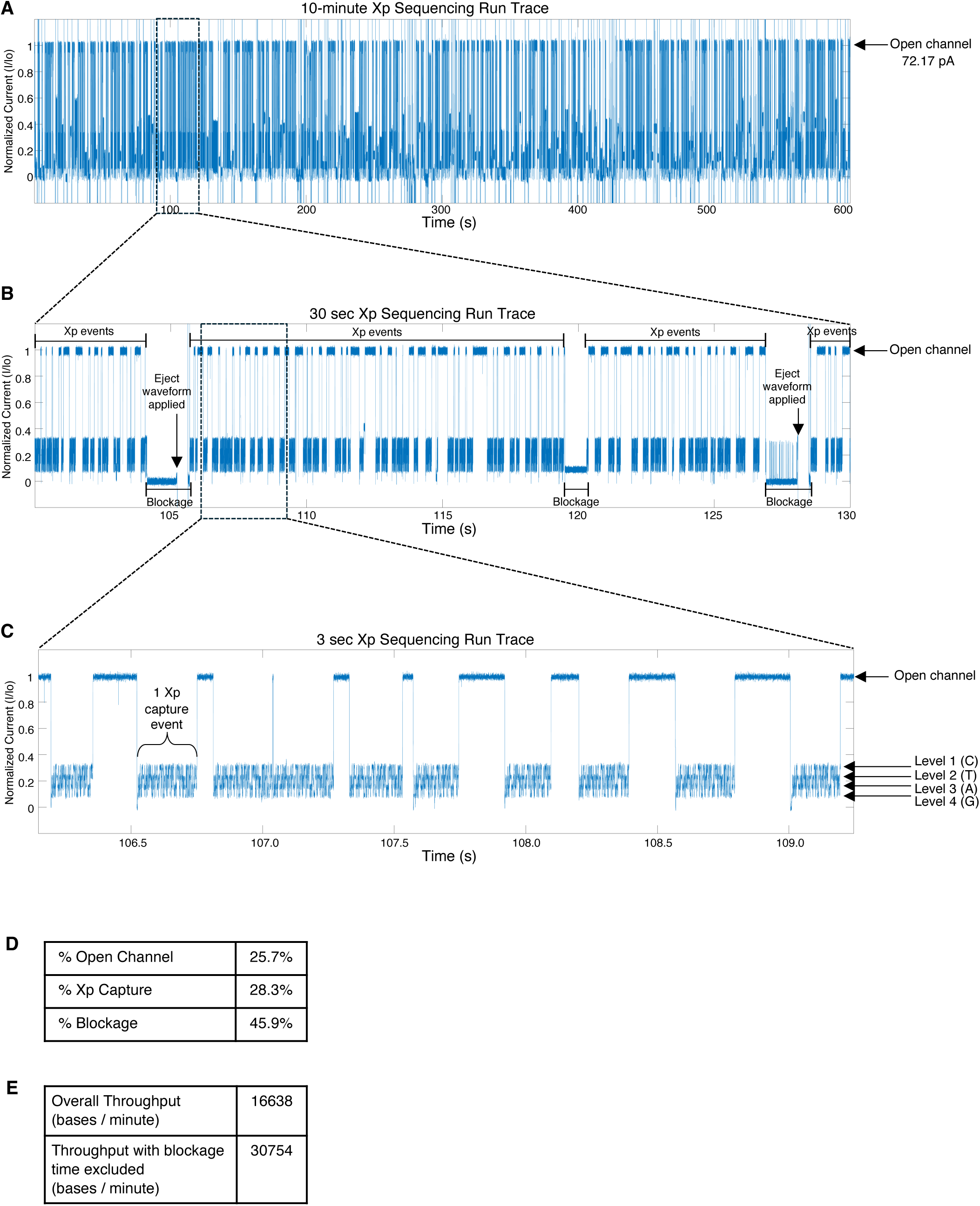
**Run overview and current trace data**. **(A)** Example of a full Xp sequencing run. Current trace represented as normalized current (I/I_0_). **(B)** The 30 s window shows clean Xp translocation events, occasional blockages and a few short-lived dysfunctional states. An eject waveform application was often required for removing blockages from the pore to resume an active capturing state. The eject waveform reversed the polarity of the voltage to push out material that was blocking the pore. **(C)** The 3 s window showcases the high capture efficiency of Xp molecules. **(D)** Relative time spent in various nanopore states. **(E)** Blockage states dominate nanopore measurement time, offering an opportunity to improve throughput.

The read rate of 1 kbase/s during an Xp translocation corresponded to the 1 kHz pulse rate. During the run, the nanopore was translocating Xp 28% of the time, and was waiting for Xp capture (open channel) 26% of the time [Fig. 10D]. The nanopore was in a blocked state 46% of the time (207 blocking events), which limited the possible throughput and yield of the run [Fig. 10E]. This occurred when the nanopore failed to return to the open channel condition. Blockages were resolved by temporarily reversing the voltage across the nanopore to eject a stuck Xp molecule from the nanopore before proceeding with the run. This 10-minute run recorded the measurement of 166,383 base reporters, for an average throughput of 280 bases/s.

We assessed the errors by aligning the reads with the DNA target using a Smith-Waterman algorithm (*49*), which provided aligned read lengths as well as insertion, deletion and substitution error counts. The run recorded 966 reads equal to or longer than 10 base calls that aligned to the target DNA sequence. Fig. 9A shows a histogram of this aligned read length distribution. Of these, 26% aligned to the full 222 base template length. Median read length was 181 bases. Smith-Waterman alignment trimmed an average of two unaligned bases from each end of the reads (see Supplement).

Mean accuracy of all reads longer than 10 bases in the run was 98.3% with errors arising from insertions, deletions and substitutions at 0.56%, 0.36% and 0.76%, respectively. Base call and error profile histograms are shown in Fig. 9B. Accuracy is defined here by normalizing the alignment errors with the number of template bases to which reads were aligned.

## Discussion

During the development of SBX, important observations and considerations were made that warrant further discussion. Foremost, the interdependence among the novel molecules in the context of the synthesis and measurement environments made SBX development a co-evolutionary process. For example, the Xp Synthase, PEM and Enhancer are all critical to produce high yields of full-length Xp, and removal of any of these three components results in synthesis failure [Fig. 7A-B]. Second, the modularity of the molecular components enabled a systems engineering approach, particularly for the PEMs, SSRT and EO, that accelerated understanding of these components and the technology in whole.

## Sequencing Fidelity

SBX sequencing errors occur during both Xp synthesis and nanopore measurement. Errors in Xp synthesis are influenced by many aspects of the reaction. Reaction temperature, order of addition, salt concentration, effective Mn^2+^ concentration, PEM composition, solvents and other reaction additives can all modulate Xp Synthase accuracy [Fig. S20F-I]. The reaction mixture that offers the highest synthesis accuracy likely balances the optimal reaction kinetics with control over solvation near the nascent base-pair, encouraging Watson-Crick geometry over the alternate configurations that are permitted by the combination of Mn^2+^ as the divalent cofactor and the solvent-exposed catalytic site of Dpo4 (*37*, *41*). Xp Synthase mutagenesis and PEMs design have led to significant improvements, and they continue to be a development focus. While the Xp Synthase errors are primarily base substitutions, indel errors also occur. The parent polymerase, Dpo4, is known to introduce deletions via a template slippage mechanism, so it is anticipated that the enzyme will contribute to the deletion rate (*50*).

Sequencing errors during nanopore measurement have multiple sources. The TCE reliably translocates the Xp by a single reporter during each pulse, but some errors do occur. In this experiment, a majority of insertions (0.47% of the total 0.56%) were repeats of their previous basecall. This insertion repeat is characteristic of a TCE insertion error. Experiments exploring pulse optimization show how small pulse voltage adjustments from optimum will drive the deletion and insertion rates in opposite directions. Small pulse force differences between the four base reporters exhibit distinctive deletion-insertion tradeoffs. For example, C and T have the highest deletions [Fig. 9B], but exhibit the lowest insertions in contrast with those of A and G. In an ongoing optimization effort, a matrix of TCE designs with associated reporters have been tested by nanopore sequencing using different reagent and pulse conditions to minimize these errors.

Substitution errors due to a reporter ion current level falling outside its correct range have been reduced through reporter design optimization by altering their size, composition, charge and position [Fig. 5A]. By avoiding the broad or bistable ion current responses exhibited by many TCE-reporter combinations and selecting for a balanced separation across the dynamic range of the ion current measurement, the selected optimized set of reporters exhibits robust, high signal-to-noise responses [Fig. 1G].

The polyamine portion of the enhancer region imparts positive charge to both arms of the tether. This localized charge density affects smooth translocation of the Xp that may cause measurement errors. To mitigate this impact, the more reactive amines of the enhancer region are modified with SA after Xp synthesis to neutralize some of the positive charge [Fig. 6C]. This modification chemistry is tunable, and control over the degree of succinylation can be used to optimize measurement accuracy. Insufficient SA modification results in the polyamine portion retaining too much positive charge, leading to an increase of insertion errors or, in some cases, erratic bistable states and blockages. Over-modification of the amines with SA causes overly fast translocation that results in an increase of deletion errors.

Other sources of noisy Xp nanopore measurement have been identified and addressed during development with minor error contributions relative to those discussed above. Noise and error contributions from steric blockages, pore gating (*51*), environmental noise and co-translocation of multiple Xp have been minimized by effective identification and filtering. Blockages are a common occurrence which are resolved during measurement by briefly reversing voltage to back the Xp out of the pore [Fig. 10B]. Examples of these blocked state ion current traces are shown in Fig. S25A-E. Barrel-first entry of the hemolysin nanopore avoids the congestion of the Xp in the nanopore vestibule that causes erratic deep blockage of the ion current [Fig. S21C].

## Near Real-Time Sequencing

A key advantage of SBX is the potential for massively parallel measurements with high accuracy and speed. Such highly parallelized measurements could produce independent streams of data with events that are highly localized in time. Under such a scenario, sufficiently fast computational hardware could execute inference routines at low latency, allowing each read to be called from its corresponding time series data, which can then be further processed shortly thereafter. The short duration and independence of Xp translocation events could allow for such processing to occur even while data acquisition continues over the course of a run. With such immediate processing after measurement, near real-time results can be generated to provide the user with flexibility to end the run when sequence requirements are satisfied. Additionally, run times can be adjusted to match output needs, enabling, for example, small batch multiplexing. By contrast, sequencing-by-synthesis (SBS) based technologies employ long standard cycle times that run to completion before information is available to call a complete read. Access to early result trends and data quality assessment are ancillary user benefits that increase efficiency and flexibility.

## Scalability

The high signal-to-noise performance of SBX reduces the need for sophisticated amplifiers and shrinks the footprint requirement of nanopore cells for CMOS-based nanopore array design (*52*). This anticipates scaling to array sizes of 8 million nanopores (*53*, *54*). In these designs, CMOS amplifier arrays are post-processed to add an electrode in a small volume cell above each array amplifier. Prior to operation, the cells can be filled with electrolyte and coated with a lipid bilayer that is a barrier to ion transport. Next, a biological nanopore can be inserted into each cell’s bilayer. All of the array cells are fluidically connected through their respective nanopores to a shared flow channel reservoir containing a shared counter-electrode. During operation, the CMOS is designed to rapidly and periodically sample the nanopore impedance of each cell. Leveraging efficient inference routines implemented on accelerated compute hardware, the raw signals produced by Xp measurement events can be translated into base calls (primary analysis). The resulting reads may then immediately be routed for post-primary processing. By adapting Xp measurement to these arrays, it is reasonable to consider that the single pore Xp sequencing rate of 280 bases/s demonstrated here can be scaled to a throughput exceeding 500 million bases/s.

## Comparison to On-Market Technology

A natural comparator for SBX would be ONT. It is challenging to find a study to compare this SBX experiment with state-of-the-art ONT sequencing given the scope and the data filtering that is applied in most ONT studies. Reports of average ‘raw’ accuracies cover a broad range (*8*, *55*, *56*) that depend heavily on the trained basecaller version (e.g. Guppy fast, high accurate (hac) and super accurate (sup) versions) with little detail of the quality cutoffs used. A recent study evaluated the MinION device using the R10.4 flow cell and SQK-LSK112 sequencing kit to sequence HCC78 genomic DNA (*8*). It compared the hac and sup basecallers aligned to the source genome using data filtering that included eliminating reads <200 bases, applying minimal MapQ score 60, and removal of indels with length <100 bases. The reported observed accuracies were 95.6% and 96.8% for the hac and sup basecallers, respectively. By comparison, this SBX experiment used a simple four-range classifier to make millisecond synchronized reporter basecalls followed by alignment and elimination of reads <10 bases to yield an average accuracy of 98.3%. This accuracy is even more significant given the high measurement throughput. In the same study, the reported MinION yield was 12.85 Gb in a 72-hour run or about 50,000 bases/s. The example of SBX scaled to an 8 M pore array would reasonably be >10,000 times that throughput and would also exceed the yields observed for state-of-the-art sequencing by synthesis (SBS) technologies (*57*).

This improved performance, simplicity of measurement and computational reduction clearly demonstrates the signal-to-noise benefits of SBX. The data presented here were captured in January of 2020 using an early chemistry that demonstrates all the critical functionality of the SBX technology and is a prelude to newer chemistry now being applied to large CMOS arrays (*47*, *54*, *58*).

## Conclusions

Specialized molecular constructs, chemistries and methods developed for SBX were utilized to demonstrate this novel single-molecule sequencing technology. This approach solves the signal-to-noise challenges of direct sequencing by converting DNA into an expanded, more measurable structure. Key innovations were required to make this technical achievement possible including the development of:

– XNTPs, nucleoside triphosphates modified with structures engineered to control translocation, expand, provide well-resolved nanopore based reporter codes and enable template-dependent incorporation with a modified polymerase (Xp Synthase).
– PEMs, a class of critical cofactors that enable polymerase extension of XNTPs beyond 10 bases by stabilizing the elongating heteroduplex structure.
– Xp Synthases, highly mutated DNA polymerases that processively and accurately incorporate XNTPs to synthesize Xpandomers.
– Reaction formulations and processes, optimized for accurate, template-dependent synthesis of Xp using a solid-phase workflow.
– Xp end structures and sequencing conditions, engineered to efficiently and accurately thread Xp molecules though a nanopore.

The single-molecule results shown provide an opportunity to contemplate a scaled implementation of SBX. As introduced here, single-nanopore SBX was demonstrated with accurate, well-resolved reporter sequencing that can be adapted to high-throughput parallel CMOS-based architectures. Based on Xp sequencing rates of 280 bases/s shown for single pores, throughputs of greater than 500 million bases per second can be reasonably considered. Furthermore, separated Xp synthesis and sequencing workflows allow each process to be independently optimized and efficiently performed. In contrast to current SBS methods (*59*), near real-time SBX data output at high-throughput would allow for flexible operation across a range of experimental scales. Such flexibility would reduce batch requirements, enable faster time to result and ultimately allow users to rethink how sequencing workflows can be implemented for existing and emerging applications.

## Supporting information

SBX Supplement

## Acknowledgements

We thank A. Stephan, H.F. Johnson, J. Steel, S. Rose, L. Huntsman, H Kawaguchi and H. Dreismann for program support; L. Loeb, A. Marziali, V. Makhijani, D. Heindl, H. Kuchelmeister, J. Mannion, D. McAndrew, A. Wallace, M. St. Louis, M. Lahman, A. Asbe, A. Holt, E. Quig, E. Hermanns, J. Loo, R. Walsh, K. McElligott, C. Berrios, A. Lee, M. Carroll, J. Horsman, E. Williams, N. Westergreen, T. Frese, C. Rosevear, Y. Umezawa, K. Fosnacht, K. Smith and J. Chang for technical support; K. Walker for patent support; B. Poppell for IT support; D. Jones for finance support; B. Anderson for 3D animation; A. Regev for publishing support. Funding for this work was provided by Stratos Genomics Inc., grants from the National Human Genome Research Institute (R21 HG006304) and Department of Defense Small Business Innovation Research (DARPA-BAA-09-22 Strategic Technologies). Funding was acquired by A. Stephan, M. Kokoris and R. McRuer. M. Kokoris and R. McRuer conceptualized the program. M. Kokoris is the first author and PI. All authors participated in the research and investigation. RM developed the software. MK, RM, MN, AJ, MP, CC, TL, CM, JT, SV, KC, supervised the research. MK, RM, MN, AJ, MP, CC, KB, TL wrote the original draft. MK, RM, MN, AJ, MP, CC, KB, LM, CW, TL reviewed and edited. Competing interests: Various authors are inventors on the following patents 7939259, 8349565, 8324360, 9920386, 9670526, 10851405, 11920184, 10457979, 10676782, 10301345, 10774105, 10745685, 11299725, 12037577, 12116570, 11530392, 11970731, 9771614, 8586301 and 10996213 held by Roche Sequencing Solutions, Inc. that cover SBX Technology. MK, RM, MN, AJ, MP, CC, KB, TL, JC, LM, ML, TR, CW, SB, AL, MK, RB, SM, BB, BK, ML, MW, AK, AA, MRP, SK, JL, MB are employees of Roche Sequencing Solutions, Inc. SBX, Xpandomer and XNTP are registered trademarks of Roche Sequencing Solutions, Inc.

