## Supplementary material for "Sequencing by Expansion (SBX) – a novel, high-throughput single-molecule sequencing technology": SBX Supplement

**Mark Kokoris *et al.***

**The PDF file includes:**

**Materials and Methods**
**Figs. S1 to S25**
**Table S1**

#### Materials and Methods

##### Synthesis of XNTP components

*Nucleoside synthesis*

Bis-alkyne nucleoside triphosphates were synthesized on solid support [Fig. S14A], undergoing multiple transformations before cleavage and purification. Propylamine derivatized 500 Å controlled-pore glass support (LGC Biosearch) was functionalized with hydroquinone-O,O'-diacetic acid (Alpha Aesar) to synthesize Q-support (*1*). 5'-ODMT 7-deaza-7-octadiynyl-N6-Pac 2'-deoxyadenosine, 5'-ODMT 7-deaza-7-octadiynyl-N2-Pac 2'-deoxyguanosine, 5'-ODMT 5-octadiynyl 2'-deoxycytidine and 5'-ODMT 5-octadiynyl 2'-deoxyuridine were purchased (Chembiotech) or synthesized through a Sonogashira coupling to the respective iodide (*2*). Nucleosides (160 µmol/gram support) were individually loaded on to Q-support through HBTU (NovaBiochem) (2 equiv.) / DMAP (Tokyo Chemical Industry) (4 equiv.)-mediated coupling for 16-36 hours. Typical loadings of 80 µmols nucleoside on 1 gram of support were obtained, and any uncoupled nucleoside was recovered. The support was washed with MeCN, MeOH, and DCM (Millipore Sigma) followed by a capping step with standard oligosynthesis reagents. Nucleosides were detritylated by addition of deblock agent (3% dichloroacetic acid in DCM) (Glen Research), then converted to the 5'-iodide by treatment with methyltriphenoxyphosphonium iodide (5 mmol) in DMF (Millipore Sigma) for 1 hour (*3*). After washing the support with DMF, MeCN, and DCM, the 5'-iodide was converted to the 5'-azide by treatment with a solution of NaI (1 mmol) and $NaN_3$ (1 mmol) (Millipore Sigma) in DMF for at least 18 hours. The 5'-azide was washed with DMF, MeCN and DCM, and was stable on support for up to a year at -20 C.

*Phosphoramidate synthesis*

The 5'-phosphoramidate was synthesized through a Staudinger reaction, where the aza-ylide undergoes a Michaelis-Arbuzov reaction (*4*, *5*). Cyanoethyl hexynyl methyl phosphite was synthesized by reacting hexynyl phosphoramidite (Glen Research) with methanol and ETT activator (Glen Research). For C nucleoside, this was replaced with the methyl phosphite of DMT-C6-spacer phosphoramidite (Glen Research). The phosphite (12 equivalents in DMSO) was added to the support-bound 5'-azidonucleoside along with LiCl (Acros Organics) (75 equivalents in pyridine (Millipore Sigma)) and the reaction was incubated for 36 h. After washing with DMF, MeCN, water (Millipore Sigma), and DCM, the support bears the CNEt protected mono-phosphoramidate. The 5'-phosphoramidate dC analogue utilized the DMT-C6-phosphite in place of the hexynyl. Beta and gamma phosphates were added through iterative phosphorylation on an automated oligonucleotide synthesis platform. Cyanoethyl protection was removed with a solution of DBU (Tokyo Chemical Industry) and Bis(TMS)acetamide (Acros Organics). ETT activator was used to add a single Bis-CNEt phosphoramidite (Chemgenes), followed by *t*-BuOOH (Millipore Sigma) oxidation. Another round of deprotection of the CNEt was followed by another addition of Bis-CNEt phosphoramidite and oxidation. If necessary for the dCTP analogue, detritylation was performed with subsequent addition of hexynyl phosphoramidite. After the final oxidation and removal of the remaining CNEt, dNTPs were cleaved off Q-support with ammonia (Millipore Sigma) for up to 15 minutes, followed by evaporation under reduced pressure. The triphosphate was purified by RP(C18)-HPLC (100 mM TEAA (Glen Research) in water/MeCN), which separated the two isomers at the alpha phosphoramidate [Fig. S14B]. Only one of these was active and used to synthesize XNTP.

*Phosphoramidite building blocks*

Pendant mPEG-4 phosphoramidite (PPA) was synthesized as described in Fig. S8A. (*R*)-(−)-2,2-Dimethyl-1,3-dioxolane-4-methanol (Tokyo Chemical Industry) was silylated in DCM, with DMAP (0.08 equiv.), TEA (1.5 equiv.) and TBDPS chloride (Millipore Sigma) (0.95 equiv.). The crude material was extracted from water and concentrated. Without purification, the foamy solid was deprotected in 67% acetic acid (Millipore Sigma), heated to 85 C, then concentrated and purified by flash chromatography. The diol was monotritylated in DCM with TEA (1.5 equiv.) and DMT Chloride (Chemgenes) (0.97 equiv.) which was added slowly. The reaction was extracted from water and concentrated, then purified by flash chromatography. Tosyl-mPEG4 was added by first cooling the trityl glyceride in THF (Millipore Sigma) in an ice bath, adding NaH (Millipore Sigma) (2 equiv.), then warming to RT for 10 minutes. The reaction was again cooled in the ice bath, and tosyl-mPEG4 was added. The alkylation was allowed to warm to RT slowly and was monitored by TLC for completion. The reaction was cooled on ice, quenched with water, then extracted with ethyl acetate (Millipore Sigma). The organic layer was concentrated, filtered, then purified by flash chromatography. The product was suspended in THF and desilylated with TBAF (Acros Organics) (2 equiv.), extracted from water with ethyl acetate, and purified by flash chromatography. Phosphitylation was performed in DCM with TEA (2 equiv.) and *N,N*-diisopropylamino cyanoethyl chlorophosphoramidite (Chemgenes) (0.9 equiv.). The reaction was concentrated and filtered to remove TEA-Cl, then purified by flash chromatography using hexanes and ethyl acetate containing 1% TEA.

Benzofuran phosphoramidite was synthesized as described in Fig. S8B. (*R*)-(+)-Glycidol (Tokyo Chemical Industry) was tritylated in DCM with TEA (2 equiv.) and DMT Chloride (0.95 equiv.) which was added slowly. The reaction was extracted from water and concentrated, then purified by flash chromatography. The heterocycle furo[3,2-c]pyridin-4(5h)-one (1.1 equiv.) was deprotonated with NaH (0.25 equiv.) in DMF for 2h at room temperature, after which a solution of the tritylated glycidol in DMF was added. The solution was incubated at 80 C overnight, then concentrated and purified by flash chromatography. Phosphitylation was performed in DCM with TEA (2 equiv.) and *N,N*-diisopropylamino cyanoethyl chlorophosphoramidite (0.9 equiv.). The reaction was concentrated and filtered to remove TEA-Cl, then purified by flash chromatography using hexanes (Millipore Sigma) and ethyl acetate containing 1% TEA.

*Symmetrically synthesized reporter tether (SSRT) synthesis*

SSRTs were prepared using standard phosphoramidite chemistry using a MerMade 12 oligonucleotide synthesizer (BioAutomation). CapA/B, oxidizer (20 mM $I_2$ in THF/Pyridine/Water), and activator (ETT) solutions were purchased from Glen Research. Commercial phosphoramidites were purchased from Glen Research and Chemgenes. A glossary of all phosphoramidites used is presented in Fig. S7. Synthesis was initiated on 1 µmol columns of 1000 Å CPG UnySupport (Glen Research), followed by the coupling cycles for the series of phosphoramidites shown in Fig. S9. The columns were washed with MeCN, DCM, DMF, then incubated with an azido modification solution (100 mM sodium azide and sodium iodide in DMF). The columns were rinsed with 10% diethylamine (Alfa Aesar) in MeCN, then cleaved from support using 30% ammonia for 1 hour, followed by addition of an equal volume of 40% methylamine (Millipore Sigma) and incubation for an additional hour. After additional incubation, the cleaved oligo was desalted by size exclusion chromatography and purified by RP-HPLC.

*Cyclizing click reaction*

SSRTs were clicked to bis-alkyne dNTPs at a ratio of 1:1.1, respectively. All reagents were from Millipore Sigma unless otherwise noted. The SSRT (0.27 µmol) and dNTP (0.3 µmol) were added to sodium phosphate (Teknova) buffered DMF (25 mM and 10% final, respectively). $MgCl_2$ (67.5 µmol), aminoguanidine (54 µmol) and sodium ascorbate (8.1 µmol) were added. A catalyst mix of THPTA (8.1 µmol), sodium ascorbate (8.1 µmol), copper sulfate (2.7 µmol) and aminoguanidine (13.5 µmol) was prepared in 10% DMF. The catalyst mix was added to the substrates, vortexed, and incubated statically for 30 minutes at RT. The reaction was quenched with EDTA (371 µmol) and placed on ice. The cyclized oligo was purified by RP(C8)-HPLC (100 mM TEAA in water/MeCN), then desalted by RP cartridge (Glen Research), concentrated to 0.25 mM and buffered to pH 7.25 with 25 mM Tris-acetate [Fig. S10C].

**Extension oligo Synthesis**

The EO contains four regions per the sequence in Fig. S15: the primer, concentrator, leader, and tether to the support. Synthesis of the EO was initiated on 1 µmol columns of 1000 Å CPG UnySupport, using standard phosphoramidite chemistry. The columns were washed with MeCN, DCM, DMF, then incubated with an azido modification solution (100 mM sodium azide and sodium iodide in DMF). The columns were rinsed with 10% diethylamine in MeCN, then cleaved from support using AMA for 30 minutes at 65 C. The oligos were protected from light and purified by PAGE. During use, photocleavage of the EO results in a 5'-phosphate.

**PEM synthesis**

*Combinatorial synthesis matrix*

After discovering a minor-groove binder that dramatically increased Xp extension, a large matrix of similar molecules was synthesized [Fig. S2A, B]. Solutions of bis-alkyne cores (1 µmol) and azide arms (2 µmol) were diluted to 50 µL with DMSO and added to a 96-well plate. An equal volume of catalyst mix containing copper sulfate (5 mM final concentration), TBTA (Tokyo Chemical Industry) (6 mM), and sodium ascorbate (20 mM) was added to each well and pipet mixed. The resulting solutions immediately showed color changes and many acquired fluorescence under 365 nm light. The consumption of starting material after 30 minutes was determined by TLC, and the clicked product appeared significantly more polar. PEMs were tested crude on a 45-mer template, diluting to 500 µM in the extension reaction (based on the alkyne core). Several molecules demonstrated a PEM effect, with the angle of the arms and the presence of the salicylic acid both correlating with improved extension. Hit molecules were resynthesized at 25 µmol scale, then purified by solid phase extraction with a C18 cartridge. A second round of testing was performed using a 100-mer template and varying the concentration of PEM.

*AZ-43,43 PEM synthesis*

The AZ core was synthesized as follows. A flask containing 2,6-dibromopyridine-4-carboxylic acid (AOB Chem) (14 g, 50 mmol) and HATU (Chem-Impex)(19 g, 50 mmol) in 62 mL DMF and 17.4 mL DIPEA (Acros Organics) was treated with aqueous ethylamine (Millipore Sigma) (13.6 mL, 125 mmol) and incubated at room temperature for 20 minutes. The reaction was monitored by TLC (1:1 hexanes:ethyl acetate) for complete consumption of starting acid. The reaction was extracted from water with DCM (3x) and the combined organic layers were concentrated. The crude product was separated by silica gel chromatography (0-40% ethyl acetate in hexanes) and dried to a white crystal (7.9 g, 25.6 mmol, 91% yield). This material was diluted in 50 mL DMF and TMS-acetylene (Tokyo Chemical Industry) (8.9 mL, 64 mmol) was added. DIPEA

(35 mL), $Pd(PPh_3)_2Cl_2$ (Tokyo Chemical Industry) (2.7 g, 3.8 mmol), and CuI (Acros Organics) (1.5 g, 7.7 mmol) were added. The reaction was stirred at room temperature for 2 hours, then was concentrated to a thick black oil. The crude product was loaded on a silica gel column in minimal DCM and was purified (0-40% ethyl acetate in hexanes) then dried to a white flakey solid (4.5 g, 13.3 mmol, 52% yield). This TMS-protected AZ core (3.4 g, 10 mmol) was dissolved in 63 mL ethanol (Koptec) and potassium carbonate (Thermo Scientific) (2.9 g, 21 mmol) was added. After 5 minutes, TLC showed complete deprotection of the alkynes. Azide 43 (4-azido-2-(trifluoromethyl) benzoic acid) (A1 Biochem) [Fig. S18A], 4.9 g, 21 mmol) was added and stirred for 10 minutes to completely dissolve. Sodium ascorbate (4.2 g, 21 mmol) was pre-dissolved in 7 mL water, and a solution of 1 M copper sulfate (5.5 mL, 5.5 mmol) was added. This mixture was immediately added to the reaction which was stirred vigorously for 1.5 hours. The dark amber solution was quenched with 0.5 M EDTA (50 mL, 25 mmol) and stirred for 2 hours until it turned deep blue. The crude reaction was concentrated and dissolved in minimal DMSO, then loaded on a C18-RP flash column (Buchi) (0-100% MeCN in water). The product was concentrated and repurified 1-2x to remove trace yellow contaminants. The final product was colorless and lyophilized to a fluffy solid (3.96 g, 6 mmol, 60% yield). The PEM was soluble in water (100 mM) and had a maximum absorption at 318 nm.

**Protein expression, purification, and screening**

*Oligonucleotides for Xp Synthase screening*

Primer extension experiments in the block lysate screen were carried out with a primer labeled with a 5' SIMA-HEX moiety (Glen Research) (W) using 2'-O-Me RNA for all non-T bases and DNA for T (Chemgenes).

Primer: 5′-WGAACCAGTATACTCCTCGATG-3′

Template:

5′–ACTGACTGACTGACTGACTGACTGACTGACTGACTGACTGACTGACTGACTGACTGACTGACTGACTGAC
TGACTGACTGACTGACTGACTGACTGACTGCATCGAGGAGTATACTGGTTC–3′

*Amino acid sequence of Xp Synthase*

Dpo4_C5275(Δ341-350)-His x6 amino acid sequence with mutated positions in bold:
MIVLFVDFDYFYAQVEEVLNPSLKGKPVVVCVFSGRFEDSGAVATANYEARKFGV**YS**G**M**PIV**R**AKKILPNAV
YLP**W**R**DL**VY**WG**VS**E**RIMNLLREYSEKIEIASIDEAYLDISDKVRDYREAYNLGLEIKNKILEKEKITVTVGI
SKNKVFA**AV**A**GR**MAKPNGIKVIDDEEVKRLIRELDIADV**Q**GI**PYF**TAEKLKKLGINKLVDTLSIEFDKLKGM
IGEAKAKYLISLARDEYNEPIRTRVRKSIGR**T**VTMKRNSRNLEEIKPYLFRAIEE**C**YYKLDKRIPKAIHVVA
**WKSYWNSQY**R**WSWF**PHGISKETAYSESVKLLQ**Q**IL**KK**D**K**RKIRRIGVRFSKFLEHHHHHH

*Nucleotide sequences of proteins in SBX*

Xp Synthase C-Terminal 6xHis:
5′–ATGATTGTTCTTTTCGTTGATTTTGACTACTTTTACGCTCAAGTTGAAGAAGTTTTAAATCCGTCTTTGA
AAGGAAAACCAGTTGTTGTTTGTGTATTTTCAGGGAGATTTGAGGATAGCGGTGCCGTGGCTACTGCAAACT
ATGAAGCTAGAAAATTTGGAGTATATTCTGGAATGCCAATCGTTCGTGCTAAGAAAATATTACCTAATGCAG
TTTACTTACCCTGGAGAGATCTTGTATATTGGGGTGTTTCCGAGAGAATAATGAACTTACTAAGAGAATACT
CCGAGAAGATCGAGATTGCAAGTATAGATGAGGCTTATCTTGATATCTCAGACAAAGTCAGAGATTATAGAG

AGGCATATAATCTAGGTTTGGAGATTAAGAACAAAATACTTGAAAAAGAGAAAATTACAGTTACTGTAGGGA
TTTCCAAGAATAAGGTATTTGCGGCTGTTGCTGGGCGTATGGCAAAGCCAAATGGAATAAAAGTTATTGATG
ATGAAGAAGTTAAAAGATTAATAAGAGAGCTAGATATAGCGGATGTACAGGGAATACCTTATTTCACTGCGG
AGAAACTAAAGAAGTTAGGTATTAACAAGCTAGTTGATACGTTAAGCATTGAATTTGATAAACTAAAGGGAA
TGATAGGCGAAGCTAAGGCTAAATATTTGATCTCTCTAGCTAGAGACGAGTATAACGAGCCTATAAGAACTA
GAGTACGAAAGAGTATTGGGAGAACTGTAACGATGAAGAGAAATAGCAGGAATCTGGAGGAAATAAAACCGT
ATTTATTTAGAGCAATAGAAGAATGCTATTATAAGTTAGATAAGAGGATTCCTAAAGCTATTCACGTAGTCG
CATGGAAAAGTTATTGGAATTCGCAGTATCGTTGGAGTTGGTTCCCTCATGGAATAAGTAAGGAAACTGCAT
ATAGTGAATCAGTAAAATTATTACAGCAGATATTGAAGAAGGATAAGAGAAAGATAAGAAGAATCGGAGTAA
GGTTCAGTAAATTTCTCGAGCACCACCACCACCACCACTGA–3'

KOD SSB N-Terminal 6xHis:
5′–ATGCACCACCATCATCACCATATGGAGGTCCTTACTAAAGACGAAATCATAAACCGCATAATTCGGGAAA
GAGGACTTTCTCGTAGTGAAATCGAAGAAAAAATCCGTGAGTTAGCGAAAATGCATGGTGTTAGTGAAAATG
CTGCGGCAGTTATGCTTGCTGAAGAGCTTGGAGTGTCTCTTGGTAAGGAAGAGGAAATGCTTTATATCAAAG
ACTTGGTGCCCGGTATGACTGGTGTAAATATCGTTGCGCGCATCAAACGTAAGTTTCCACCTAGAGAATATA
CACGTAGAGATGGATCAACCGGACGGGTCGCCGATCTTATCATATATGACTCAACAGGACAGGCGCGTCTTG
TGCTCTGGGACGCCATGGTGGCGAAGTATTATGACGATTTGAATGTGGGAGATGTCATCAAAGTCATCGACC
CAACCGTAAAAGAAGGCATGCGGGGTGTGGAATTGCATGCGAATTTTCGGACCCGCATTATAAAAAATCCAG
AGGATCCACGCGTAGAAGAAATTCCACCTCTTGAGGAAGTCCGTTCCTATAACTATCGTCGCGTCCAGATCA
AAGAGTTGCAGGGAGGCGAACGCTTCGTCGAGGTGAGAGGAACAATAGCCAAGCTTTATCGGGTTTTGGTGT
ATGATGCGTGTCCTGAATGTCGCAGACGCGTCGATTACGATCCTTCAACTGATACTTGGATCTGTCCAGAGC
ATGGCCCCGTGAATCCTGTGAAAATCACTGTACTTGATTTTGGCTTGGATGACTCAACAGGATACATACGCA
CGACACTGTTCGGCGACAGTGCGGCAGAGCTCATAGGGGAGGAACCAGAGGTAATCGACGAGAAACTTAAAA
AATTGATAGACGAAGGACTCACACCAAAAGAGGCTGGAAAAAGACTTGCGGAAGACGAGTATTACCCCTTAA
TAGGAAAGGAAATCGTAGTACGCGGATCCGTCGTAGAAGACAAATTCTTGGGCACTCTCTTTAAGGCCAGAT
CCTGGGATGAGGTAAATGAGAAAGCGGAAATAGAAAGAGTACGCAGAGAACTCTATAGAGAACTCAAAGAGT
ACGGATTGGAATGA–3'

**α**-HL C-terminal 6xHis with TEV cleavage:
5′–ATGGCAGATTCTGATATTAATATTAAAACCGGTACTACAGATATTGGAAGCAATACTACAGTAAAAACAG
GTGATTTAGTCACTTATGATAAAGAAAATGGCATGCACAAAAAAGTATTTTATAGTTTTATCGATGATAAAA
ATCACAATAAAAAACTGCTAGTTATTAGAACGAAAGGTACCATTGCTGGTCAATATAGAGTTTATAGCGAAG
AAGGTGCTAACAAAAGTGGTTTAGCCTGGCCTTCAGCCTTTAAGGTACAGTTGCAACTACCTGATAATGAAG
TAGCTCAAATATCTGATTACTATCCAAGAAATTCGATTGATACAAAAGAGTATATGAGTACTTTAACTTATG
GATTCAACGGTAATGTTACTGGTGATGATACAGGAAAAATTGGCGGCCTTATTGGTGCAAATGTTTCGATTG
GTCATACACTGAAATATGTTCAACCTGATTTCAAAACAATTTTAGAGAGCCCAACTGATAAAAAAGTAGGCT
GGAAAGTGATATTTAACAATATGGTGAATCAAAATTGGGGACCATATGATAGAGATTCTTGGAACCCGGTAT
ATGGCAATCAACTTTTCATGAAAACTAGAAATGGTTCTATGAAAGCAGCAGATAACTTCCTTGATCCTAACA
AAGCAAGTTCTCTATTATCTTCAGGGTTTTCACCAGACTTCGCTACAGTTATTACTATGGATAGAAAAGCAT
CCAAACAACAAACAAATATAGATGTAATATACGAACGAGTTCGTGATGATTACCAATTGCATTGGACTTCAA
CAAATTGGAAAGGTACCAATACTAAAGATAAATGGACAGATCGTTCTTCAGAAAGATATAAAATCGATTGGG
AAAAAGAAGAAATGACAAATGAAAACCTGTACTTTCAGGGACTCGAGCACCACCACCACCACCACTGA–3'

*Expression of Dpo4, KOD SSB, and alpha-hemolysin (α-HL)*

Wild type Dpo4 (residues 1-340), KOD SSB, or *S. aureus* α-HL were cloned into pET26b(+) expression vector (Millipore Sigma), in-frame with a C-terminal histidine tag, and underwent similar expression and purification workflows. Most of the 485 libraries used to introduce mutations into wild type Dpo4 or subsequent variant parent molecules were made with mutagenic primers for use in the Q5 Site-directed Mutagenesis kit (New England Biolabs). Similarly, the majority of variant designed clones were also constructed with mutagenic primers (Integrated DNA Technologies), and more complex designed clones were made using gBlocks gene fragments (Integrated DNA Technologies). Plasmids were transformed into T7 Express LysY electrocompetent cells (New England Biolabs), which were subsequently plated onto LB kanamycin agar plates (50 µg/mL) (Teknova). Fresh colonies were picked into overnight primary culture of LB broth (Corning) and 50 µg/mL kanamycin (Teknova), and then diluted 1:100 into 100 mL Super Broth (Teknova) and kanamycin. Flask cultures were grown, shaking at 37 C and 175 RPM to an optical density of 0.6-0.8, and then isopropyl β D-1 thiogalactopyranoside (IPTG) (Millipore Sigma) was added to a final concentration of 0.5 mM. Induction growth proceeded for 12-14 h at 25 C, 175 RPM. Cultures were chilled on ice, then pelleted by centrifugation at 4 C and 4000 x g, before storage at -20 C.

*Purification of polymerase and SSB*

All reagents were from Millipore Sigma unless otherwise noted. Frozen pellets were resuspended on ice in lysis solution containing 20 mM Tris-acetate (Thermo Scientific), 50 mM $NH_4OAc$, 1 mM EDTA, 0.1% Triton X-100, 5 mM imidazole, pH 8.3. A cocktail of ReadyLyse (Lucigen) and PMSF was added to fully resuspended pellets at 0.0025% and 0.5 mM final concentrations, respectively. Lysis proceeded for 30 minutes, followed by mechanical shearing of DNA achieved by 4 passes through a 21-ga needle. Lysates were incubated at 50 C for 15 minutes to inactivate host cell proteins. Streptomycin sulfate (0.5% final) was added dropwise to lysates while vortexing at 1000 rpm until chromatin had visibly condensed, about 45 seconds. Crude lysates were cleared by centrifugation at 20,000 x g for 15 minutes at 4 C. Cleared lysates were equilibrated for binding to HisPur Ni-NTA resin (Thermo Scientific) by adding EQ45 solution (20 mM Tris base (JT Baker), 45 mM imidazole, pH 7.5) to a final imidazole concentration of 10 mM. Ni-NTA resin that had been washed twice in EQ45 was combined with the lysate, and was incubated for 15 minutes, rotating at 15 RPM on a Rotator Genie. The protein-bound resin was rinsed twice with Wash buffer containing 20 mM Tris-acetate, 50 mM imidazole, 0.01% Triton X-100, 1 M NaCl (pH 7.5), before harvesting in Elution buffer containing 20 mM Tris-acetate, 50 mM imidazole, 0.01% Triton X-100, 1000 mM NaCl (pH 7.5). The eluates were desalted on PD Midi columns (Cytiva) into 0.2x Storage buffer containing 10 mM Tris pH 8.0, 10% glycerol (Teknova), 50 mM NaCl. Finally, the concentrated samples were concentrated 5x in a SpeedVac under no heat. Polymerase and SSB preparations were stored at -20 C.

*Synthesis of dNTP-2c Acetamide*

CPG-Q support bearing bis-alkyne nucleoside was transformed into dNTP-2c acetamide [Fig. S19C] by clicking to methyl azidoacetate and cleavage in ammonia. All reagents were purchased from Sigma. A 250 µL solution of 80% DMSO with 10 mM TBTA, 5 mM $CuSO_4$, 1% Triton X-100, 50 mM sodium ascorbate and 50 mM TEAA pH 7 was applied to CPG support containing approximately 700 nmol of bis-alkyne dNTP. This pretreatment was incubated for 20 minutes at RT in an 850 RPM orbital shaker. The tube was spun to pellet the CPG and the pretreatment was removed by pipet. A reaction mixture of the same composition, but with the addition of 140 mM methyl azidoacetate was added to the support and incubated for 1 h, RT, 850

RPM. The support was transferred to a fritted column on a vacuum manifold and the reaction mixture was removed. The support was washed (3x 400 µL) with a solution of 10 mM EDTA in 37% DMSO, 13% t-BuOH. The support was then washed with 10 mM EDTA, 50 mM $MgCl_2$, water, and MeCN (2 mL each). The support was dried with a 1 mL wash of DCM, continuing to pull vacuum until the CPG was free flowing. This material was cleaved off support with 1 mL 30% ammonia for 15 minutes at room temperature, then dried down in a SpeedVac to 100-200 µL without heat. The solution was separated from the CPG, which was washed (3x) with 500 µL 50% MeCN and pooled. The dNTP-2c acetamide was concentrated to remove MeCN before use, and could be purified by PAGE or RP(C18) HPLC.

*High-throughput block lysate screen I: Expression and lysis*

Plasmid libraries were transformed by electroporation into T7 Express LysY cells, and were plated onto LB agar with kanamycin (Teknova). Colonies were picked into 1 mL LB broth and 50 µg/mL kanamycin in 96 deep-well blocks, and primary cultures were grown 12-15 h shaking at 900 rpm at 37 C. Primary cultures were diluted 1:100 into another 1 mL deep-well block for expression culture. The primary culture block was centrifuged for 15 minutes at 4000 x g to pellet the cultures for subsequent freezing at -20 C to preserve cells for DNA miniprep. The expression cultures were grown for 3 h at 900 RPM, 37 C. IPTG was added to 1 mM final, and the cultures were grown for another 4 h. Cultures were pelleted by centrifugation, the media was aspirated, and the cell pellets were frozen at -80 C. Thawed pellets were resuspended in 200 µL lysis solution (20 mM Tris-acetate, 50 mM $NH_4OAc$, 1 mM EDTA, 0.1% Triton-X 100, 0.5 mM PMSF, 5 µg/µL Chicken Egg Lysozyme (Millipore Sigma), pH 8.3), with vortexing to assist in lysis. Blocks were incubated in a 56 C water bath for 7.5 minutes to denature host proteins. Streptomycin sulfate was added to 0.5% to each well, and the blocks were vortexed until visible chromatin condensation had occurred. An additional 600 µL of lysis solution was added to each well to dilute, and the blocks were centrifuged at 2300 x g for 10 minutes to pellet debris.

*High-throughput block lysate screen II: Primer extension*

To catalyze primer extension [Fig. S19C], 2 µL of protein lysate was added to 8 µL of a master mix containing 1.1 pmols primer (SGO-05698), 0.8 pmols template (SGO-07412), 10 mM Tris-acetate (8.3), 100 mM $NH_4OAc$, 100 pmols equimolar dNTP-acetamide nucleotides, and 2 mM $MnCl_2$ (Millipore Sigma) in a 384-well reaction plate. Reactions were incubated for 20 minutes at 37 C. Reactions were quenched by pipette mixing with 5 µL of a digest mix containing 0.5 mg/mL Proteinase K (New England Biolabs) and 0.0083% sodium dodecyl sulfate (SDS) (VWR), and the plates were heated to 85 C for 2 minutes. 2 µL of each reaction was added to 18 µL of formamide (Applied Biosystems) with mixing, and these dilutions were heated to 85 C for 2 minutes. Finally, samples were loaded onto a 3730 96-capillary DNA analyzer (Applied Biosystems) to assess length of extension products.

*Streptococcus 222mer Xp template Synthesis*

The Xp synthesis template is a 222mer fragment of *Streptococcus* that is linked to a 16-mer EO binding site, and has an actual length of 238 bases. Production of the single-stranded template follows this workflow (*6*): 1) Amplification of a 437 bp gene fragment from *Streptococcus* DNA; 2) Amplification of the 238 bp template, including EO hybridization sequence; 3) Removal of the biotinylated strand by binding the PCR product to streptavidin beads, and then heating to denature the strands; 4) cleanup and quantification of single-stranded DNA template.

*Oligonucleotides*

Primers for 437 bp *Streptococcus pneumoniae* 16s rRNA fragment:

Antisense: 5′-(BIOTIN)(BIOTIN)(PEG3)(PEG3)(PEG3)TGACAAGTGAAAGTGGCTGT-3'

Sense: 5′-PO4-CTAGCTAATACAACGCAGGTCCATCT-3′

Nested primers for 238 bp SBX template:

Sense: 5'-A(phosphorothioate)GAGTTTGATCCTGGCTCAG-3′

Antisense: 5'-(BIOTIN)(BIOTIN)(PEG3)(PEG3)(PEG3)TCATAAGACGAACGGAGGTAGTGA
TGCAAGTGCACCTTTTA-3′

437 bp fragment of *Streptococcus pneumoniae* (ATCC-700669) 16s rRNA:

5′–TGACAAGTGAAAGTGGCTGTGATATACTAATATAGTTGTCGCTTGAGAGAAGCAAGTGACAAAGACCTTT
GAAAACTGAACAAGACGAACCAATGTGCAGGGCGCTACAACATAAGTTGTAGTACTGAACAATGAAAAAACA
ATAAATCTGTCAGTGACAGAAATGAGTAAGAACTCAAACTTTTAATGAGAGTTTGATCCTGGCTCAGGACGA
ACGCTGGCGGCGTGCCTAATACATGCAAGTAGAACGCTGAAGGAGGAGCTTGCTTCTCTGGATGAGTTGCGA
ACGGGTGAGTAACGCGTAGGTAACCTGCCTGGTAGCGGGGGATAACTATTGGAAACGATAGCTAATACCGCA
TAAGAGTAGATGTTGCATGACATTTGCTTAAAAGGTGCACTTGCATCACTACCAGATGGACCTGCGTTGTAT
TAGCTAG–3′

100 bp HIV2 template:

5′–(PEG–3)GGGAAGATCTGGCCTTCCTACAAGGGAAGGCCAGGGAATTTTCTTCAGAGCAGACCAGAGCCA
ACAGCCCCACCAGAAGAGAGCTTCAGGTCTGGGGTAGTCCGTTCGTCTTATGA(PEG–3)–3′

60 bp 4-base repeat template:

5′–(PEG–3)ACTGACTGACTGACTGACTGACTGACTGACTGACTGACTGACTGACTGACTGACTGACTGTCC
GTTCGTCTTATGA(PEG–3)–3′

238 base Xp template:

5′–A(phosphorothioate)GAGTTTGATCCTGGCTCAGGACGAACGCTGGCGGCGTGCCTAATACATGCA
AGTAGAACGCTGAAGGAGGAGCTTGCTTCTCTGGATGAGTTGCGAACGGGTGAGTAACGCGTAGGTAACCTG
CCTGGTAGCGGGGGATAACTATTGGAAACGATAGCTAATACCGCATAAGAGTAGATGTTGCATGACATTTGC
TTAAAAGGTGCACTTGCATCACTACCTCCGTTCGTCTTATGA–3′

*Amplification of Strep 437mer*

50 µL reactions included 0.2 µM dNTPs (Promega), 10 ng genomic *Streptococcus pneumoniae* (ATCC-700669) DNA, 1 µM MolTaq 16S polymerase with 1x MolTaq buffer (MolZyme). Cycling conditions were 95 C for 30 seconds; 37 cycles at 95 C for 10 seconds, 60 C for 20 seconds, 72 C for 30 seconds; and final extension at 72 C for 3 minutes. PCR reactions were purified by the QIAquick PCR Purification Kit (Qiagen), eluting in water (Corning). Purified PCR product was quantified by A260 absorbance on a Nanodrop (ND2000).

*Amplification of Xp template (Strep 222mer)*

100 µL PCR reactions included 1 µM primers, 350 µM dNTPs, 1.2 pg of prepared Strep 437mer template, KOD polymerase (Millipore Sigma) and 1x PCR reaction buffer (Millipore Sigma). Cycling conditions were 95 C for 2 minutes; 33 cycles at 95 C for 10 seconds, 67 C for 20 seconds, 72 C for 30 seconds; and final extension at 72 C for 3 minutes. The 238 bp PCR product was then purified via QIAquick PCR Purification Kit (Qiagen), eluting in water. Purified PCR product was quantified by A260 absorbance measurement on a Nanodrop.

*Preparation of single-stranded Xp template (Strep 222mer)*

The purified 238 bp PCR product was combined with magnetic Dynabeads MyOne T1 streptavidin beads (ThermoFisher 65601) that had been pre-washed in 2x binding buffer (2 M NaCl, 20 mM Tris HCl, pH 8) at a ratio of 0.72 µL stock beads:1 pmol dsDNA, to achieve a final 1x concentration of binding buffer. After a 10-minute incubation at RT, the beads were washed for 1 minute in 500 µL 1x binding buffer including 280 pmols of mPEG biotin (Nanocs) per µL stock beads used to bind to any remaining free streptavidin. The beads were pelleted and resuspended in 30 µL of 20 mM Tris HCl, pH 8. The DNA was eluted by heating to 95 C for 1 minute, followed by pelleting of the beads and removal of the supernatant to a fresh 1.5 mL lo-bind DNA tube (Eppendorf). A 1 mL reverse phase column (Glen Research 60-5100) was prepared by pretreatment with 1 mL each of 100% acetonitrile (ACN), and 2M triethylammonium acetate (TEAA). The single-stranded product was brought to 500 µL in 100 mM TEAA and bound to the pretreated column. One mL each of 100 mM TEAA, 5 mM TEAA, and molecular biology grade water, in series, was used to wash the column. The product was eluted in 300 µL of 50% ACN. The sample was concentrated by speed vacuum centrifuge to about 130 µL to ensure ACN was evaporated. Final concentration was measured by nanodrop using an extinction coefficient of 2317200 L/(mole·cm), and template quality was assessed by 6% TBEU gel (Invitrogen) post-stained with SYBR Gold (ThermoFisher).

**Xp Synthesis Process**

*Microfluidic Card*

Xp Synthesis was performed in a cyclic olefin copolymer (COC) microfluidic chip, model Fluidic 243 from ChipShop [Fig. 2A]. The flow channel was nominally D-shaped, and was 0.6 mm wide x 0.30 mm deep. It was selected for high surface area to volume, high transparency below 365 nm and through the visible range, chemical resistance, and the ability to handle variable viscosities. Prior to elution, we used a single-direction flow from inlet to waste outlet for all reagents to best control the reaction conditions and washes. A short, dedicated fluidic path was used to limit passive adsorption of 'sticky' reaction components like Xp, template, and polymerase, while permitting larger volume washes. Copper tape (Digi-Key) was applied to the chip to assist heat spreading prior to mounting on a hot plate. Temperature was regulated between 23 C and 90 C as the process required.

*Surface Functionalization of Flow Channel*

The surface area along the first 50 µL of flow channel volume from the inlet was functionalized for Xp synthesis. Propargyl-PEG4-maleimide (Broadpharm) was diluted to 2 mM in 80% DMSO, 1% SDS for surface preparation. COC microfluidic chips were first rinsed with 80% DMSO, 1% SDS, then charged with propargyl-PEG4-maleimide solution and irradiated (365 nm, 4.2 mW/cm$^2$) for 20 minutes. These functionalized cards were washed with a solution of 50 mM sodium phosphate, 0.01% SDS, 10% DMSO.

*EO click to Flow Channel*

20 pmol of EO was combined with 0.6 mM THPTA, 6 mM Sodium ascorbate, 0.2 mM $CuSO_4$, 5 mM Aminoguanidine, 0.5 mM $MgCl_2$, and 10% DMF in a 50 µL click reaction, and was reacted with the functionalized flow channel for 20 minutes at 23 C. We estimate approximately 50% efficiency from this method (data not shown). The flow channel was flushed with Hybridization (H) buffer, (50 mM Tris-HCl (JT Baker), 500 mM $NH_4OAc$ (Millipore Sigma), 1 M Urea (JT Baker), 2% polyethylene glycol (PEG 8000) (Promega), pH 8.0) to remove copper and unclicked EO in preparation for template hybridization (*7*).

*Template hybridization*

4 pmol of Xp template (Strep 222mer) was added to 50uL of H buffer. Template + H was denatured at 90 C for 15 seconds, and then added to the pre-functionalized chip to hybridize template to EO. Hybridization was performed at a starting temperature of 72 C and ramped down to 37 C at a rate of 0.1 C/second.

*Xp synthesis reaction*

Holding the chip temperature at 37 C, 60 µL of SBX Extension Pre-wash (W) (50 mM Tris-HCl, 200 mM $NH_4OAc$, 20% PEG 8000, 10% N-Methyl-2-pyrrolidine (NMP) (Millipore Sigma), 0.05 µg/µL *Thermococcus kodakarensis* single-strand DNA binding protein (KOD SSB), 50 mM tetramethylammonium chloride (TMACl) (Tokyo Chemical Industry), 50 mM guanidinium chloride (GuHCl) (Millipore Sigma), 0.1 M Urea, 15 mM AZ-43,43 PEM, pH 8.0) was flowed through the channel. Next, 75 µL of SBX Extension Master Mix (A) (50 mM Tris-HCl, 200 mM $NH_4OAc$, 20% PEG 8000, 10% NMP, 0.05 ug/µL KOD SSB, 50 mM TMACl, 50 mM GuHCl, 0.1 M Urea, 15 mM AZ-43,43 PEM, 100 µM XNTPs, 1.1 mM $MnCl_2$, 0.26 µg/µL Xp Synthase, pH 8.0) was added to the chip and incubated at 37 C for 1 hour.

*Post-extension processing*

After primer extension, PTFE tubing was connected to the inlet adapter and the waste line was opened. 1000 µL of B Buffer (B) (30% Acetonitrile (ACN), 100 mM HEPES (JT Baker), 100 mM $NaHPO_4$, 1% Tween-20, 3% Sodium Dodecyl Sulfate (SDS) in $D_2O$ (Millipore Sigma), pH 8) was added to the chip to quench the extension reaction and wash residual extension reaction components off of the chip. The chip temperature was adjusted to 23 C, and 200 µL of Cleavage Buffer (C) (7.0 M HCl (Millipore Sigma)) was added to the chip to quantitatively cleave Xpandomer P–N backbone bonds for 40 minutes. The cleavage reaction was neutralized by flushing the chip twice with 1000 µL of B buffer. B Buffer was mixed with Modification Buffer (Mod) (2.5 M SA in 100% ACN) to a final SA concentration of 667 mM, and 300 µL of the mixture was incubated for 5 minutes on the chip to neutralize the positive charge on the cleaved Xpandomer backbone. The B + Mod buffer was flushed twice with 1000 µL of Post-mod buffer (35% ACN, 15% isopropanol (Millipore Sigma)).

*Photocleavage and selective Xp elution*

The chip temperature was adjusted to 37 C, and 100 µL of buffer D1 (50% ACN in water) was added. The chip was placed 2.5 cm below a UV light source (Firefly 25x10) for 15 seconds to photocleave the Xpandomer from support, and then returned to 37 C on the breadboard. After incubation for 2 minutes, 100 µL was drawn from the chip inlet side as elution 1 (E1), and removed to a Protein LoBind collection tube (Eppendorf). The remaining Xp was stripped from the chip by adding 1000 µL of Buffer D2 (100 mM GuHCl, 20 mM sodium hexanoate (Millipore Sigma), 100 mM HEPES, 35% ACN, 15% DMSO, pH 7.4) and incubating for 2 minutes before drawing off 100 µL from the inlet side as E2.

*In-Solution Xp Synthesis*

In-solution Xp syntheses were performed on 45-mer, 100-mer, or 222-mer *Streptomycin* templates. 1 pmol of template mixed with 1 pmol of 5′-SIMA-HEX-labeled EO was added to a 10 µL reaction of SBX Master Mix (50 mM Tris-HCl, 200 mM $NH_4OAc$, 20% PEG8K, 10% NMP, 0.05 µg/µL KOD SSB, 50 mM TMACl, 0.1 M Urea, 15 mM PEM, 100 µM XNTP, 1.1 mM $MnCl_2$, 0.26 µg/µL Polymerase, pH 8) in a PCR tube and incubated for 60 minutes at 37 C in a thermocycler. Reaction was quenched by adding 10 µL of quench solution (0.4% SDS, 1.15x TBE-Bromophenol Blue Gel Loading Buffer ((12% w/v Ficoll, 1X TBE Buffer (Corning), 0.01% w/v Bromophenol Blue (JT Baker)), 0.02 U/µL Proteinase K) and incubating at 70 C for 2 minutes. Half of the reaction was run on a 2.5% TBE agarose gel (Lonza) for visualization of extension products [Fig. S20A-I].

**Sequencing Data Collection & Analysis Methods**

*Sequencing Platform & Instrumentation*

The methods and the sequencing system used herein were adapted from Marziali (*8*). Fig. S22 shows a schematic overview of the instrumentation. Fig. 4 shows a schematic of the sequencing cell. The sequencing chemistry and biological components were assembled on the surface of a PTFE/FEP support and positioned in the PTFE cell. The cell had *cis* and *trans* chambers that were fluidically connected through a U-shaped tube. AgCl electrodes were placed in each chamber and were connected to the CV 203-vBU headstage attached to an Axopatch 200B amplifier. One end of the tube is narrowed to a 13-micron diameter aperture across which the lipid bilayer was formed. The Axopatch 200B applied a potential across the nanopore via the headstage and amplified the ion current measurements with a 1 mV/pA transimpedance gain using a lowpass 4-pole Bessel filter at 10 kHz. The National Instruments USB-6351 digitized this output at 16 bits and 100 kS/s simultaneously with the applied voltage measurement. In addition, it generated the analog waveform for control of the applied potential. A LabVIEW program stored the digital data and controlled the USB-6351 operation [Fig. S22]. The PTFE cell was coupled to a custom thermal controller to maintain the operating temperature at 20 C. External electromagnetic interference was dampened by a faradic cage that covered the cell while external mechanical noise was reduced by positioning the cell on an inflated rubber dampener.

*Buffers*

Lipid Mix: 18.49 mg/mL 1,2-di-O-phytanoyl-sn-glycero-3-phosphoethanolamine (DPhPE) (Corden), 0.12% hexadecene (Tokyo Chemical Industry), 99.88% hexane (Alfa Aesar), prepared in an argon environment (0.6-1.1 ppm $H_2O$) and stored at -35 C

Pore Insertion/*Trans* Buffer: 2 M $NH_4Cl$ (Millipore Sigma), 100 mM HEPES (JT Baker) at pH 7.4

Nanopore Mix: 0.06 µg/µL of αHL-nanodisc complex stored in Nanodisc Elution Buffer

*Cis* Buffer: 0.4 M $NH_4Cl$, 0.6 M GuHCl (Macron Fine Chemicals), 100 mM HEPES (JT Baker) at pH 7.4

Xp Dilution Buffer: 0.8 M $NH_4Cl$, 1.2 M GuCl, 200 mM HEPES; pH 7.4

Xp Sample Load: 50:50 ratio of Xp elution stored in 50% ACN (Glen Research) and Xp Dilution Buffer. Vortex at 3000 rpm for 5 s, followed by 2 s spin down. (resulting product): Xp in 0.4 M $NH_4Cl$, 0.6 M GuHCl, 100 mM HEPES; pH 7.4, 25% MeCN

Lysis Solution: 50 mM $NH_4OAc$ (Millipore Sigma), 1 mM EDTA (Millipore Sigma), 20 mM Tris-OAc (Thermo Scientific), 0.0125% ReadyLyse (LGC Biosearch), 0.5 mM PMSF (Millipore Sigma), pH 8.3

EQ10 solution: 20 mM tris base (JT-Baker), 10 mM imidazole (Millipore Sigma), pH 7.5

αHL Wash Buffer: 25 mM tris-OAc (Millipore Sigma), 25 mM imidazole (Millipore Sigma), 150 mM NaCl (Millipore Sigma), pH 7.5

αHL Elution Buffer: 25 mM tris-OAc (Millipore Sigma), 250 mM imidazole (Millipore Sigma), 150 mM NaCl (Millipore Sigma), pH 7.5

0.2x αHL Storage Buffer: 25 mM NaCl (Millipore Sigma), 10% glycerol (Teknova), and 10 mM Tris (JT-Baker), pH 8.5

Nanodisc Running Buffer: 500 mM NaCl (Millipore Sigma), 20 mM Tris-HCl (JT-Baker), pH 7.4

Nanodisc Wash Buffer: 500 mM NaCl (Millipore Sigma), 10 mM imidazole (Millipore Sigma), 20 mM Tris-HCl (JT-Baker), pH 7.4

Nanodisc Elution Buffer: 250 mM imidazole (Millipore Sigma), 20 mM Tris-HCl (JT-Baker), pH 7.5

Sequencing Buffer: 0.4 M $NH_4Cl$ (Millipore Sigma), 0.6 M GuHCl (Macron Fine Chemicals), 100 mM HEPES (JT Baker) at pH 7.4, 5% glycerol (AMRESCO)

*Pore-lipid Assembly*

Reagent mixtures referenced here are defined in the Buffers section. To insert a single nanopore into a lipid bilayer, <1 µL of the Lipid Mix was deposited and dried on the FEP support. Immediately after drying, argon gas was forced from the *trans* side of the tube to clear any excess Lipid Mix that had seeped past the aperture. The cell was then placed under vacuum for 1 minute and positioned in the faradic cage. The tube, *cis* chamber, and *trans* chamber were then filled with Pore Insertion/*Trans* Buffer.

Lipid bilayers were formed across the aperture by pushing and pulling air bubbles with a micropipette while a low voltage was applied to monitor current. Bilayer stability was assessed through a destructive test using linear voltage sweeps from 0 V to 1 V at a rate of 10 V/s to determine bilayer resilience. A discrete breakage point at >600 mV indicated stable bilayers were forming with the bubbling technique. If needed, additional vigorous bubbling across the aperture was used to increase bilayer strength.

To assemble a single alpha hemolysin (αHL) pore in a bilayer, air bubbles were pushed and pulled across the aperture with a micropipette while a voltage is applied to drive, orient, and monitor pore insertion. 1.0 µL of the Nanopore Mix was added to the *cis* well above the support surface and triturated. While using the bubbling technique, voltage was regularly adjusted so that bilayers were forming and breaking at an approximate 50:50 ratio. The magnitude of the working voltage typically varied within 500 mV and 800 mV using either a positive or negative potential. Voltage was turned off once a pore was detected so that pore characterization metrics could be collected. A fully formed single pore oriented with the barrel facing the *cis* chamber was determined by assessing the conductance asymmetry and amplitude (mean current at +100 mV was 160 pA to 180 pA with <1.45 pA $I_{RMS}$, mean current differential between +100 mV and -100 mV was -28 pA to -35 pA). After qualifying a properly oriented single nanopore, the solution in the *cis* chamber was replaced with 3 mL of *Cis* Buffer while siphoning away the existing solution.

*Xp sample loading*

6 μL of prepared Xp Sample Load was dispensed and mixed thoroughly by triturating 40 times around the *cis* chamber, then incubated for 2 minutes. This step was performed 4 times to deliver a total of 24 μL. The *cis* chamber was replaced with 3 mL of Sequencing Buffer while siphoning away the existing solution. The measurement voltage waveform was applied for ion current recording.

*Alpha-Hemolysin Synthesis*

Reagent mixtures referenced here are defined in the Buffers section. αHL pore was expressed and stored in *E. coli* using the same method as for Dpo4 and SSB, elaborated above. For purification, frozen cell pellets were resuspended on ice in 8 mL Lysis Solution. Lysis proceeded on ice for 30 minutes, followed by mechanical shearing of DNA achieved by 4 passes through a 21-gauge needle. Streptomycin sulfate (1% final) was added dropwise from a 30% solution while vortexing the lysates at 1000 rpm, until chromatin had visibly condensed. Crude lysates were cleared by centrifugation at 20,000 x g for 15 minutes at 4 C. Cleared lysates were prepared for binding to Ni-NTA resin (Thermo Scientific) by adding 6 mL EQ10 Solution. Ni-NTA resin that had been washed twice in EQ10 was combined with the lysate, and incubated for 15 minutes, rotating at 15 RPM on a Rotator Genie. The protein-bound resin was rinsed twice in aHL Wash Buffer, before eluting with αHL Elution Buffer. The eluates were desalted with PD Midi columns into 0.2x αHL Storage Buffer. Tobacco Etch Virus (TEV) protease (New England Biolabs) reactions were incubated overnight according to manufacturer's recommendations at a ratio of 0.3 U TEV protease:1 μg αHL, to cleave off the N-terminal 6x His tag. The cleaved tags were removed by incubation with Ni-NTA resin that had been pre-equilibrated with 0.2x αHL Storage Buffer. The flowthrough was collected, and the resin was washed with two column volumes of αHL Wash buffer. The combined wash and flowthrough fractions were desalted as before, and then concentrated 5x in a SpeedVac under no heat. After quantification by 280 nm absorbance, 2 μg of each sample was evaluated by protein gel to verify cleavage.

*aHL-nanodisc Complex Synthesis*

Reagent mixtures referenced here are defined in the Buffers section. αHL monomers were isolated by size exclusion chromatography (SEC) using a Superose 6 Increase 10/300 GL column (Cytiva) on an Prostar HPLC system (Varian). aHL monomer was combined with reconstituted MSP1D1 (Sigma-Aldrich) and 1,2-diphytanoyl-sn-glycero-3-phosphocholine (DPhPC) (Avanti) at a ratio of 1:0.86:17.11 in a buffer containing 19.7 mM Sodium Cholate (Millipore Sigma). After a 1 h incubation at RT, 0.5 mg/μL Biobeads (Bio-Rad SM-2) were added to the complex-forming mixture. The Biobead mixtures were shaken at 1200 rpm for 2.5 h to remove free lipid, and the samples were eluted from the beads. The samples were passed through a 0.45 μm syringe filter before loading onto a Superose 6 Increase 10/300 GL column to isolate the monodisperse complexes. The purified nanodisc-aHL complexes were further purified by Ni-NTA batch purification, using resin pre-equilibrated with Nanodisc Running Buffer. The resin was washed using Nanodisc Wash Buffer, and eluted in Nanodisc Elution Buffer. This reagent product is referred to here as the aHL-nanodisc complex elution and stored at 4 C.

*Sequencing Run & LabVIEW operation*

The waveform applied to the *cis* reservoir relative to the *trans* reservoir was -70 mV DC interspersed with a 6 µs pulse of -600 mV applied at 1 kHz rate. A LabVIEW program supported single pore formation, operational control, structured header information, timed data capture, and the eject cycle. All subsequent analysis was performed with MATLAB scripts.

The eject cycle was called dynamically during the run when the pore ion current exhibited a blockage. To remove the blockage, -250 mV was applied for 400 ms to eject the stalled Xp molecule from the pore before returning to the measurement waveform.

*Data Analysis*

The ion current and applied voltage data for the run were recorded in a raw data file for post-processing with 16 bits sampling at 100 kS/s. A MATLAB script processed the raw data file following a data path summarized in Fig. S23A.

Initial post-processing of the raw ion current file removed periodic pulses that appear on the current trace in response to the voltage pulses used to step the Xp translocation. Pulse removal replaced data samples with extrapolated data of the reporter following the pulse and was done to maintain data timing alignment and read visualization. No extrapolated data was used for the base calling. Fig. S23B-C shows an example of raw ion current data traces before and after the pulse removal. The latter graph shows an initial open channel ion current of 71 pA, followed by a drop in ion current to the four reporter blockage levels ranging between 6 to 22 pA when the Xp molecule enters the nanopore. The pulses coincide with distinct shifts in current level.

After pulse removal, the data was downsampled to 20 kS/s, parsed into the individual translocation events and normalized to the open channel current. Events were then prefiltered to remove events shorter than a pulse period, those with voltage abnormalities, or deep blockage events.

A histogram of samples from this refined data subset [Fig. S23D] were plotted and basecall demarcation ranges were determined using midpoints between the 4 primary current level peaks. Data was parsed into 1 ms periods, of 20 ion current samples, that start at a pulse trigger and end just prior to the following pulse trigger. The first 10 data points of each period were ignored, which included pulse-substituted samples and allowed for post-pulse settling time. The median of the remaining 10 data points was designated the basecall level. The basecall level was then converted to the corresponding basecall using the level demarcation range.

The sequence of basecalls for each translocation event was aligned with the known template DNA sequence using a Smith-Waterman routine (*9*) using gap score weighting of [1, -1, -1, -1] for base matches, deletions, insertions and substitutions, respectively. Outputs from this analysis provided aligned reads and detailed error counts. There were no read truncations from the Smith-Waterman alignment on 59% of the reads, though the mean truncation was 4.1 bases per read. Most of this truncation mean came from <1% of the reads that appeared to be in a temporary blocking state with false basecalls that caused >100 truncations per read in this limited set. The run error report used only reads exceeding 10 bases in length, and was defined by the ratio of the sum of the errors of each type (insertions, deletions and substitutions) over the sum of the aligned read lengths. Examples of ion current traces with indicated errors are presented in Fig. S24.

*Blocked State examples*

Blocked states have multiple phenotypes of which a few examples are shown in Fig. S25A-E. Most cases begin as a normal Xp translocation and stop at a noisy state, where normalized ion currents range from 0 to 0.5, deeper and higher than the 4 reporter levels (0.09, 0.16, 0.23 and 0.31). In some cases, blockages lead to oscillation between 2 states.

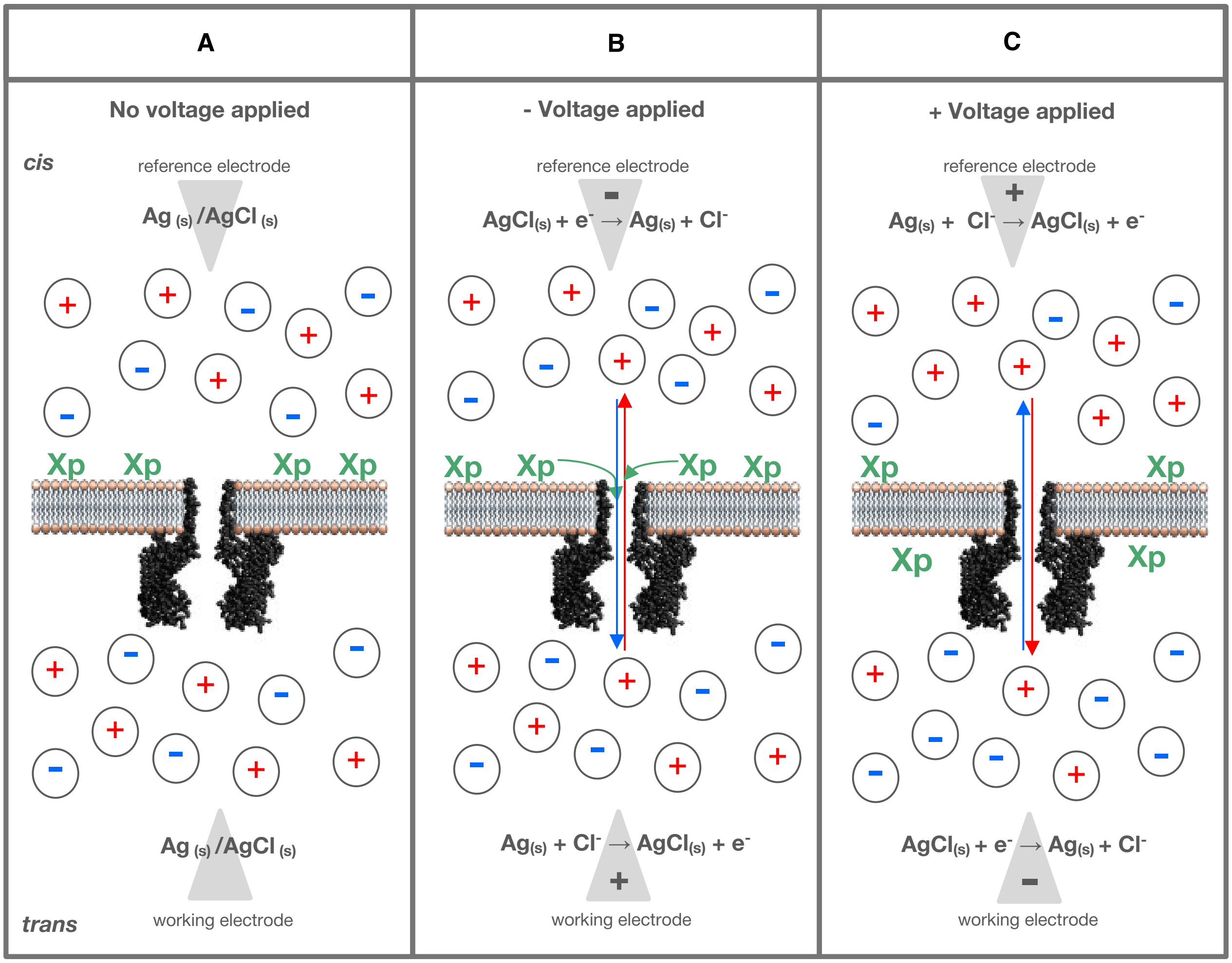


**Fig. S1. Ag/AgCl redox reaction controls ion migration in solution. (A)** The sequencing electrolyte contains both cations (red: $NH_4^+$, $Gu^+$) and anions (blue: $Cl^-$). Xp sample (green) is loaded into the lipid bilayer prior to sequencing waveform application. Reference and working solid AgCl electrodes are positioned on the *cis* and *trans* side of the bilayer, respectively. **(B)** To measure and translocate Xp sample through the pore, a positive voltage potential is set at the working electrode, with a counter negative voltage potential at the reference electrode. Under an applied voltage, an electric field drives redox reactions at both electrode-solution interfaces generating $Cl^-$ ions at the reference electrode and absorbing them at the working electrode, causing an ion current through the solution. Anions and Xp molecules migrate through the pore towards the *trans* electrode while cations migrate towards the *cis* electrode. **(C)** To clear blockages in the pore, the voltage is occasionally inverted, reversing the ion current and ejecting the blocking Xp back into the *cis* reservoir. Translocated Xp molecules are not captured again when the voltage is reversed due to the leader length relative to the vestibule size [Fig. S21A]. This allows for one directional sequencing and prevents re-sequencing Xp molecules.

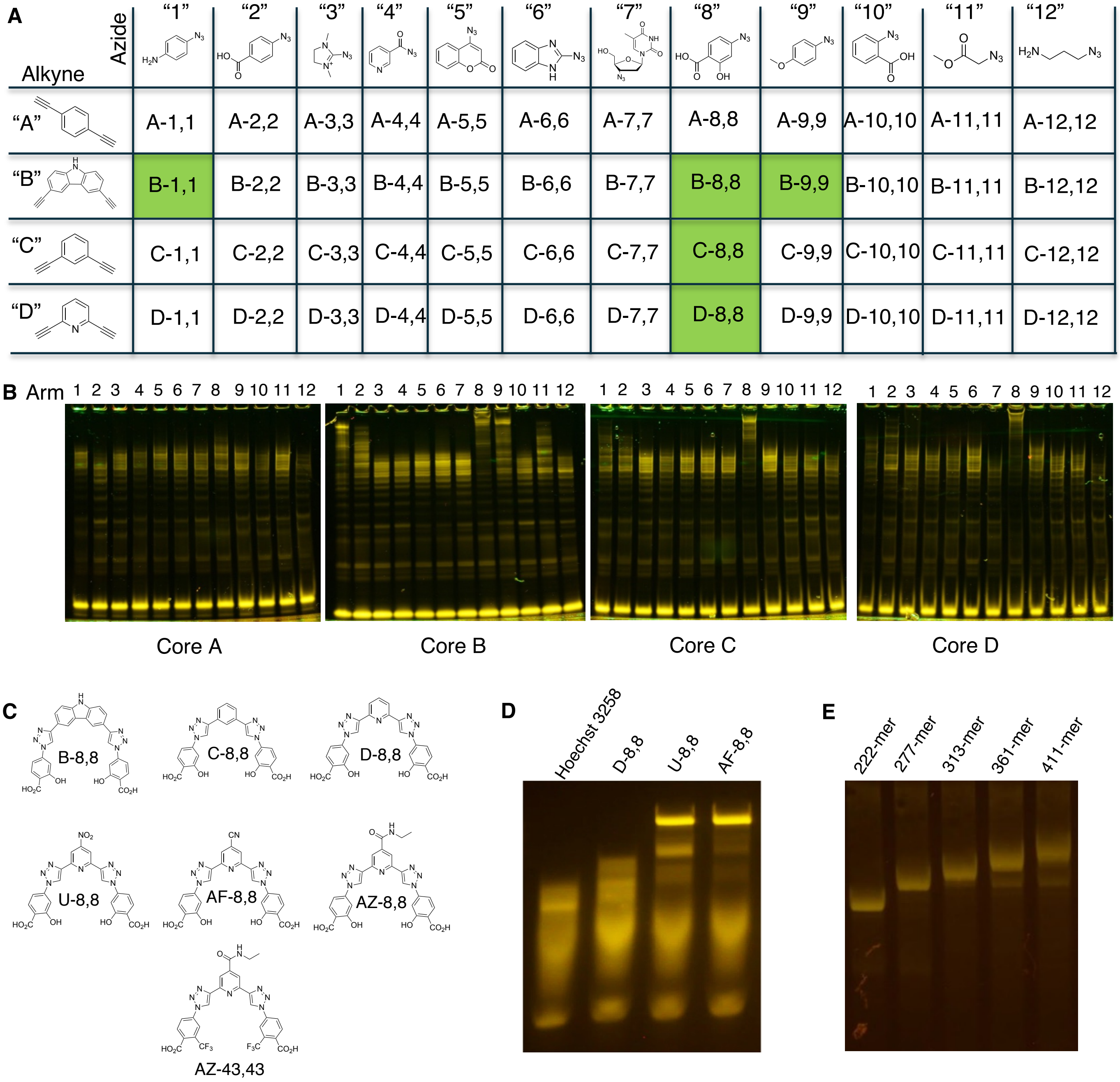


**Fig. S2. Polymerase Enhancing Moieties (PEMs). (A)** A matrix of bis-alkyne cores was clicked with azide arms to explore a diverse functional space. **(B)** The products of the click were used as additives in Xp synthesis using a fluorescent primer, a 45-mer template and analyzed (without acid expansion) by 4-12% TBE PAGE. Several hits resulted in significantly enhanced Xp length (i.e. "PEM effect", shaded green in the matrix). Of these hits, the D-8,8 structure had the largest enhancement. **(C)** Structures demonstrating the evolution of PEM cofactors. **(D)** Additional screening of targeted functional group changes on a 100-mer template, analyzed by electrophoresis in 4% TBE agarose. **(E)** 2.5% TBE agarose gel analysis of extension reactions that all feature AZ-43,43 PEM shows minimal 'shortmers' up to a template length of 313, and significant full-length extension product from a 411-mer template.

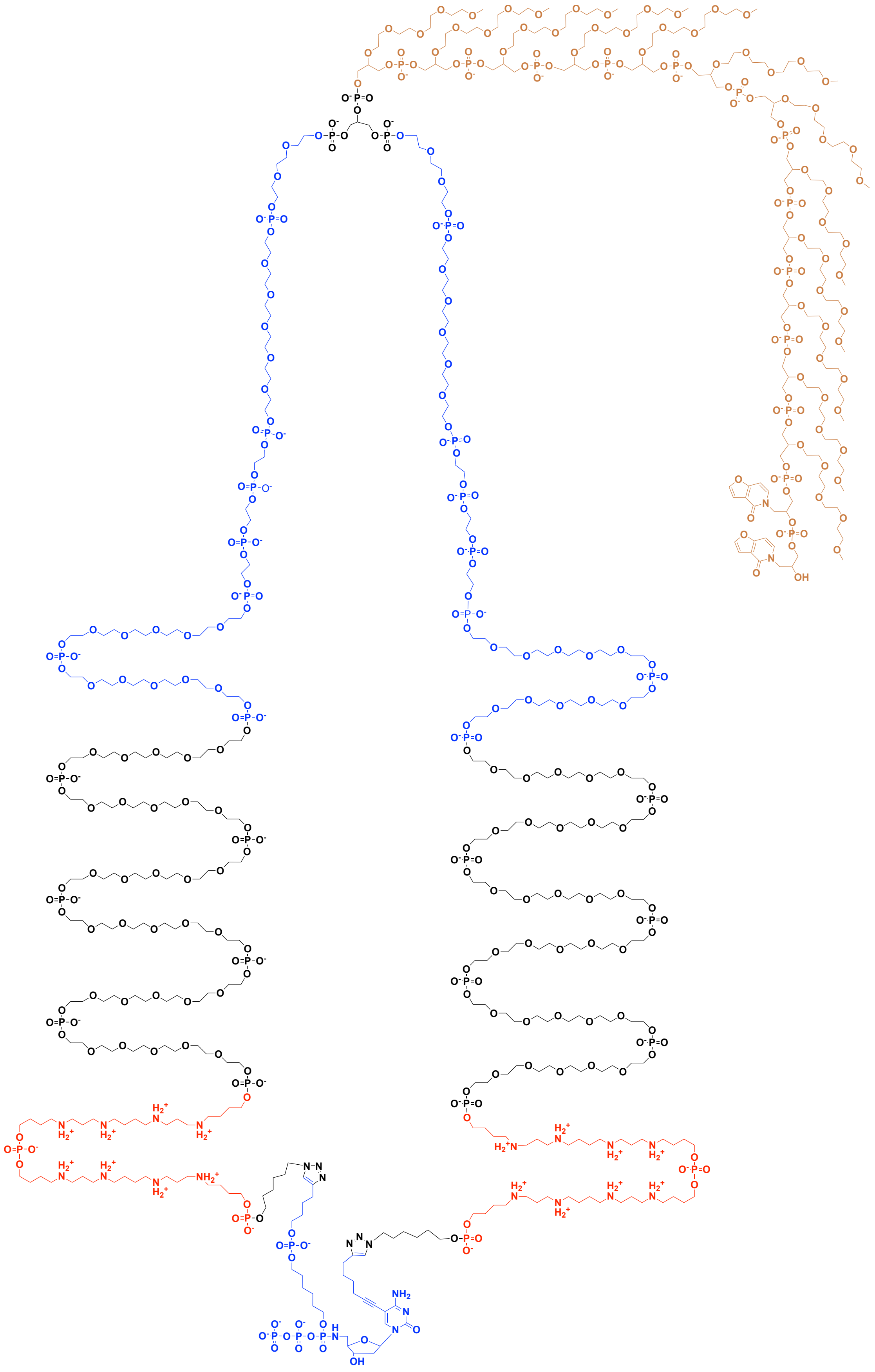

**Fig. S3. Structure of XCTP with level 1 tether.** The SSRT containing the enhancer (red), level 1 (blue) and TCE (gold) are clicked to the bis-alkyne dCTP.

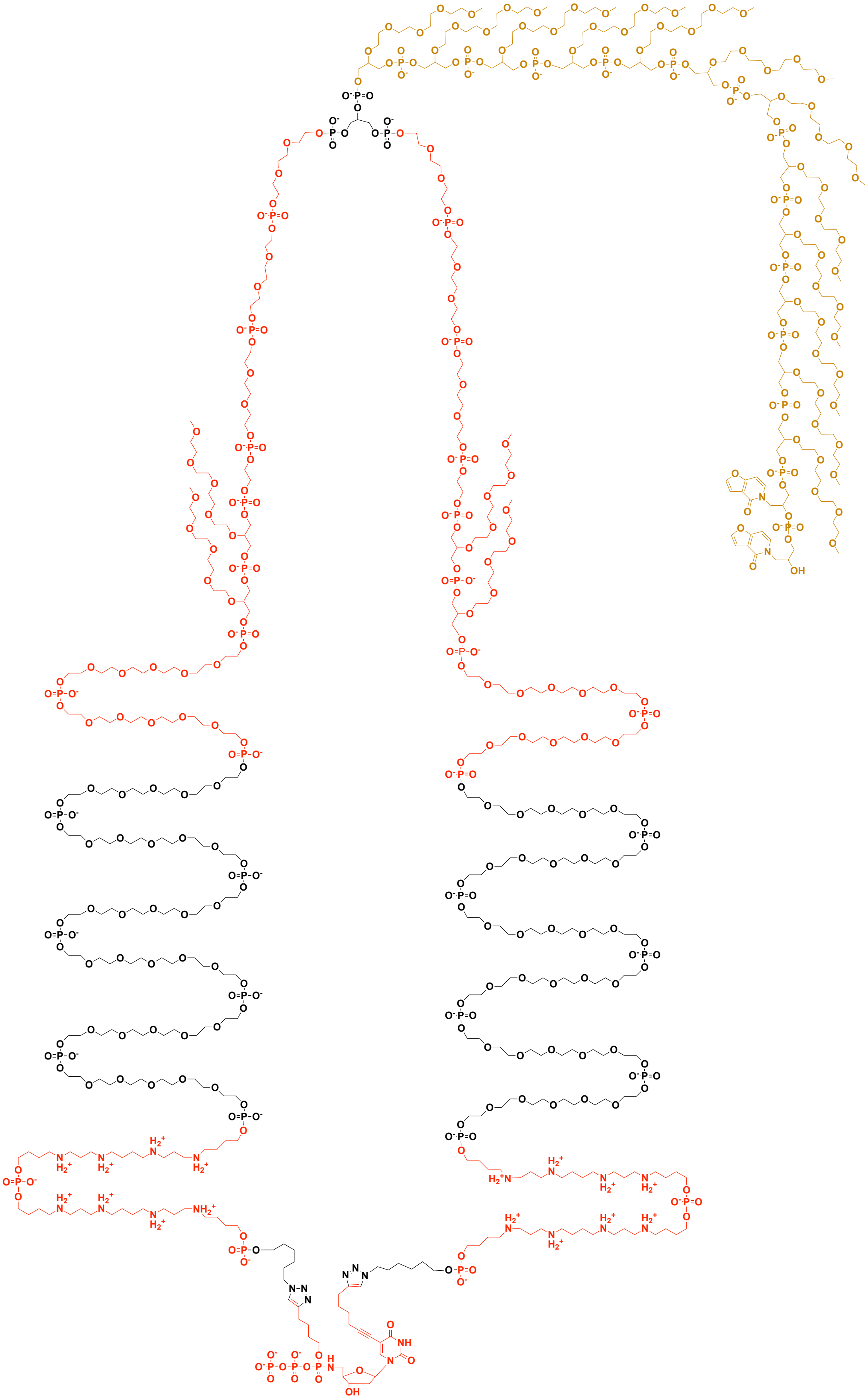

**Fig. S4. Structure of XTTP with level 2 tether.** The SSRT containing the enhancer (lower red), level 2 (upper red) and TCE (gold) are clicked to the bis-alkyne dTTP.

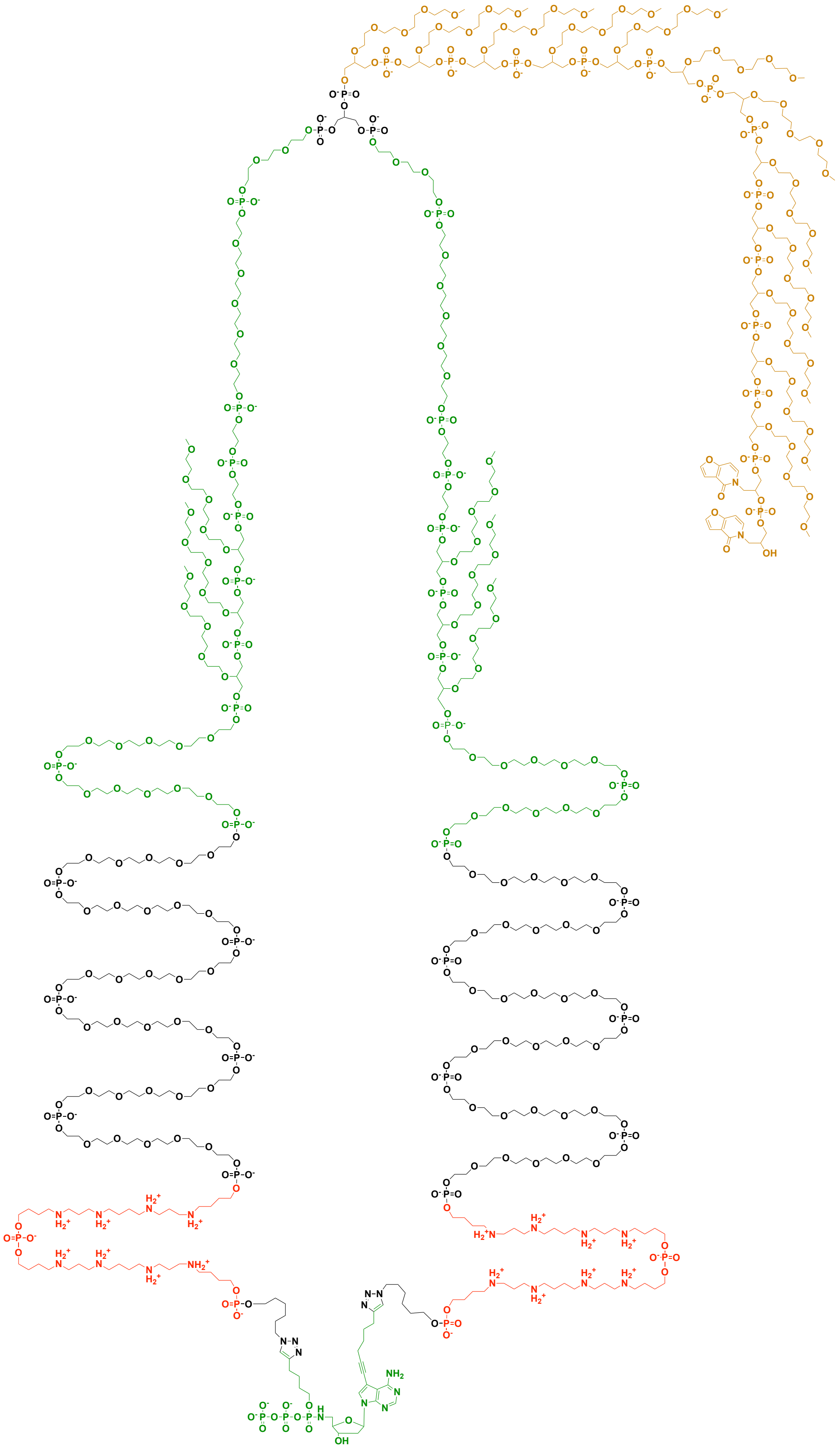

**Fig. S5. Structure of XATP with level 3 tether.** The SSRT containing the enhancer (red), level 3 (green) and TCE (gold) are clicked to the bis-alkyne dATP.

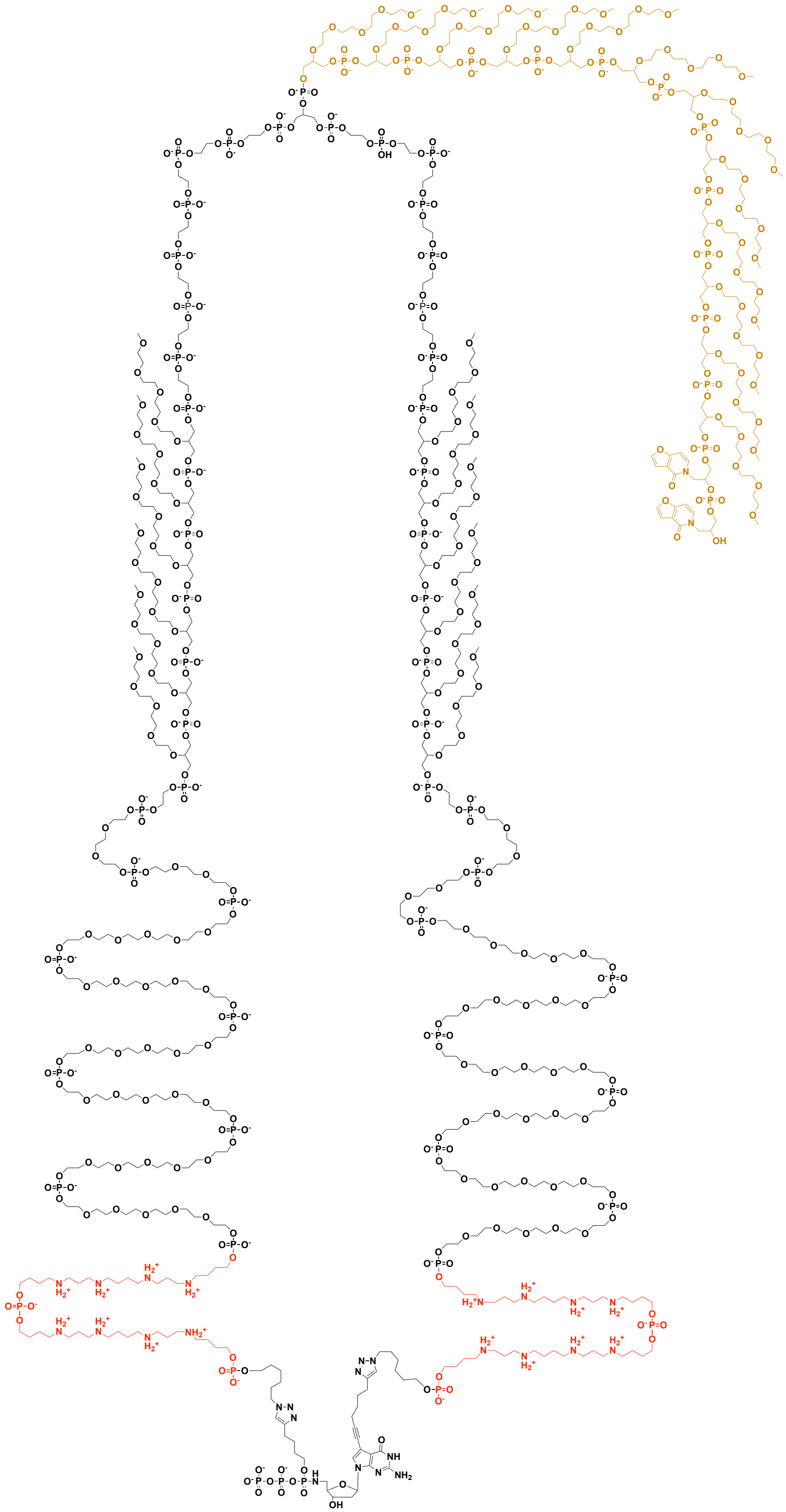

**Fig. S6. Structure of XGTP with level 4 tether.** The SSRT containing the enhancer (red), level 4 (black) and TCE (gold) are clicked to the bis-alkyne dGTP.

**Pendant mPEG-4**
**(4)**

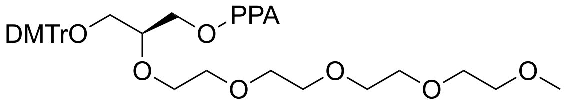

**PEG-6**
**(D)**

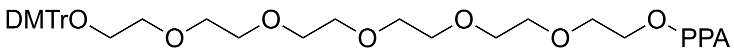

**PEG-3**
**(X)**

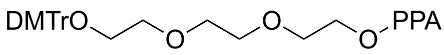

**C12**
**(Z)**

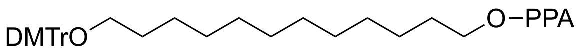

**Spermine**
**(Q)**

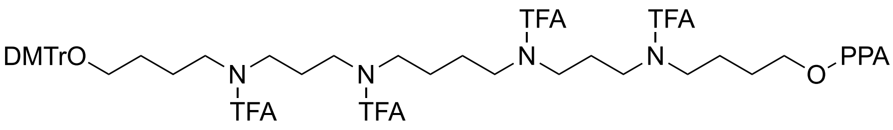

**Photo Cleavable Spacer**
**(Pc)**

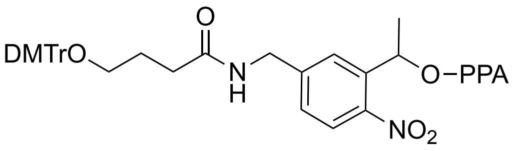

**Brancher**
**(Y)**

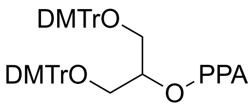

**C2**
**(L)**

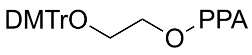

**Bromohex**
**(R)**

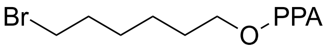

**Benzofuran**
**(5)**

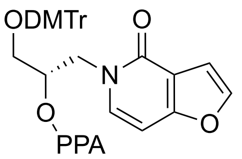

**Fig. S7. Glossary of Phosphoramidites (PPA).**

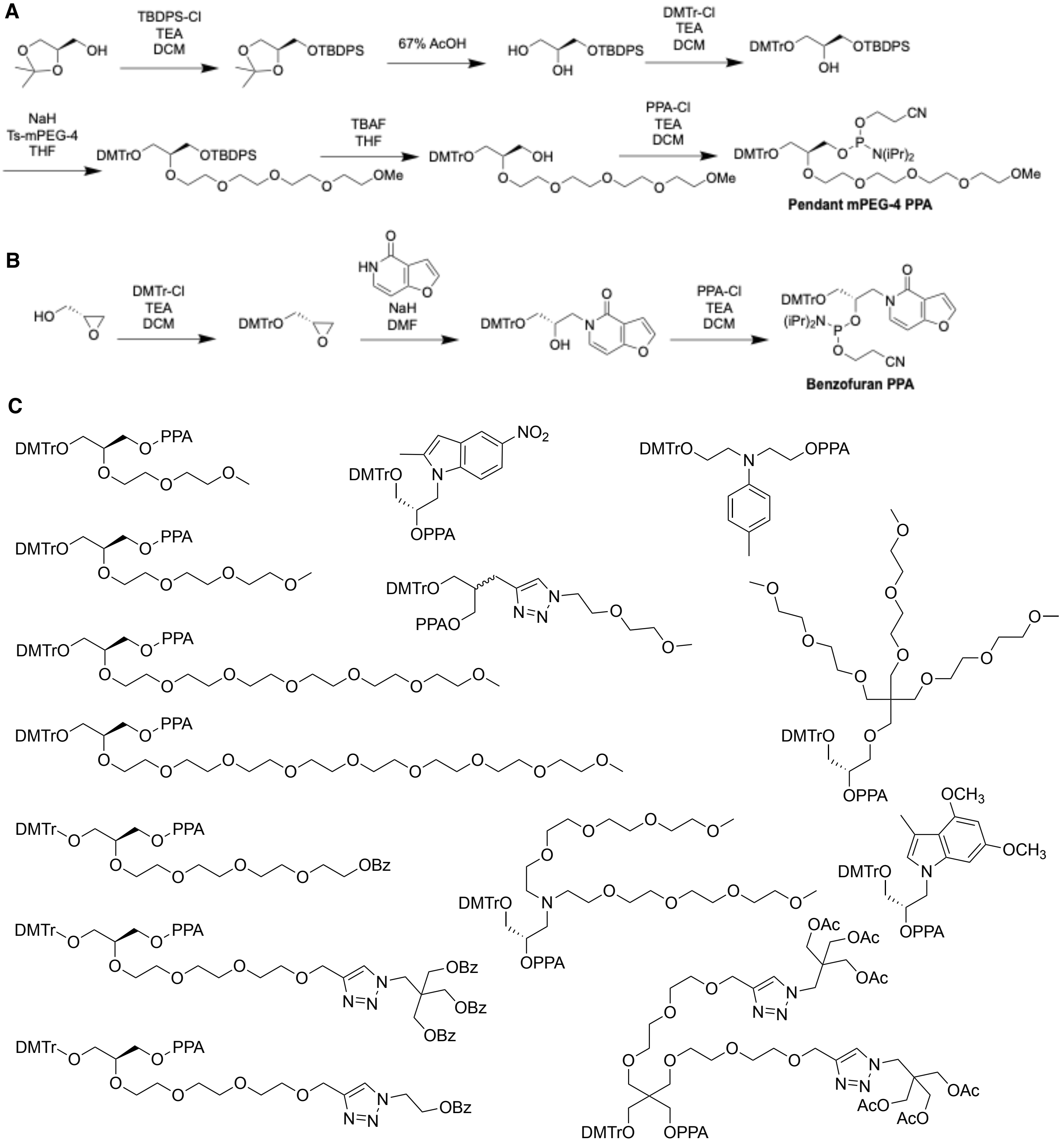


**Fig. S8. (A)** Synthesis of pendant mPEG-4 from the optically pure isopropylidene glycerol. The free alcohol was protected as the t-butyldiphenylsilyl ether, then the diol was deprotected with acid. The primary alcohol was selectively tritylated, then a tosyl-mPEG-4 was added. The silyl ether was removed and the alcohol was phosphitylated. **(B)** Synthesis of the benzofuran phosphoramidite began with optically pure epoxide, which was tritylated. The heterocycle was added to open the ring and the resulting alcohol was phosphitylated. **(C)** A selection of the phosphoramidites synthesized for testing in SSRT structures.

| Level | XNTP | Linker | Enhancer | Spacer | Reporter | Brancher | TCE |
|---|---|---|---|---|---|---|---|
| **1** | **C** | **R** | **QQ** | **DDDDDD** | **DDLLLDX** | **Y** | **44444444444455** |
| **2** | **T** | **R** | **QQ** | **DDDDDD** | **DD44LXXX** | **Y** | **44444444444455** |
| **3** | **A** | **R** | **QQ** | **DDDDDD** | **DD444LLDX** | **Y** | **44444444444455** |
| **4** | **G** | **R** | **QQ** | **DDDDDD** | **XXL444444LLLLLLL** | **Y** | **44444444444455** |

**Fig. S9. SSRT PPA Sequence.** SSRTs are synthesized with conventional oligonucleotide methods. Using universal support, synthesis starts with the benzofuran PPA (5). After 12 incorporations of the pendant mPEG-4 PPA (4), a branching PPA (Y) is incorporated. From the brancher, two symmetric arms are synthesized starting with the reporter. Each reporter has a unique sequence that varies charge density via phosphate spacing with linear PEG or alkyl chain and number of pendant PPAs, allowing for 4 distinct levels. This is followed by 6 PEG-6 PPA (D) spacers, 2 spermine PPA (Q) incorporations and finishing with Bromohex PPA (R) that is post synthesis modified to an azide.

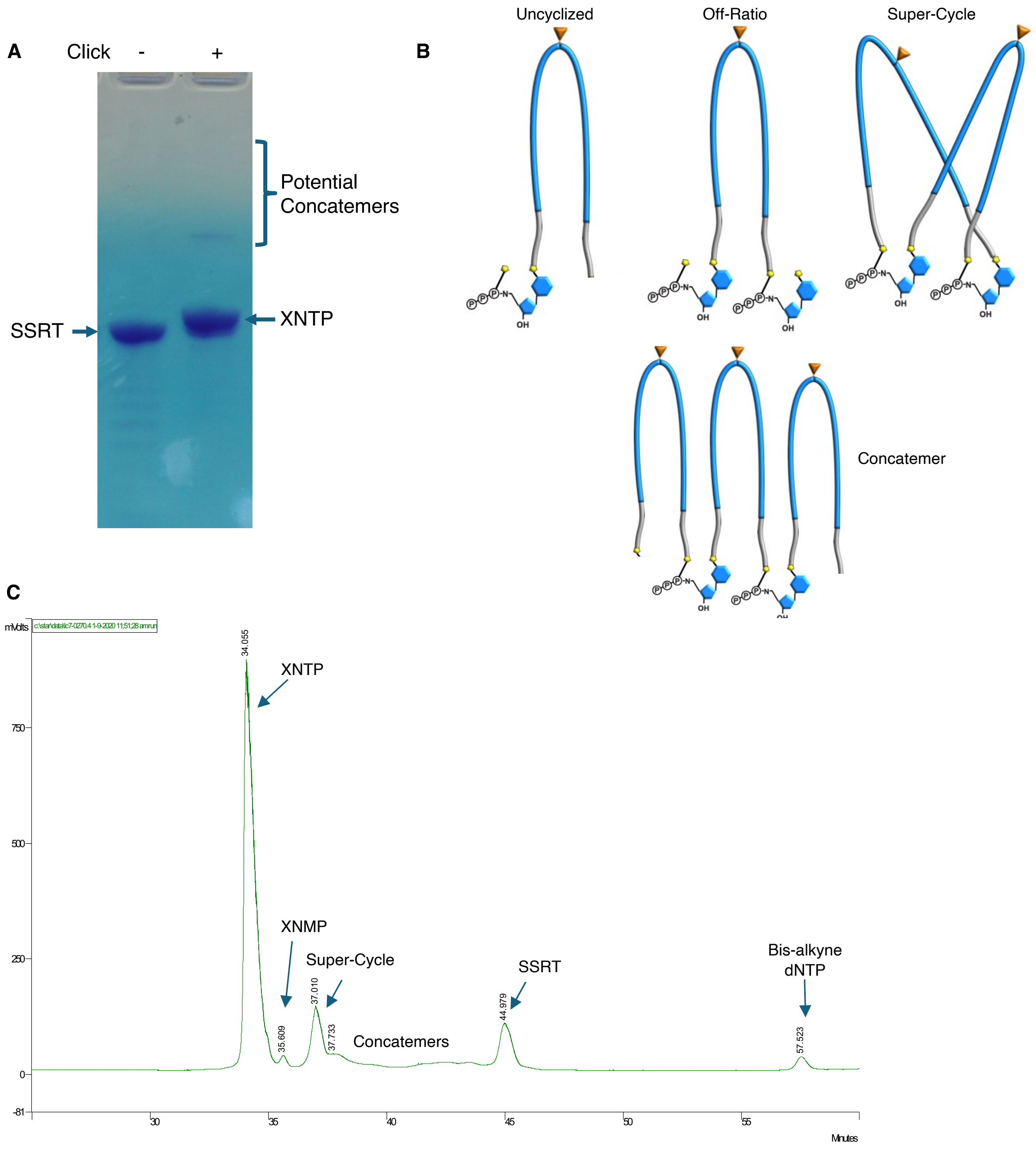


**Fig. S10. Click of XNTP. (A)** PAGE analysis (250 pmol load on 15% acrylamide gel, post-stained with toluidine blue) of a representative SSRT before and after click to form an XNTP. The cyclization slightly impacts the gel mobility, though potential concatemers migrate significantly slower. **(B)** Identity of several potential click side-products that would result in strand breaks. **(C)** Preparative chromatogram of XNTP click. The cyclized product was purified by RP (C18) HPLC in 100 mM TEAA water/acetonitrile (product elutes ~24% acetonitrile), 250 nm detection with > 50% purified yield.

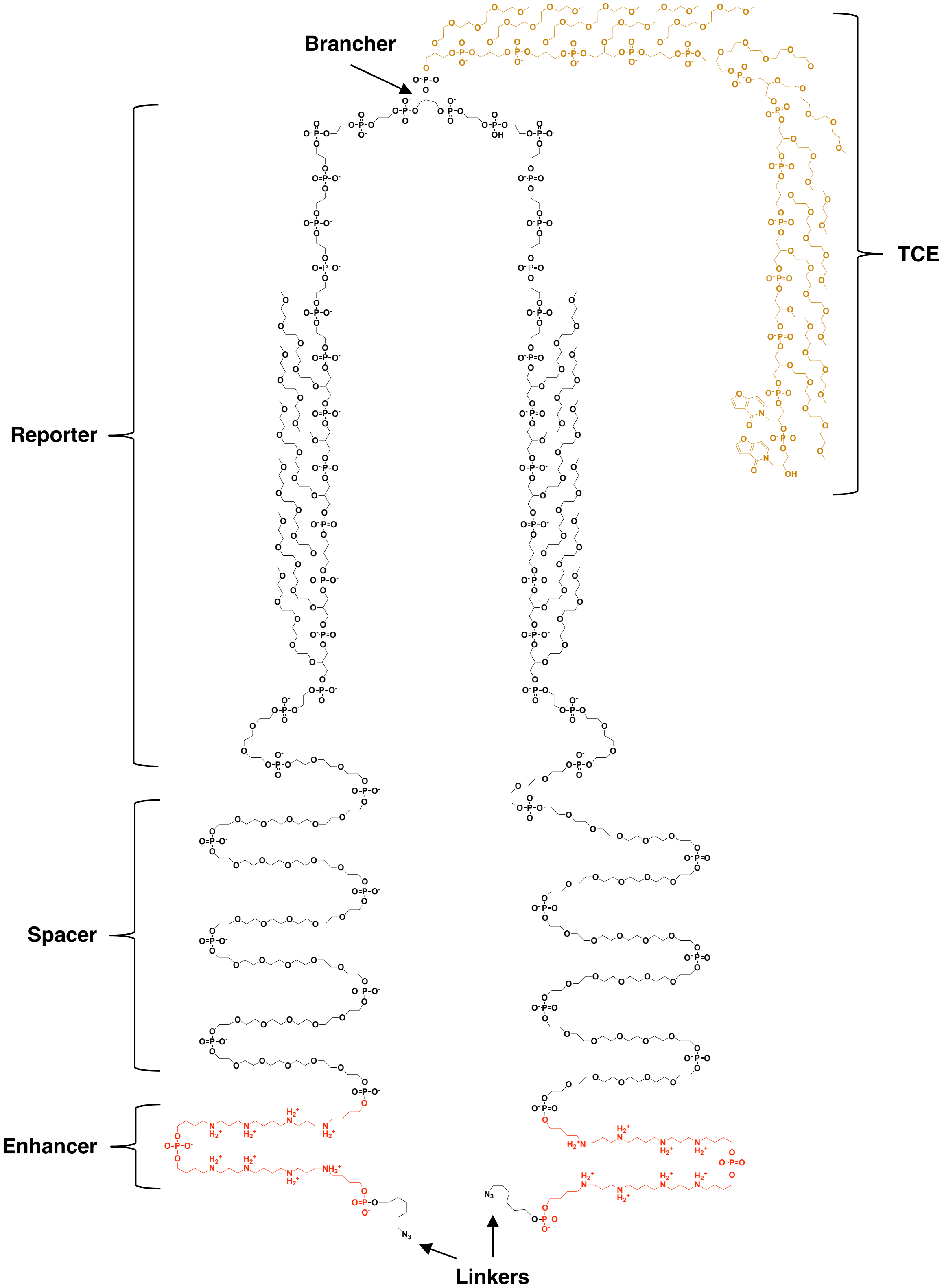


**Fig. S11. Level 4 SSRT.** Structure of the symmetrically synthesized reporter tether that is clicked to bis-alkyne dGTP to form the XGTP. Design and functional elements are annotated.

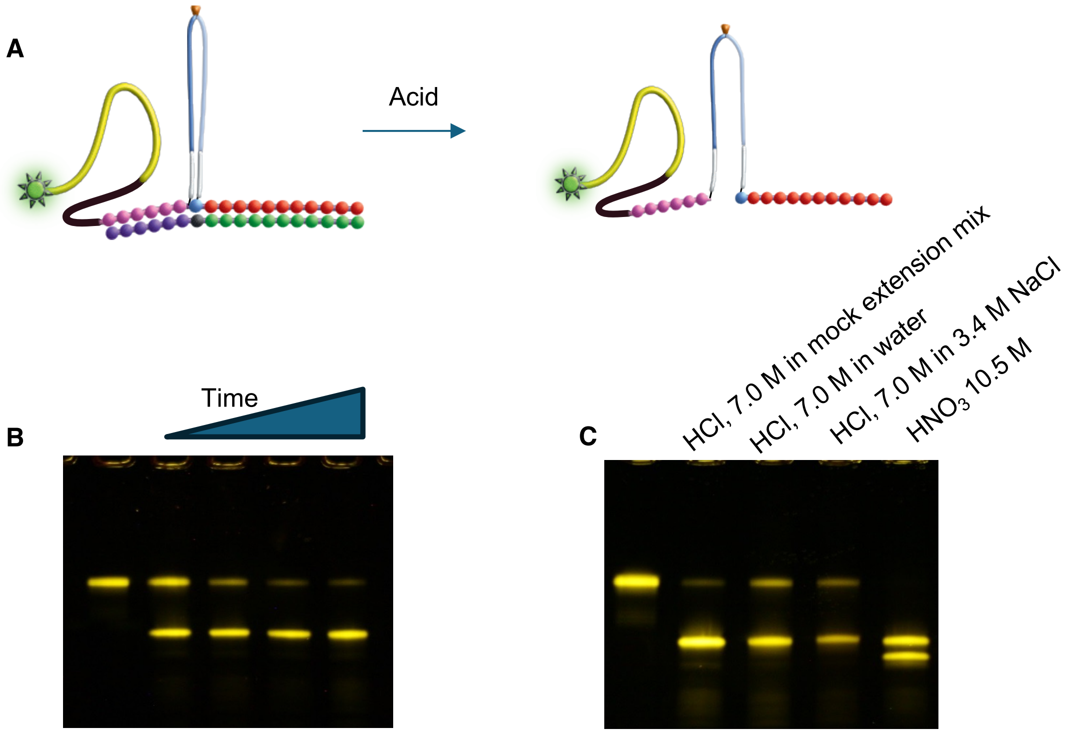


**Fig. S12. Cleavage of single XNTP incorporation. (A)** A fluorescent EO (100 pmol) was hybridized to a template with the sequence 3'-GAAAAAAAAAA. An extension mix including XCTP (500 pmol) was added and incubated 2.5 h, after which dTTP (1 nmol) was added and incubated an additional 30 min. The products were separated by electrophoresis on a 15% acrylamide gel. The product band was excised and extracted from the gel, resulting in primer-XC-TTTTTTTTTT (~35 pmol) which was used in the following cleavage assays. **(B)** Cleavage time course of single XCTP incorporation. Extension product (1 pmol) was diluted in a mock extension mix of 20 mM tris acetate (pH 8.5), 200 mM ammonium acetate, 20% DMSO, 2 mM $MnCl_2$, 5% PEG-6K with or without 7.0 M HCl (lane 1). The acid-treated samples in lanes 2-5 were incubated for 10, 20, 30, or 60 minutes at 37 C, then quenched with 52 µL of a buffer (450 mM sodium phosphate pH 7, 5 M NaOH, 0.17% Tween, 17 mM EDTA) to neutralize the acid. These samples were precipitated by adding 1 µL 25% Ficoll, 1 mL EtOH and spinning 18k rcf, cold, for 20 min. Pellets were washed with 200 µL EtOH and spun for another 5 minutes. Pellets were resuspended in gel loading buffer and analyzed by electrophoresis in 15% acrylamide gels. Breaking of the P–N bond results in a linearized molecule that migrates faster in the gel. Any depurination products would migrate even faster as a result of the lost 10T portion of the molecule. **(C)** Cleavage of the extension product in alternative buffers and acids. Extension product (1 pmol) was diluted in mock extension mix, water, or brine with 7.0 M HCl. An additional sample was diluted in water with 10.5 M nitric acid. All samples were incubated for 60 minutes at 37 C before quenching and precipitating. PAGE analysis shows that nitric acid caused additional cleavage beyond the ring opening.

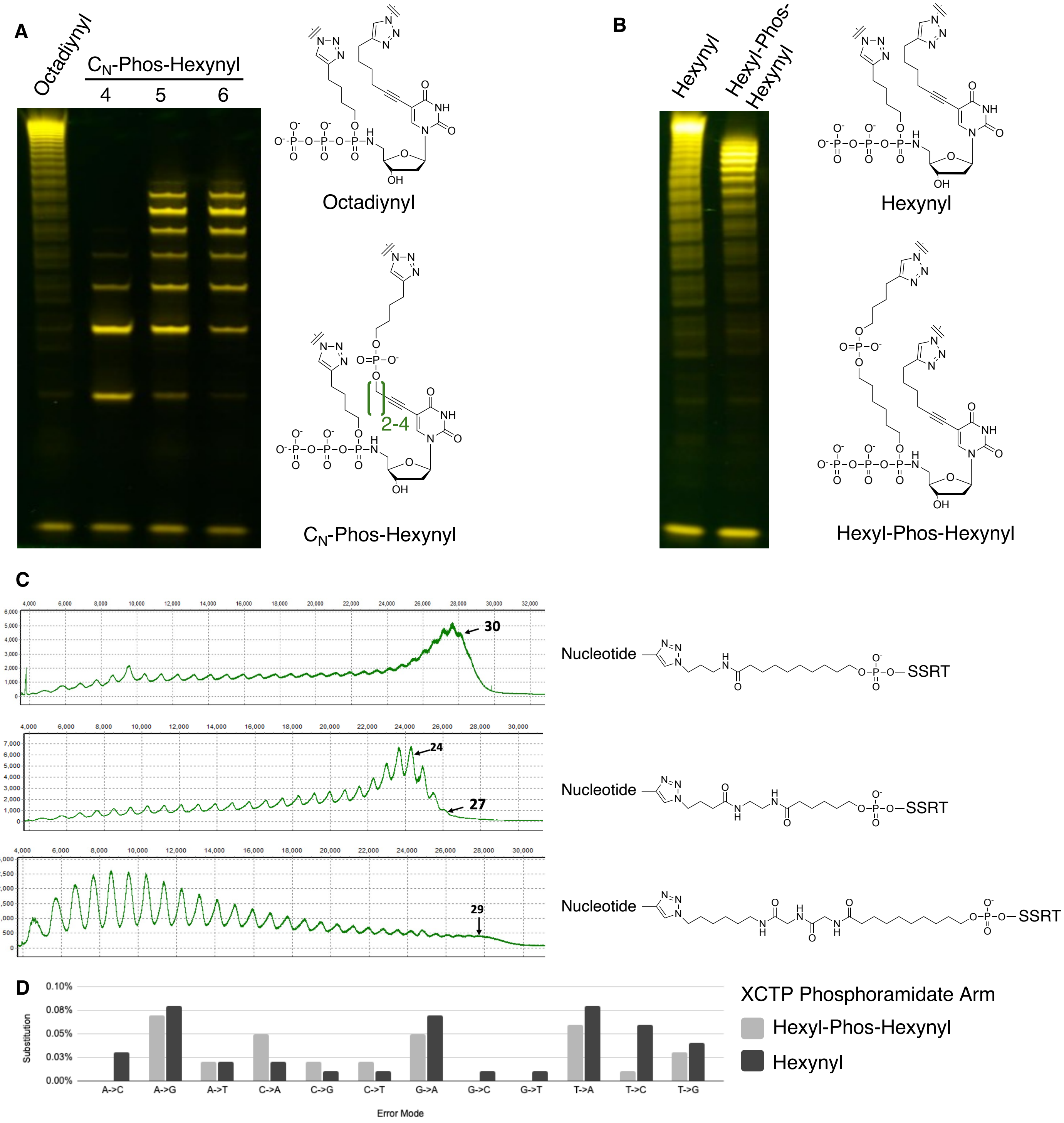


**Fig. S13**. **Optimization of linker arms for nucleotides and SSRTs. (A)** The alkyne linker group on the heterocycle was varied from octadiynyl to butynyl/pentynyl/hexynyl-phosphate-hexynyl. Cleaved extension products of XTTP incorporation using a polyA template were analyzed by TBE 4-12% PAGE. Linkers which brought the phosphate more proximal to the nucleobase inhibited extension compared to octadiynyl. **(B)** The alkyne linker group on the phosphoramidate was varied from hexynyl to hexyl-phosphate-hexynyl. Cleaved extension products of XTTP incorporation using a polyA template were analyzed by TBE 4-12% PAGE. Inclusion of the phosphate in the linker produced shorter Xp products. **(C)** The azide linker group of an early SSRT test structure was varied to include multiple amides and length differences. Cleaved extension products of XGTP incorporation using a polyC template were analyzed by capillary electrophoresis. Hydrophobic regions with minimal amides produced longer Xp products. The spacing may also play a role, as longer linkers could move the enhancer region further from the nucleotide. **(D)** Error profile of Xp synthesized from XCTP bearing either hexynyl or hexyl-phosphate-hexynyl phosphoramidate arms, clicked to the same alternative SSRT. The hexynyl arm leads to significant increases in T->C substitutions.

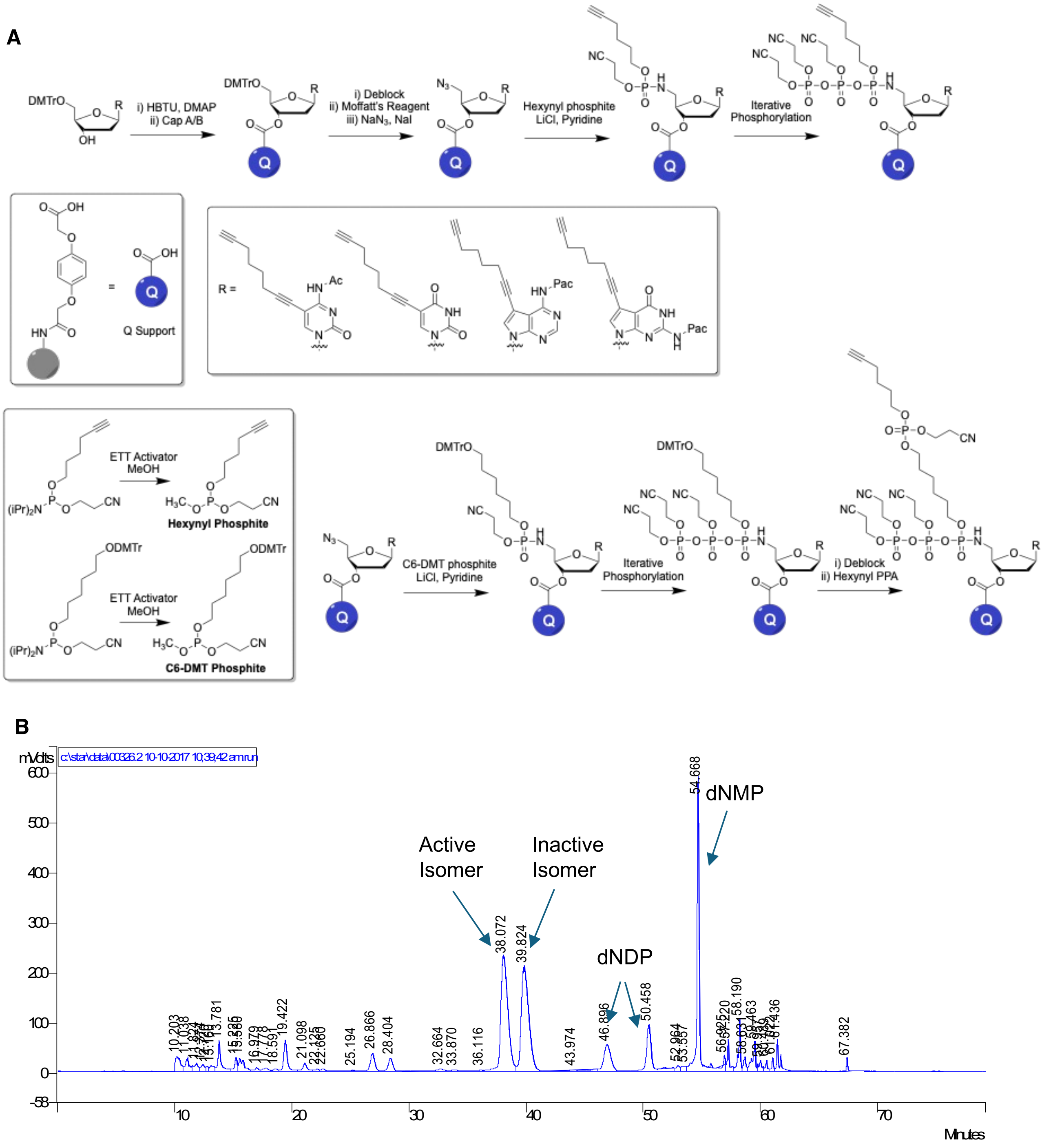


**Fig. S14. (A)** Synthesis of bis-alkyne nucleotide triphosphates. All nucleosides (upper right inset) were synthesized on support (upper left inset) through HBTU / DMAP coupling, then converted to the 5'-azide. Phosphites (lower inset) for hexynol or DMT-hexane diol were prepared from their respective phosphoramidites. The azides were reacted with the phosphites in the presence of LiCl to convert them to the protected phosphoramidate. Iterative phosphorylation with bis-cyanoethyl phosphoramidite built out the triphosphate. In the case of a compound arm, the DMT was removed and hexynyl phosphoramidite was added. **(B)** After the triphosphate was cleaved from support with ammonia and concentrated, it was purified by HPLC (Waters C18 column, 40-43% MeCN in 100 mM TEAA). Only the first eluting triphosphate peak is active for extension. Nucleotide diphosphate and monophosphate elute later.

**<u>EO Sequence</u>**

**RDDDDDDDDDD(Pc)LLLLLLLLLLLLLLLLLLLLLLLLLZZZZZZTCATAAGACGAACGGA**

**Cleavable support spacer** | **High charge leader** | **Concentrator** | **Acid stable primer**

**<u>Cleaved from support  EO – post Xp synthesis</u>**

**p-LLLLLLLLLLLLLLLLLLLLLLLLLZZZZZZTCATAAGACGAACGGA-Xp**

**Fig. S15. Extension Oligo PPA Sequence.** Synthesized from the primer on a universal support, 2'-methoxy C, A & G are used for acid stability. This is followed by the concentrator, composed of 6 x C12 PPA (Z) and the leader, composed of 25 x C2 PPA (L). The photocleavable linker (Pc) is then incorporated with 10 x PEG-6 (D) and a bromohex (R) that is converted to an azide. When UV light is used to cleave the Pc, the terminal C2 is capped by a phosphate (p).

**A**

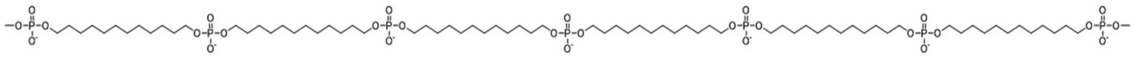

**B**

| Concentrator | EO sequence (5’ -> 3’) |
|---|---|
| Inline Z6 | LLLLLLLLLLLLLLLLLLLLLLLLLZZZZZZTCATAAGACGAACGGA |
| None | LLLLLLLLLLLLLLLLLLLLLLLLLTCATAAGACGAACGGA |

Leader, Concentrator, Priming site

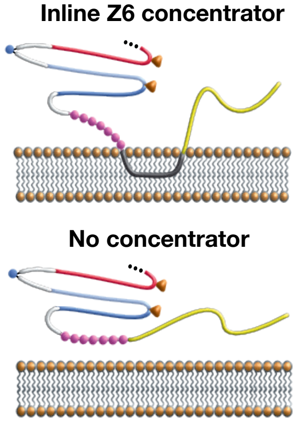


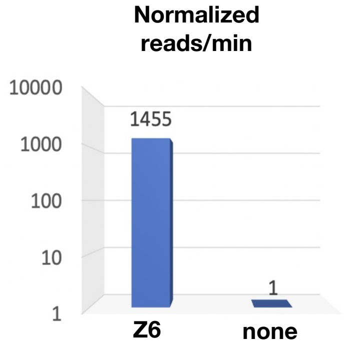


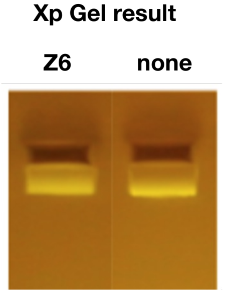


**C**

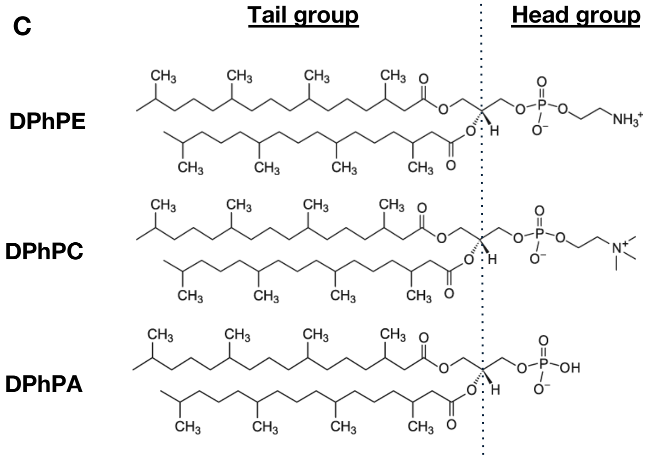


| XP sample ID | Lipid molecule | Bases/min |
|---|---|---|
| XP34526 | 100% DPhPE | 363 |
| | 100% DPhPC | 57 |
| XP34527 | 100% DPhPE | 378 |
| | 50% DPhPE : 50% DPhPC | 392 |
| XP34814 | 100% DPhPE | 566 |
| | 100% DPhPA | 61 |

**Fig. S16. Concentrator-lipid interaction. (A)** The EO contains a hydrophobic region called the concentrator which is positioned downstream of the leader. The concentrator is designed to associate with the lipid bilayer, which localizes the Xp leader near the lipid, enabling higher Xp capture frequency. **(B)** The concentrator is critical to high throughput sequencing. Synthesizing Xp without a concentrator present produces comparable yield by gel, but very few molecules can be captured by pore measurement. **(C)** This concentrating effect is in enhanced by using a DPhPE headgroup which improves throughput 6x compared to the DPhPC headgroup. Mixtures of lipids may provide improved balance of concentrator association and surface diffusion leading to higher Xp throughput.

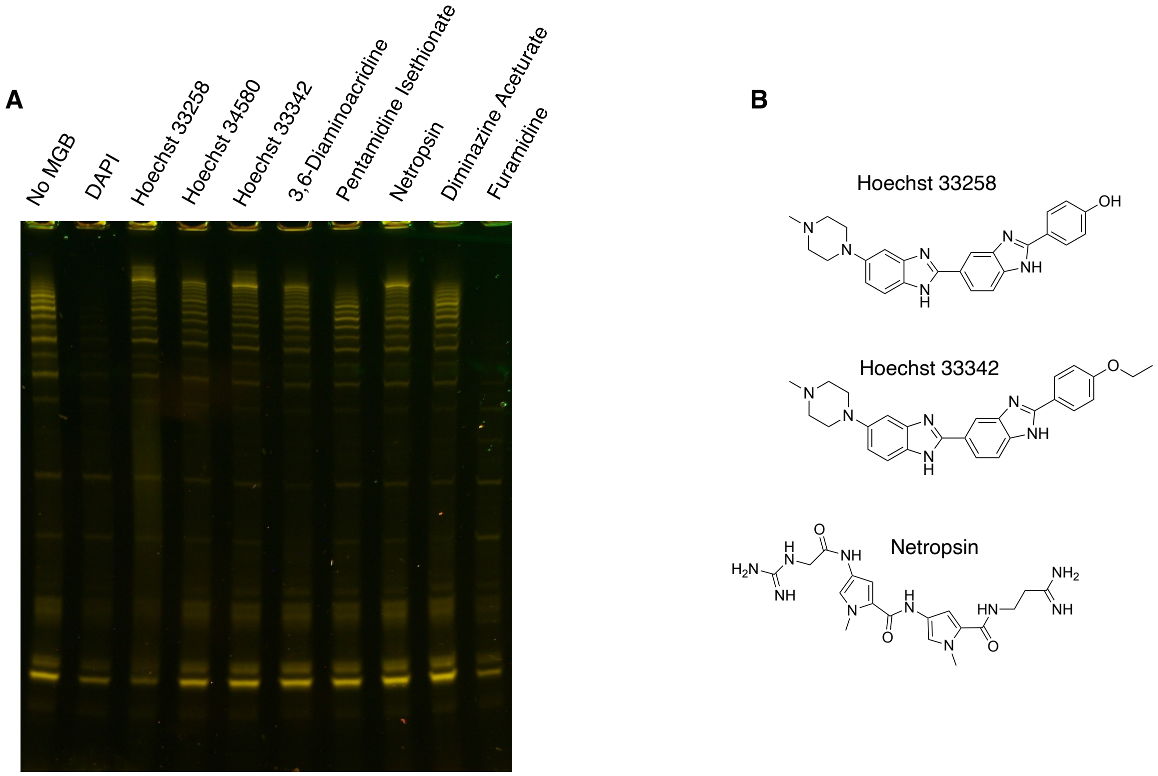


**Fig. S17. Minor groove binders in Xp synthesis. (A)** Analysis of Xp products synthesized from a 45-mer template. Fluorescent primer was extended in solution, and the unexpanded Xp products were visualized on a 4-12% acrylamide gel. Inclusion of 150 μM of various minor groove binding molecules enhances the length of the Xp products. **(B)** Structure of some of the best performing minor groove binders.

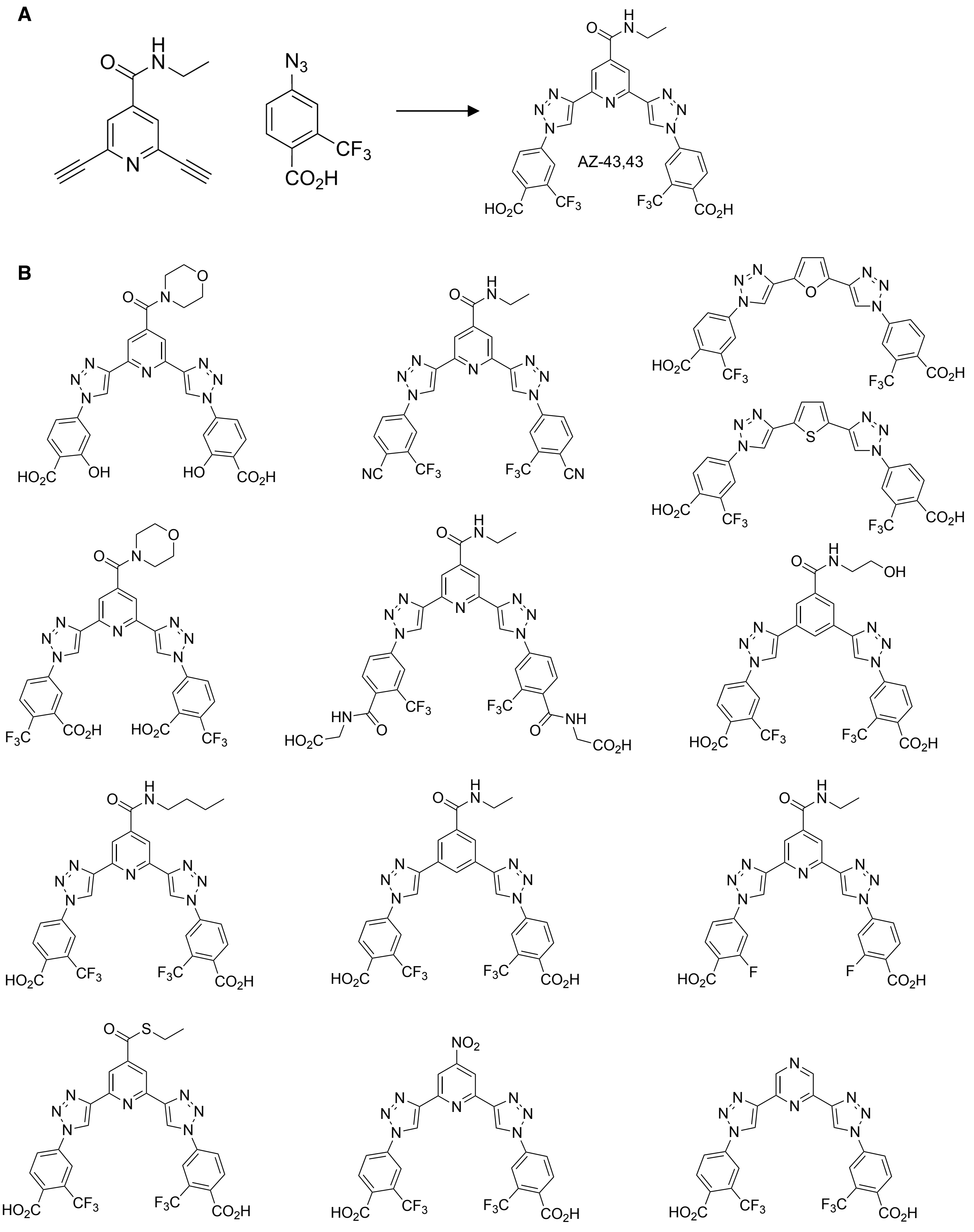


**Fig. S18: (A)** AZ (N-ethyl-2,6-diethynylisonicotinamide) core and 43 2-(trifluoromethyl)benzoic acid arms used in preparing the PEM AZ-43,43. **(B)** A selection of PEMs (amongst 264 synthesized) which showed a PEM effect. AZ-43,43 was selected due to a pronounced improvement in length when replicating a 222-mer template.

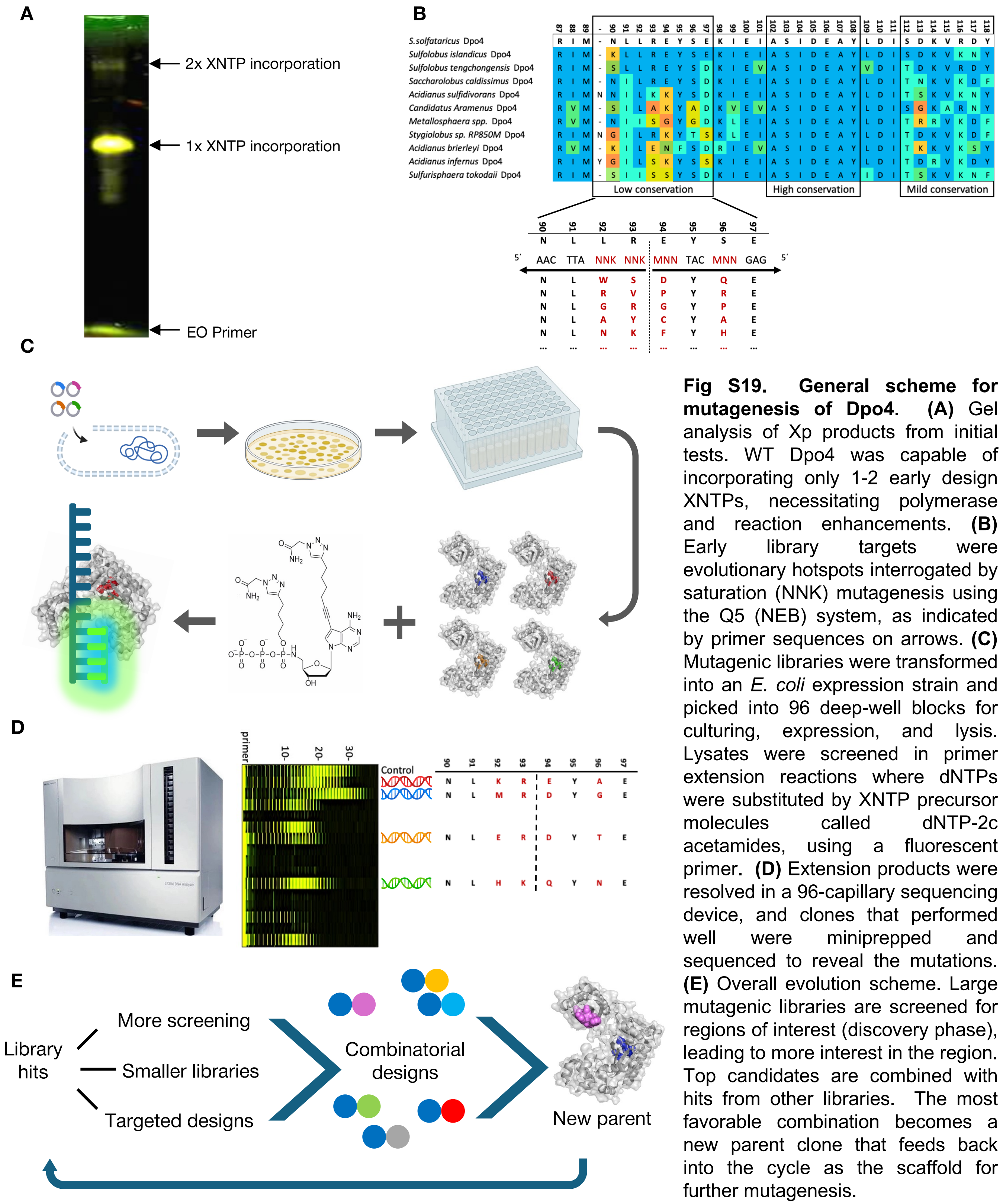


**Fig S19. General scheme for mutagenesis of Dpo4.** **(A)** Gel analysis of Xp products from initial tests. WT Dpo4 was capable of incorporating only 1-2 early design XNTPs, necessitating polymerase and reaction enhancements. **(B)** Early library targets were evolutionary hotspots interrogated by saturation (NNK) mutagenesis using the Q5 (NEB) system, as indicated by primer sequences on arrows. **(C)** Mutagenic libraries were transformed into an *E. coli* expression strain and picked into 96 deep-well blocks for culturing, expression, and lysis. Lysates were screened in primer extension reactions where dNTPs were substituted by XNTP precursor molecules called dNTP-2c acetamides, using a fluorescent primer. **(D)** Extension products were resolved in a 96-capillary sequencing device, and clones that performed well were miniprepped and sequenced to reveal the mutations. **(E)** Overall evolution scheme. Large mutagenic libraries are screened for regions of interest (discovery phase), leading to more interest in the region. Top candidates are combined with hits from other libraries. The most favorable combination becomes a new parent clone that feeds back into the cycle as the scaffold for further mutagenesis.

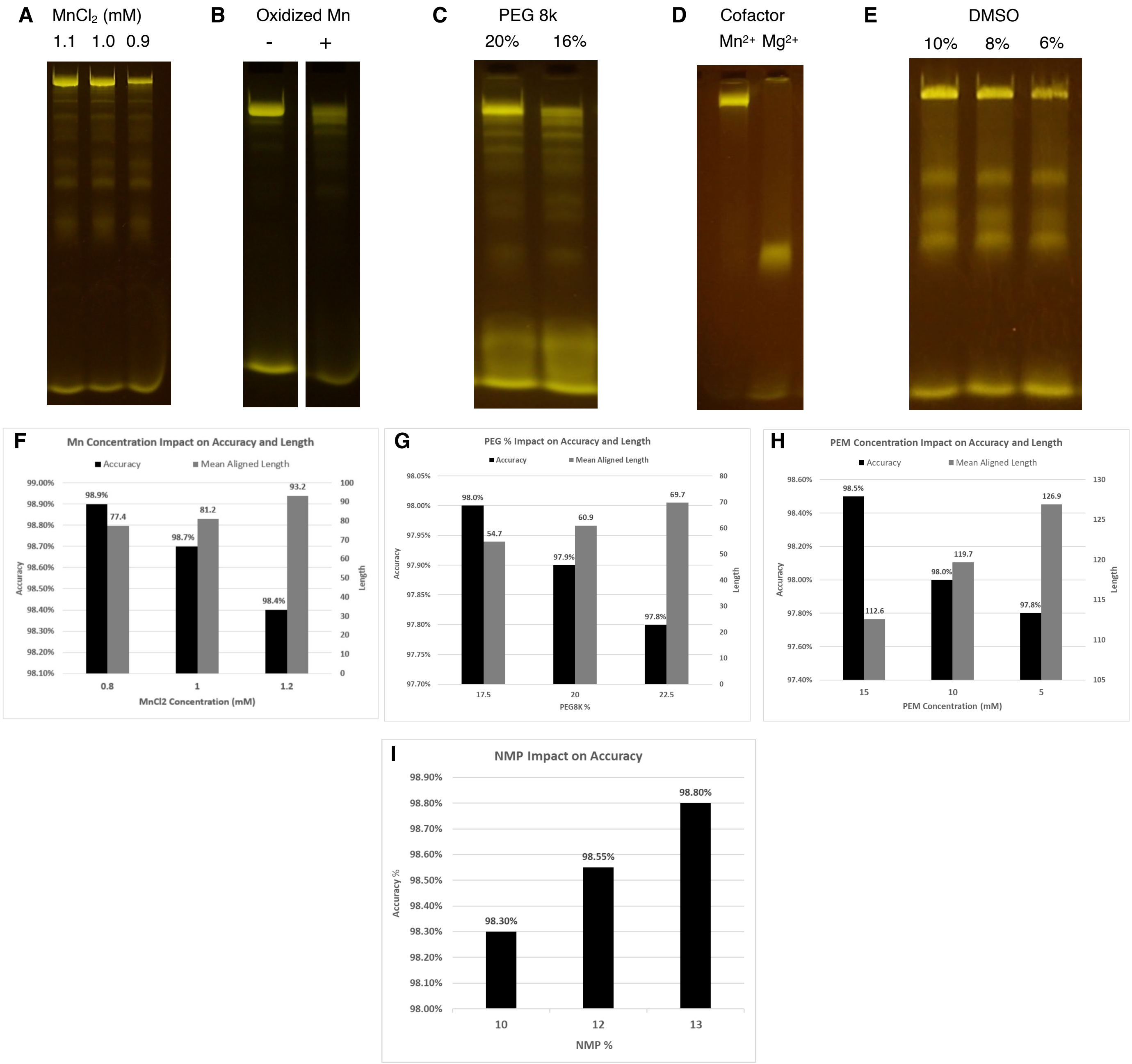


**Fig. S20. Synthesis reaction formulation. (A)** Analysis of Xp products synthesized from a 222-mer template. Uncleaved Xp products were extended using a fluorescent primer, quenched with loading buffer and electrophoresed through a 2.5% agarose gel. Depleting the reaction of $MnCl_2$ results in a slower reaction and less full-length product. **(B)** $MnCl_2$ was oxidized by exposure to air, which turned it hazy brown. Less full-length product was observed in these samples, consistent with lower effective $Mn^{2+}$ concentration. **(C)** Xp synthesized from 222-mer template with variable amounts of PEG 8k. When the concentration of PEG 8k was reduced from 20% to 16%, the yield of full-length product decreased. **(D)** When $Mn^{2+}$ was replaced by $Mg^{2+}$, the Xp products were very short. **(E)** Analysis of uncleaved Xp products from 222-mer template using a fluorescent primer. With 15 mM PEM in the Xp synthesis, lowering the DMSO in the reaction lead to shorter products. **(F)** Sequenced Xp synthesized from a 100-mer template using different concentrations of $MnCl_2$. Reducing the $Mn^{2+}$ concentration in the extension reaction results in shorter average product size, but higher accuracy. **(G)** Sequenced Xp from HIV2 100-mer with variable PEG. Increased PEG led to longer products, but also lower accuracy. **(H)** Sequenced Xp from Strep 222-mer with variable PEM. Increased PEM led to shorter products, but also higher accuracy, potentially due to $Mn^{2+}$ sequestration. **(I)** Sequenced Xp from 4-base repeat 60-mer with variable NMP. Accuracy increased with increased solvent.

**A**

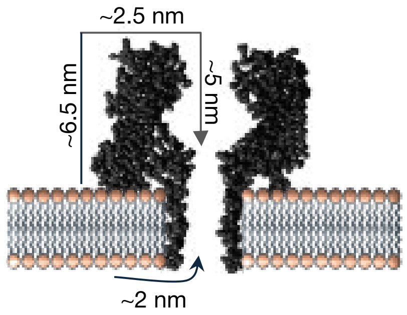


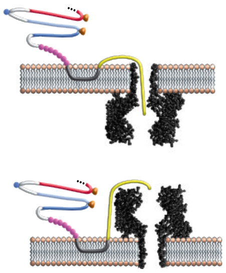

| Leader | Approximate Length | Barrel → vestibule events/min | Vestibule → barrel events/min |
|---|---|---|---|
| L5 | 2 nm | -- | -- |
| L25 | 10 nm | 115 | 0 |
| L50 | 20 nm | -- | -- |
| L100 | 40 nm | 131 | 6 |

**B**

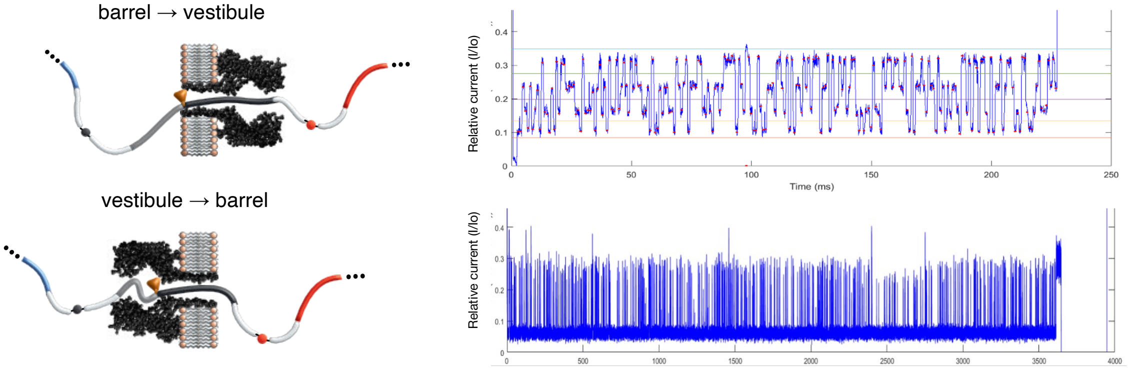


**C**

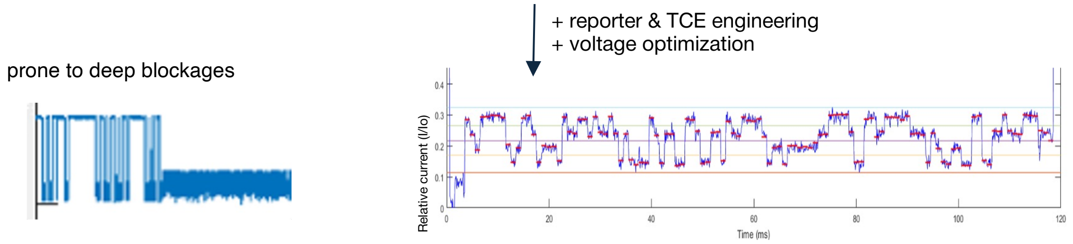


**D**

| | Bases / minute | Accuracy | Deletions | Insertions | Substitutions |
|---|---|---|---|---|---|
| Barrel → vestibule | 16638 | 98.2% | 0.4% | 0.6% | 0.8% |
| Vestibule → barrel | 2364 | 85.5% | 4.23% | 8.1% | 2.30% |

**Fig. S21. Nanopore orientation. (A)** Traditionally, nanopore sequencing using αHL is done in a vestibule-to-barrel orientation. SBX sequencing uses the reverse orientation of barrel-to-vestibule, which leads to a higher capture rate. Sequencing in the barrel-to-vestibule orientation also prevents recapturing and subsequent resequencing of translocated Xp molecules from the *trans* side when reversing the current. Given the lipid-concentrator relationship, the length of the leader plays a key role in capture efficiency for each orientation. **(B)** Due to the asymmetry of the hemolysin pore, the directionality of the pore will alter the current characteristics and Xp interactions. In the vestibule-to-barrel orientation, the vestibule cavity leads to deep blockages within the pore. **(C)** SSRTs and nanopore run conditions required extensive reengineering to accommodate vestibule-to-barrel sequencing, which still produced suboptimal results and continued to be prone to deep blockages. **(D)** Sequencing results show substantial advantage of the barrel-to-vestibule orientation in both accuracy and throughput.

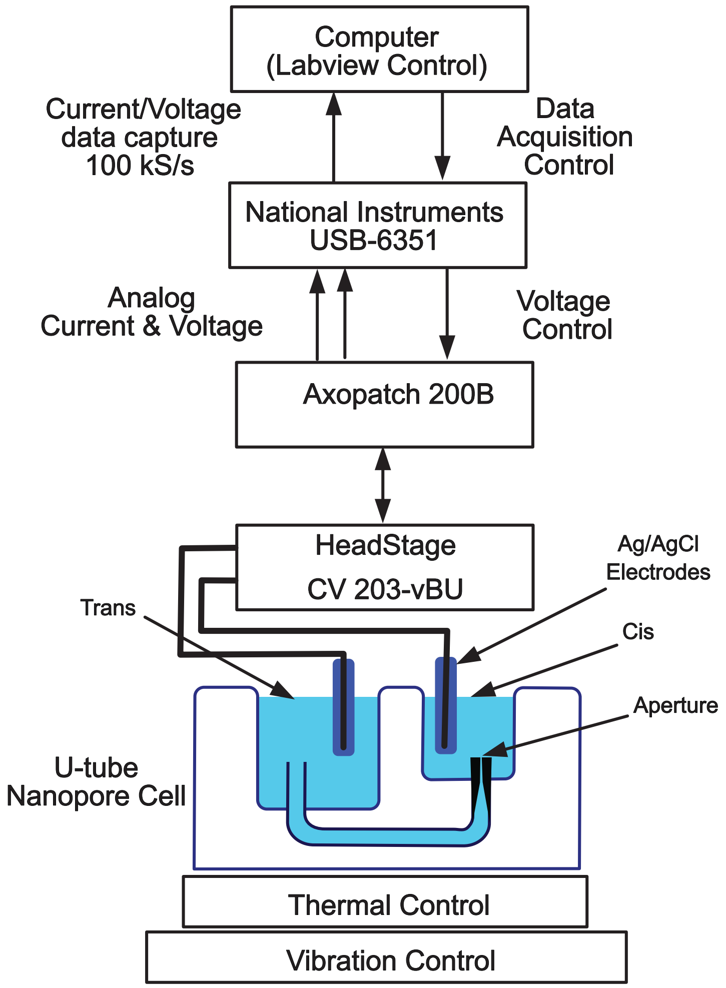


**Fig. S22. Schematic of the nanopore instrumentation.** This instrument is based upon that of Marziali (*8*). Adapted PTFE U-tube cell had a 13 μm aperture for the lipid membrane support and volumes for the *cis* and *trans* reservoirs were 80 and 300 μL respectively. A Labview script controlled timing and levels of voltages applied by the Axopatch 200B amplifier to the nanopore. The amplifier returned analog outputs proportional to the nanopore ion current and applied voltage which were synchronously digitized, captured and stored.

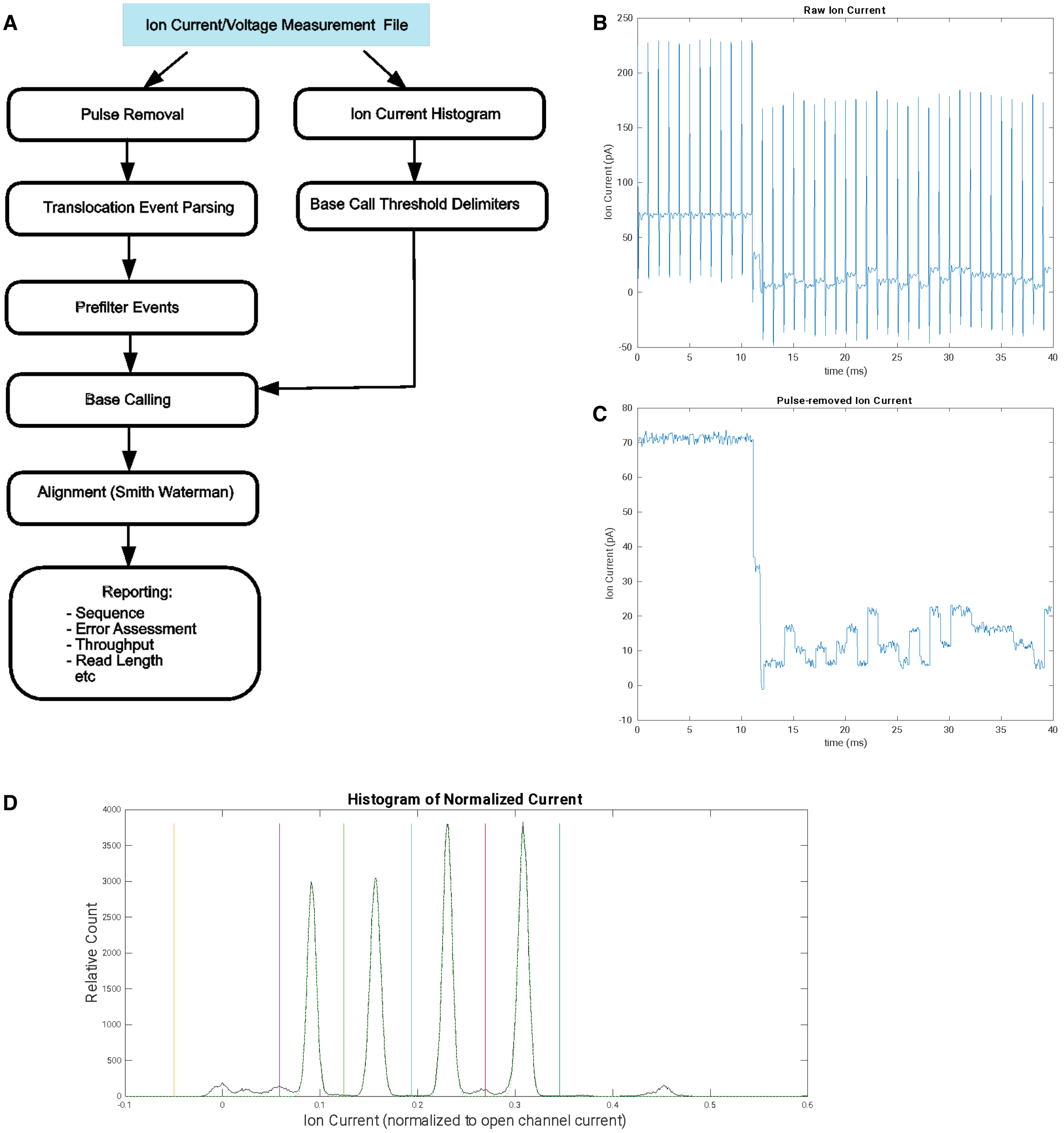


**Fig. S23. Ion current measurement. (A)** The ion current and applied voltage data for the run were recorded in a raw data file for post-processing with 16-bit sampling at 100 kS/s. A MATLAB script processed the raw data file following this data path. **(B)** Example of a raw current trace showing pulses that occur in response to the voltage pulses used to step the Xp translocation. **(C)** Initial post-processing removes pulses from raw data as the processed example shows. **(D)** Histogram plot of Xp data samples after pulse removal normalized to the 71 pA open channel ion current. Demarcation lines were created between (and along edges) of the four primary peaks. These determined the ranges for basecalling.

### Additional Ion Current Traces

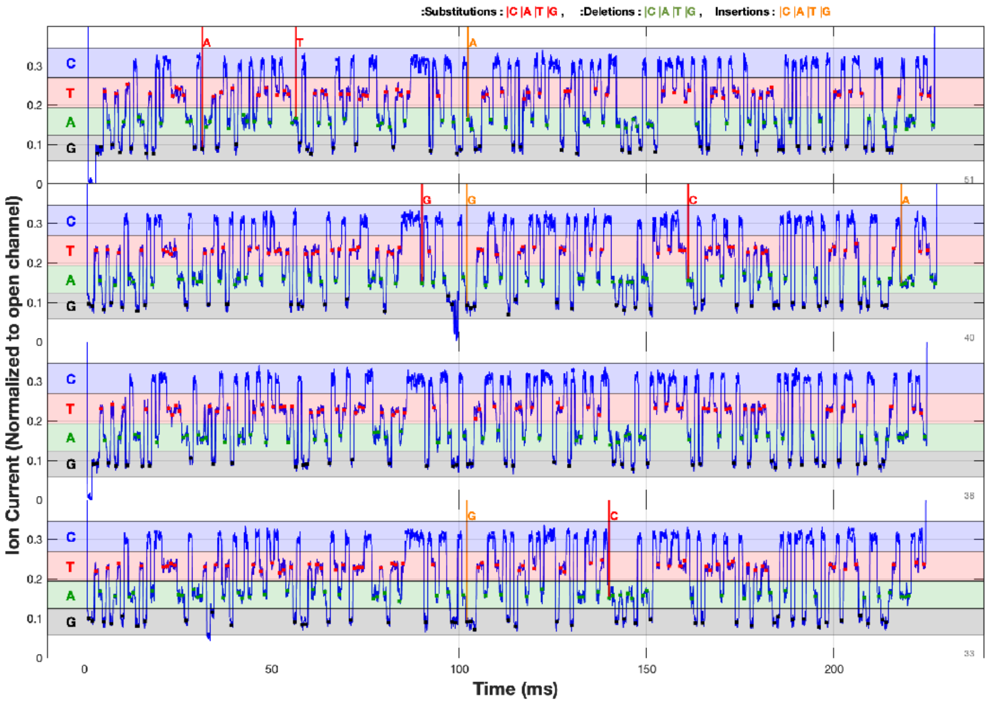


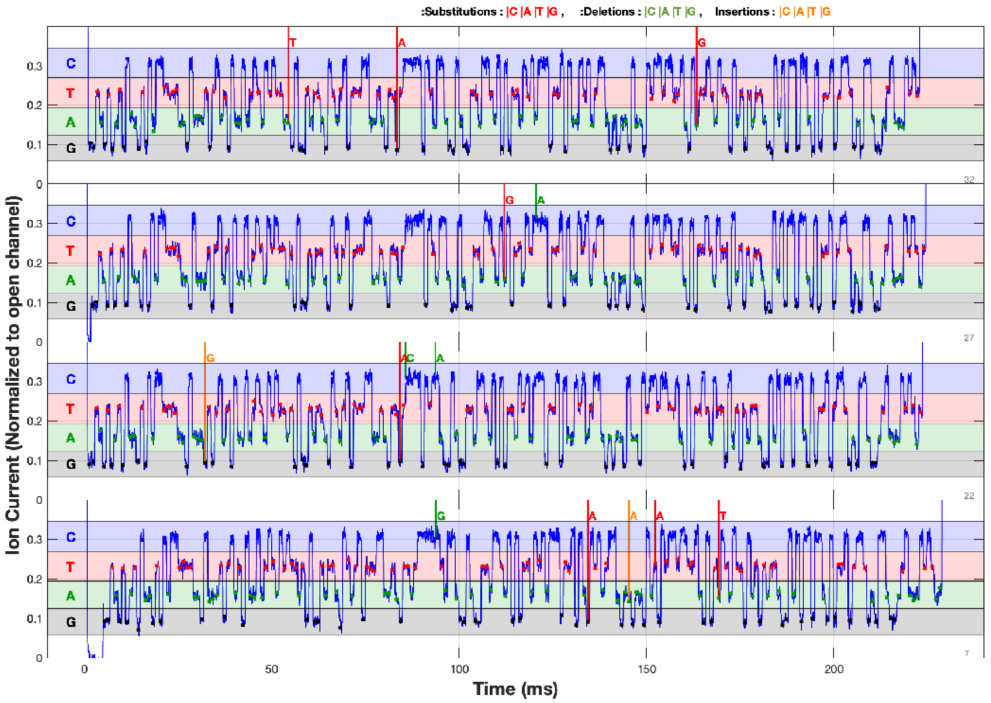

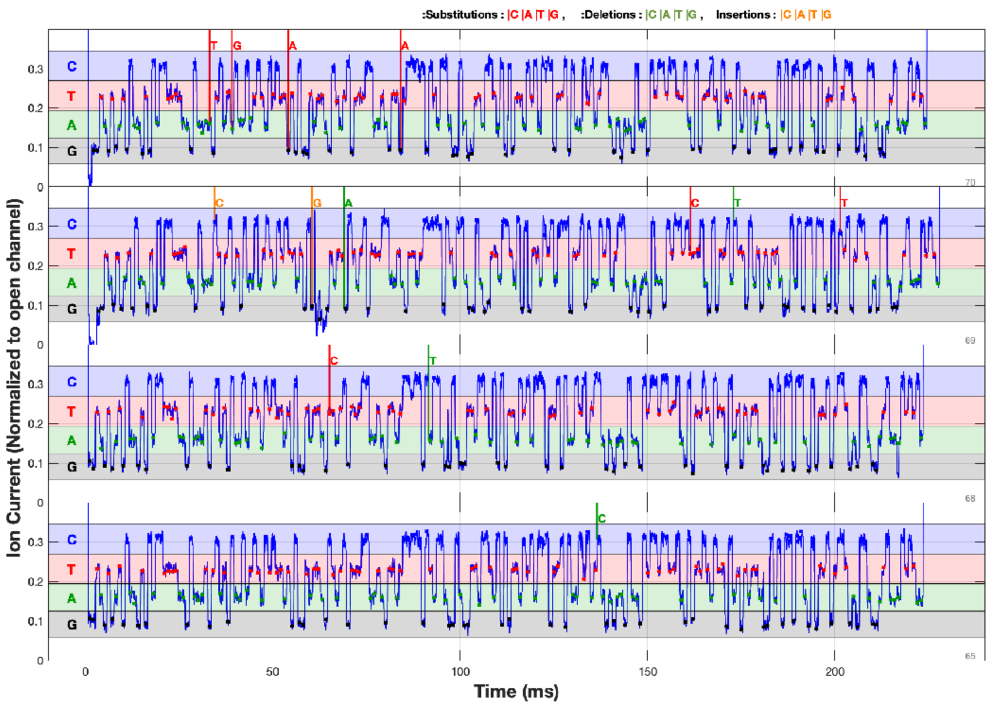
:Substitutions : |C |A |T |G , :Deletions : |C |A |T |G , Insertions : |C |A |T |G
C
T
A
G
Ion Current (Normalized to open channel)
Time (ms)
0
50
100
150
200

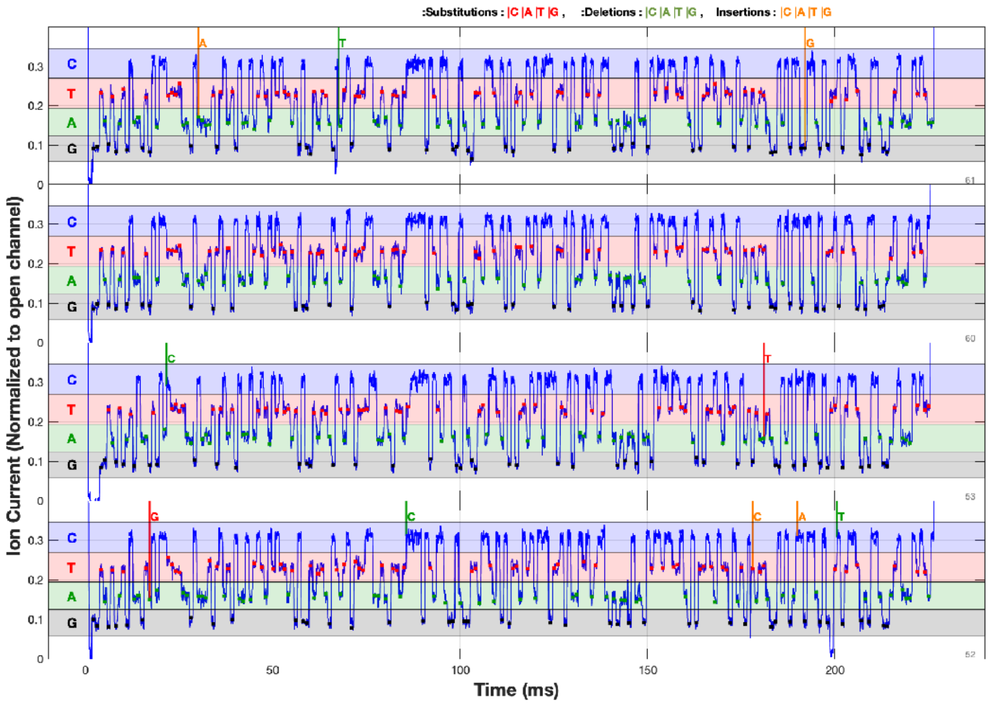
:Substitutions : |C |A |T |G , :Deletions : |C |A |T |G , Insertions : |C |A |T |G
C
T
A
G
Ion Current (Normalized to open channel)
Time (ms)
0
50
100
150
200

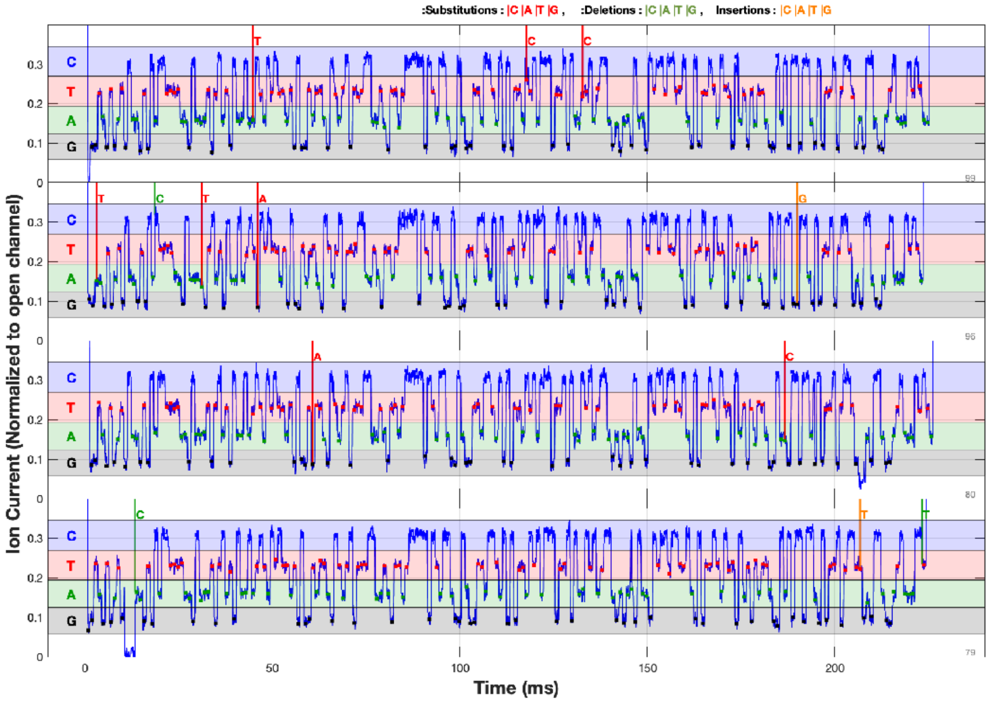


**Fig. S24. Additional full-length Xp ion current traces.** Errors are indicated above each trace as called by the Smith-Waterman algorithm. Substitutions, deletions and insertions are indicated by red, green and orange base letters depicting the substituted, missing or inserted base respectively.

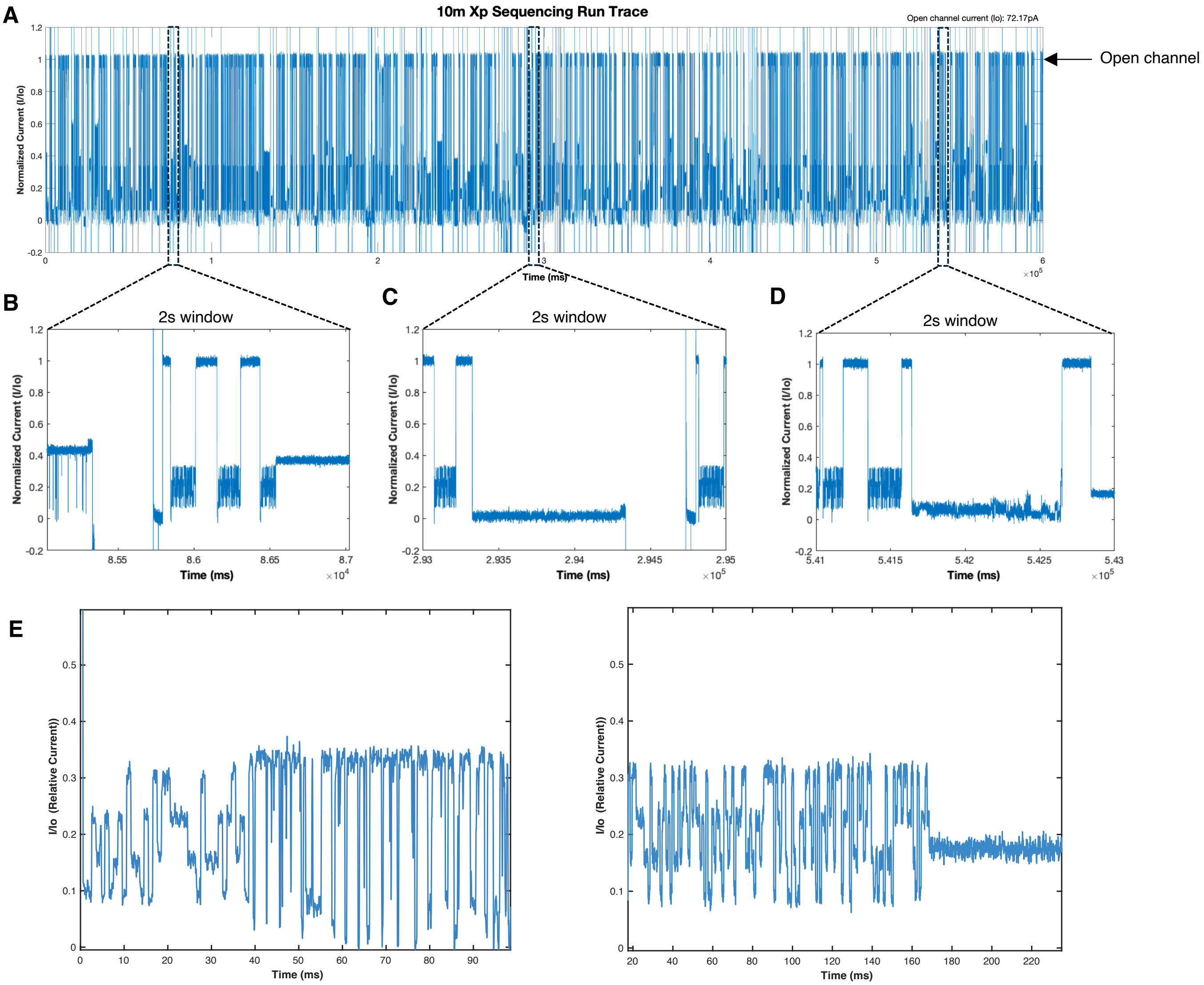


**Fig. S25**. **Various examples of nanopore blockages. (A)** Blockages occur randomly and often throughout a measurement run and reduce the potential throughput of SBX sequencing. A variety of blockage signals are seen ranging from high **(B)** to low **(C)** ion current levels. Data is assessed in real time and if blockage criteria are met the Xp is ejected automatically with a 400 ms/250 mV voltage pulse in the reverse polarity before reverting to measurement conditions again. **(D)** Certain blockage types can translocate fully through the pore without voltage intervention but are slow to translocate and may yield some or no sequence data. **(E)** Additional examples of blockage types.

| Clone # | Genotype | Advantage |
|---|---|---|
| C0001 | Dpo4_I341L/Δ343-352 (removed PIP box) | Y |
| C0005 | C0001_M76W/K78N/E79L/Q82W/Q83S/S86D | L |
| C0050 | C0005_A42V/Q83G/S86E/T141S/F150L/I153F/A155V/D156Y/I217V/I226F/V289W/T290K/E291S/D292Y/L293W/D326E | L |
| C0157 | C0050_**S141T/L150F**/K152G/**F153I/V155A**/Y156W/**V217I/F226I/E326D** | Y |
| C0345 | C0157_K56Y | L, A |
| C0416 | C0345_A155M/I248T | L |
| C0534 | C0416_G187P/N188Y/T190Y | L |
| C1067 | C0534_G152L/I153T/M155G/W156R | L, Y |
| C1816 | C1067_D294N/I295S/V296Q/S297Y/G299W/R300S/T301W/K317Q/K321Q/E324K/E325K/E327K | L |
| C3405 | C1816_N78D | A |
| C3488 | C3405_L152A/T153V | L |
| C4552 | C3488_L184Q/W189F/**Y190T** | L, A |
| C4760 | C4552_E63R | L |
| C5086 | C4760_S272C | T |
| C5173 | C5086_**Q317K** | A |
| C5213 | C5173_A57S | A |
| C5250 | C5213_I59M | Y |
| C5275 | C5250_**V42A** | A |
| C5275 | C0001_K56Y/A57S/I59M/E63R/M76W/K78D/E79L/Q82W/Q83G/S86E/K152A/I153V/A155G/D156R/P184Q/G187P/N188Y/I189F/I248T/S272C/V289W/T290K/E291S/D292Y/L293W/D294N/I295S/V296Q/S297Y/G299W/R300S/T301W/K321Q/E324K/E325K/E327K | A |

**Table S1. Lineage of SBX polymerase clones from wild type to Xp synthase C5275.** The relevant mutation(s) from parent is shown for each clone. The C5275 entries show both the parent mutation and the final suite of mutations on that clone, which summarizes the changes in the table. A=accuracy, L=extension length, Y=soluble protein yield, T=thermostability. Bold indicates a reversion to the wild type sequence from a previously mutated position. Data for the observed length and accuracy changes are not given for all changes shown here, because the methods and materials used to determine these values changed dramatically over the 6 years of polymerase development reflected in this work.
